# Parameter Inference for an Astrocyte Model using Machine Learning Approaches

**DOI:** 10.1101/2023.05.16.540982

**Authors:** Lea Fritschi, Kerstin Lenk

## Abstract

Astrocytes are the largest subset of glial cells and perform structural, metabolic, and regulatory functions. They are directly involved in the communication at neuronal synapses and the maintenance of brain homeostasis. Several disorders, such as Alzheimer’s, epilepsy, and schizophrenia, have been associated with astrocyte dysfunction. Computational models on various spatial levels have been proposed to aid in the understanding and research of astrocytes. The difficulty of computational astrocyte models is to fastly and precisely infer parameters. Physics informed neural networks (PINNs) use the underlying physics to infer parameters and, if necessary, dynamics that can not be observed. We have applied PINNs to estimate parameters for a computational model of an astrocytic compartment. The addition of two techniques helped with the gradient pathologies of the PINNS, the dynamic weighting of various loss components and the addition of Transformers. To overcome the issue that the neural network only learned the time dependence but did not know about eventual changes of the input stimulation to the astrocyte model, we followed an adaptation of PINNs from control theory (PINCs). In the end, we were able to infer parameters from artificial, noisy data, with stable results for the computational astrocyte model.

## 1 Introduction

Together with the well-studied neurons, glial cells make up the nervous system. Astrocytes are the largest group of glial cells and provide structural support, perform diverse metabolic and regulatory functions, and are responsible for the regulation of synaptic transmission. They connect to neurons at synaptic clefts [Araque et al., 1999] and absorb glutamate and other neurotransmitters released by firing neurons. As a reaction to the glutamate, the intracellular calcium (Ca^2+^) concentration of astrocytes rises, causing Ca^2+^ transients that can spread over multiple astrocytes, and causes the release of ions and transmitter molecules that affect the neurons. Malfunctions in astrocytes have been connected to multiple diseases such as Alzheimer’s, Huntington’s [Siracusa et al., 2019], schizophrenia [Notter, 2021], and epilepsy [Verhoog et al., 2020]. Research has also shown that astrocytes play an important role in the acquisition of fear memory, offering new ways to potentially treat anxiety-related disorders [Liao et al., 2017, Li et al., 2020].

To this day, astrocytes remain difficult to study and observe. Therefore, many of their pathways remain unknown. To aid the general understanding and research of astrocytes, several computational models have been proposed. The types of models range from network models, that attempt to simulate whole neuron and astrocyte networks [Lenk et al., 2020], over single cell models, used to study Ca^2+^ wave propagation and neuron interaction [Larter and Craig, 2005, Nadkarni and Jung, 2007, De Pitta and Brunel, 2016], to single compartment models [Denizot et al., 2019, Oschmann et al., 2017] focusing on only a small part of an astrocyte.

However, these models are often incomplete and rely on parameters that often are not available. Respective measurements are too expensive or not possible with current technology. Thus, one major challenge of computational astrocyte models, and computational models in general, is the fast and accurate inference of parameters. Well-known methods for parameter inference include least squares fitting [Liu et al., 2012, Dattner et al., 2019], genetic algorithms [Mitchell, 1998], Bayesian inference methods such as Markov Chain Monte Carlo (MCMC) [Valderrama-Bahamóndez and Fröhlich, 2019] or, though more often used in robotics, Kalman filters [Lillacci and Khammash, 2010]. More recently, Yazdani et al. [2020] proposed to use physics informed neural networks (PINNs) to infer parameters. In contrast to the more traditional methods, PINNs make use of the underlying physics to infer parameters and, if necessary, dynamics that can not be observed.

In this study, we focus on the computational model of an astrocytic compartment developed by Oschmann et al. [2017]. Using the parameter inference algorithm originally developed by Yazdani et al. [2020], we demonstrate different problems with the algorithm and propose solutions that aim to stabilize the algorithm. Using the stabilized inference algorithm, we then go on and infer the parameters for different currents underlying the molecular dynamics of the astrocytic compartment model.

## 2 Background

In this section, we introduce the two main topics of this study: astrocytes and parameter inference in the context of machine learning.

### 2.1 Biology of Astrocytes

Astrocytes are a type of glial cell usually found in the brain and spinal cord. For many years, it was assumed that astrocytes only serve structural, metabolic, and regulatory functions. However, in the last 30 years, this view has been challenged by a multitude of research suggesting that astrocytes are also involved in the control of synaptic transmission [Vesce et al., 1999, Araque et al., 1999, Haydon and Carmignoto, 2006, Nedergaard and Verkhratsky, 2012, Araque et al., 2014].

According to research, the number of astrocytes and their exact morphology can vary widely between different species, brain regions, and brain layers [Zhou et al., 2019]. For example, it has been shown that astrocytes in the human neocortex are 2.6 times larger in diameter and exhibit up to 10 times as many primary processes as the astrocytes of rodents [Oberheim et al., 2009]. Experiments performed by Buosi et al. [2018] showed distinct astrocytic gene expressions between different brain regions. Furthermore, Lanjakornsiripan et al. [2018] described differences in cell orientation, territorial volume, and arborization between different layers in the somatosensory cortex of mice.

While astrocytes are very heterogeneous in form and function [Verkhratsky and Nedergaard, 2018], they can generally be described as star-formed and highly branched cells. Each cell consists of a soma with several outgoing branches that split into smaller branchlets and then into distal processes. Several intracellular Ca^2+^ storages (endoplasmatic reticulum, ER) and mitochondria are placed along astrocytic processes. The volume of ER decreases along the astrocytic process [Patrushev et al., 2013]. Astrocytic distal processes can enclose neuronal synapses, thereby forming a so-called tripartite synapse [Araque et al., 1999] consisting of pre- and postsynaptic neurons as well as of an astrocyte. Furthermore, neighboring astrocytes communicate with each other through gap junctions, thereby forming a separate network.

In contrast to neurons, astrocytes are not electrically excitable [Verkhratsky and Nedergaard, 2018]. Instead, the main signal of astrocytes is considered to be Ca^2+^ transients. Ca^2+^ transients can either involve the whole astrocytic cell body as well as neighboring astrocytes or different proportions of an astrocytic process [Di Castro et al., 2011, Srinivasan et al., 2015]. The propagation of Ca^2+^ waves through gap junctions is assumed to be mediated either intracellular, through the direct diffusion of IP3 [Giaume and Venance, 1998], or by an extracellular diffusion of ATP [Guthrie et al., 1999, Fujii et al., 2017]. As a reaction to increased intracellular Ca^2+^ levels, astrocytes release gliotransmitters, such as glutamate, D-Serine, adenosine triphosphate (ATP), and gamma-Aminobutyric acid (GABA), that modulate the synaptic properties of enclosed neurons [Serrano et al., 2006, Henneberger et al., 2010, Sahlender et al., 2014, Harada et al., 2015].

In 2011, Di Castro et al. [2011] used high-resolution two-photon laser scanning microscopy (2PLSM) to observe endogenous Ca^2+^ activity along an astrocytic process. By subdividing the astrocytic process into smaller subregions (compartments) and recording their respective Ca^2+^ activity, they were able to observe two different categories of Ca^2+^ transients. Focal transients, mostly occurring at random and being confined to single compartments, and extended transients, cause larger, compartment-overlapping Ca^2+^ elevations. Furthermore, the authors noticed that the occurrence of transients was directly influenced by blocking or potentiating action potentials and transmitter release, proofing that Ca^2+^ transients might in part be triggered by neuronal activity.

The mechanism underlying Ca^2+^ dynamics can be separated into two different pathways [Wallach et al., 2014, Helen et al., 1992], both being attributed to the uptake of glutamate by astrocytes. On the one hand, the released glutamate binds to respective metabotropic receptors (mGluR) in the astrocytic plasma membrane, causing a release of inositol 1,4,5-trisphosphate (*IP*_*3*_) into the cytosol. Larger concentrations of *IP*_*3*_ increase the probability of open *IP*_*3R*_ channels between the astrocytic ER and intracellular space, leading to an increase in intracellular Ca^2+^ levels [Bezprozvanny et al., 1991]. The increased intracellular Ca^2+^ concentration elevates the probability of open *IP*_*3R*_ channels further, leading to a Ca^2+^-induced Ca^2+^ release (CICR) mechanism. Ca^2+^ is transported back into the ER using ATP via the sarco endoplasmic reticulum *Ca*^2+^-ATPase (SERCA) pump). On the other hand, the released glutamate activates glutamate transporters (GluT). In exchange for one potassium (K^+^) ion, GluT one glutamate-, one hydrogen, and three sodium (Na^+^) ions into the intracellular space. The changes in Na^+^ and K^+^ level influence two other transport mechanisms, namely the Na^+^-Ca^2+^ exchanger (NCX) and the Na^+^-K^+^ adenosine triphosphatase (NKA). Depending on the intracellular Na^+^ levels, NCX transports three Na^+^ ions out/into the cell and one Ca^2+^ ion into/out of the cell, respectively. Similarly, NKA exchanges three intracellular Na^+^ ions for two extracellular K^+^ ions. Additionally, depending on the current membrane voltage and the Nernst potentials of Na^+^ and K^+^ respectively, Na^+^ and K^+^ ions leak out of the cell.

### 2.2 Computational Models of Astrocytes

So far, a multitude of computational astrocyte models have been developed. Generally, different models can be categorized into network models, single cell models, or single compartment models [Oschmann et al., 2018, González et al., 2020].

#### 2.2.1 General Overview

Many astrocyte models focusing on the interaction between astrocytes have been published. For example, Goldberg et al. [2010] studied Ca^2+^ signaling through gap junctions inside a small astrocyte chain. Assuming that Ca^2+^ waves are propagated through the exchange of *IP*_*3*_ molecules through gap junctions, they found that long-distance Ca^2+^ waves require the astrocyte network to be sparsely connected, to have a non-linear coupling function and a threshold, that, if not reached, causes the wave to dissipate. Similar observations were made in a later paper by Lallouette et al. [2014] that includes more complex networks. An astrocytic network model including both, the propagation of waves using *IP*_*3*_ and ATP, was proposed by Kang and Othmer [2009]. In their paper, they showed that the *IP*_*3*_ and ATP pathways can be distinguished from each other by looking at the delay between cells. Since astrocyte morphology and spatiotemporal patterns were found to play an important role in astrocyte function, Verisokin et al. [2021] proposed an algorithm to create realistic, data-driven astrocyte 2D morphologies. Other network models include both astrocytes and neurons. Using a simple neuron-astrocyte architecture based on anatomical observations made in the hippocampal area, Amiri et al. [2013] showed the influence of astrocytes on neuron synchronicity. Lenk et al. [2020] presented a discrete computational astrocyte-neuron model consisting of a neuronal network, an astrocyte network, and joint tripartite synapses. They used the model to study the effects of astrocytes on neuronal spike- and burst rate.

Several models simulate the interaction between neurons and astrocytes at a tripartite synapse. For instance, Nadkarni and Jung [2007] simulated a tripartite synapse of an excitatory pyramidal neuron. Their model assumes that astrocytes release glutamate in response to synaptic activity, thereby regulating Ca^2+^ at the presynaptic terminal. The effects of glutamatergic gliotransmission were further studied using a computational model by De Pitta and Brunel [2016]. In that model, the authors assumed that the release of gliotransmitters by the astrocyte is Ca^2+^-dependent and showed that gliotransmitter release is able to swap the synaptic plasticity between depressing and potentiating effects. Oyehaug et al. [2011] studied the effect of high K^+^ accumulation during neuronal excitation using a tripartite synapse model with detailed glial dynamics. They found that the presence and uptake of K^+^ by astrocytes are necessary to keep neurons from deactivating due to membrane depolarization.

Most models introduced so far release gliotransmitters that act on connected neurons, but not on the releasing astrocyte itself. An exception to this is the single-cell model developed by Larter and Craig [2005]. In this model, the astrocyte reacts to the glutamate release of a neuron by releasing more glutamate, triggering a glutamate-induced glutamate release (GIGR) similar to the concept of CICR. The authors show that the proposed mechanism accounts for Ca^2+^ bursts in astrocytes.

Other single-cell models are mostly concerned with *IP*_*3*_ dependent Ca^2+^ dynamics. Early models, such as the one proposed by Goldbeter et al. [1990] or Li and Rinzel [1994], use a constant concentration of *IP*_*3*_ to show that Ca^2+^ fluctuations are possible even without oscillation in *IP*_*3*_ level. Later models then started to include more complete *IP*_*3*_-Ca^2+^ dynamics [Goto et al., 2004] and finally included both, the Ca^2+^-dependent synthesis and the degradation of *IP*_*3*_ [De Pittà et al., 2009].

The behavior of different signaling pathways and enzymes is prevalently modeled through ordinary differential equations (ODEs). For example, Taheri et al. [2017] presented a single-compartment model focused on intracellular Ca^2+^ dynamics in an astrocytic compartment. Using ODEs and information from experimental data, they described the influence of *IP*_*3*_ on Ca^2+^ signaling and used their results to categorize four different types of Ca^2+^ transients. A more specific, particle-based model of an astrocytic compartment was implemented by Denizot et al. [2019]. Using their model, the authors were able to recreate stochastic Ca^2+^ signals and showed that the occurrence of Ca^2+^ signals is heavily dependent on the spatial positioning of *IP*_*3R*_ channels. Oschmann et al. [2017] created a model including intracellular Ca^2+^ dynamics and their dependence on both GluT and mGluR, using it to study how the different pathways affect the Ca^2+^ dynamics throughout an astrocytic process. In this study, we will focus on the computational model developed by Oschmann et al. [2017]. The details will be explained further in the next section.

#### 2.2.2 Astrocytic Compartment Model by Oschmann et al. [2017]

Oschmann et al. [2017] developed a single compartment model that takes both aforementioned Ca^2+^ pathways into account: The mGluR-dependent pathway, leading to the production of *IP*_*3*_ and thereby to the exchange of Ca^2+^ between ER and cytosol, and the GluT-dependent pathway, employing glutamate transporters and, together with NCX and NKA, influencing the exchange of glutamate, Ca^2+^, Na^+^ and K^+^ between extracellular space and cytosol. A schematic drawing of the different currents resulting from these pathways is shown in Figure 1.

**Figure 1.**
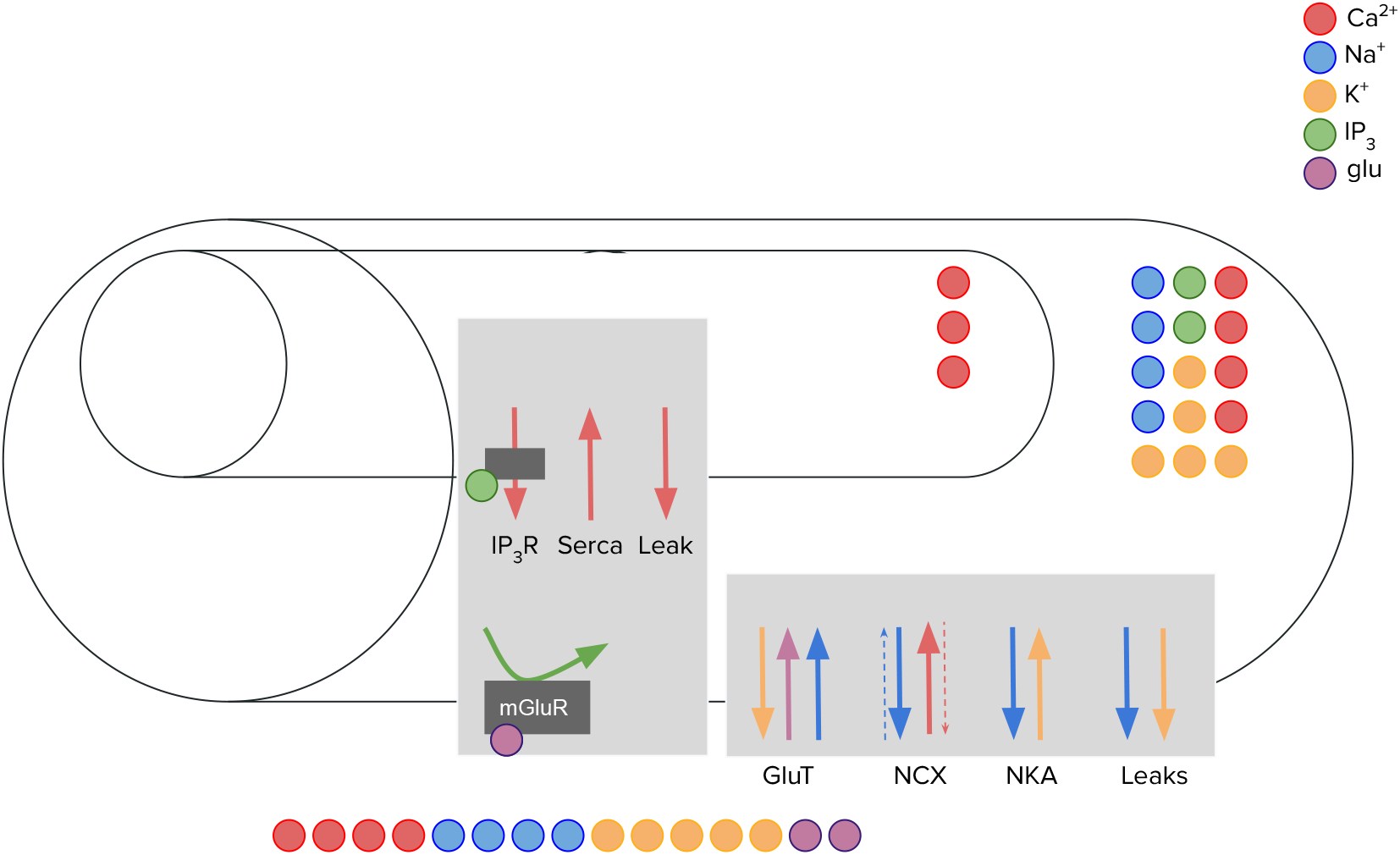
Schematic drawing of the computational astrocytic compartment model as it was implemented by Oschmann et al. [2017]

In this model, the intracellular space of an astrocytic compartment is represented by a cylindrical shape. Another smaller cylinder is placed within the intracellular space representing the ER. The distance between soma and simulated compartment proportionally decreases the volume of the intracellular space, the volume of the ER, and the volume ratio ratio_*ER*_ between the two (Figure 2). For simplicity, the extracellular space is assumed to be exactly on top of the mantel area of the intracellular space. Diffusion between neighboring compartments is not considered.

**Figure 2.**
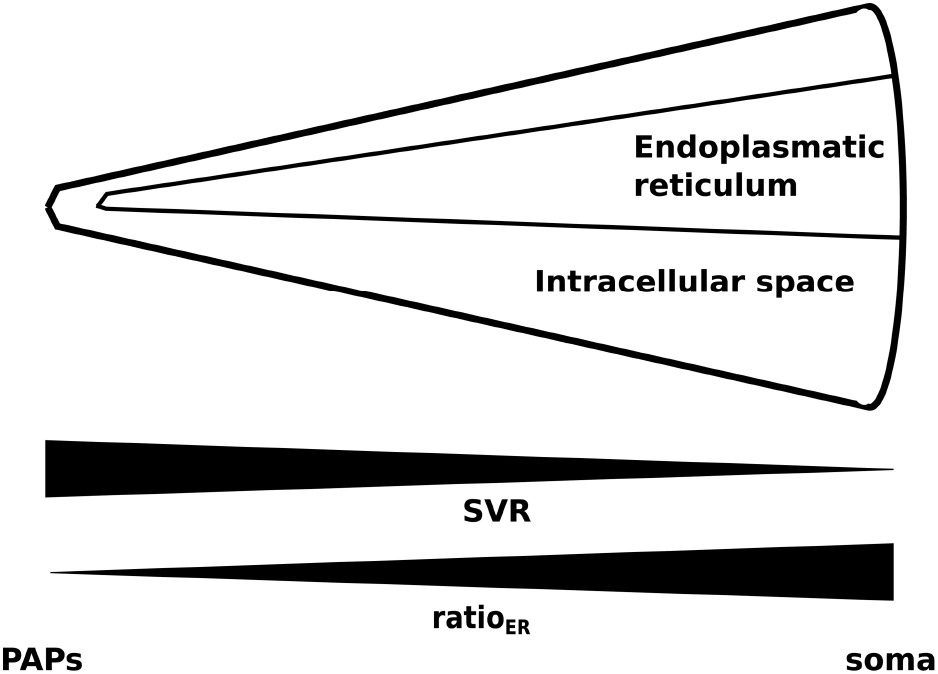
Figure depicting the change in SVR in compartments along an astrocytic process (taken from Oschmann et al. [2017]).

The computational model consists of seven ODEs that describe the change in ion concentrations over time. Each ODE is a weighted sum of particle currents or, in the case of *IP*_*3*_, the production and degradation of *IP*_*3*_. Each current accounts for the change in electrical charge caused by a specific mechanism.

Namely, the following currents are considered:

- For the GluT-dependent pathway:
  – I_GluT_: Based on the transport of glutamate by glutamate transporters. As a byproduct, Na^+^ is transported into and K^+^ out of the intracellular space.
  – I_NCX_: Based on the Na^+^-Ca^2+^ exchanger [Luo and Rudy, 1994].
  – I_NKA_: Based on Na^+^-K^+^ adenosine triphosphatase and a simplified form of its mathematical description [Luo and Rudy, 1994].
  – 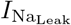 and 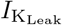 : Leak currents dependent on the current membrane voltage and the Nernst potentials of Na^+^ and K^+^, respectively.
- For the mGluR-dependent pathway:
  – I_*IP3R*_: Ca^2+^ current from the ER into the intracellular space through *IP*_*3*_ receptor channels. It is based on the mathematical description by Li and Rinzel [1994]. The exact current depends on the probability of activated *IP*_*3*_ receptor channels. The probability is modeled through the ODE (Equation 6) described in the next paragraph.
  – I_Serca_: Pump to transport Ca^2+^ from the intracellular space into the ER [Li and Rinzel, 1994].
  – 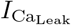 : Leak current out of the ER. It is important to note that, other than 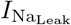 and 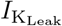, this leak current does not depend on the membrane voltage but on the Ca^2+^ concentration gradient between ER and intracellular space [Li and Rinzel, 1994].

The intracellular and ER Ca^2+^ concentration are computed using the following equations:

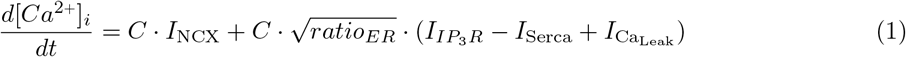

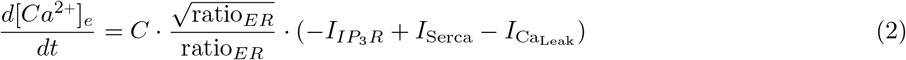

where *C* is a constant accounting for the ratio between the area of the internal Ca^2+^ storage and the volume of the intracellular space. Similarly, the derivatives of intracellular Na^+^ and K^+^ concentrations are defined as:

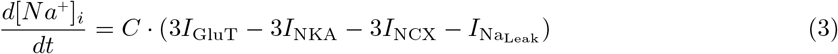

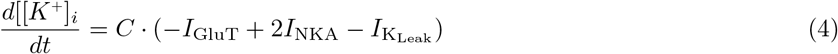

The production and degradation of *IP*_*3*_ are governed by mechanisms dependent on extracellular glutamate and internal Ca^2+^ concentration which are further discussed in the original paper [Oschmann et al., 2017] and a paper by De Pittà et al. [2009]. The amount of internal *IP*_*3*_ directly influences the open probability *h* of *IP*_*3R*_ channels.

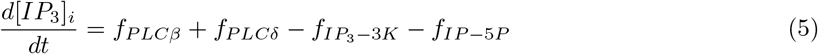

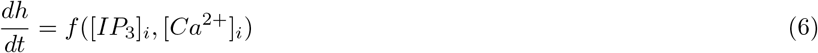

Last, the currents also influence the membrane voltage through the equation

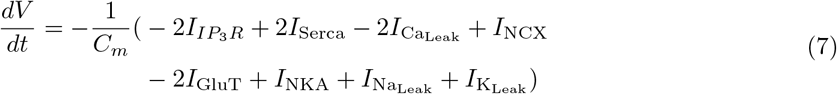

where *C*_*m*_ is the membrane capacitance. The total concentrations of Ca^2+^, Na^+^, and K^+^ are assumed to be constant.

A more detailed description of the computational model can be found in the original article [Oschmann et al., 2017].

### 2.3 Parameter Inference

One of the major challenges in computational modeling is the accurate and efficient estimation of system parameters. Parameters are often not directly transferable from experiment to model or might not be measurable at all. Especially in system biology, parameters might further vary vastly between different species. Hence, a lot of effort has been put into the exploration of appropriate parameter inference methods. Most of these methods can be summarized as algorithms that attempt to minimize an objective function.

One of the simplest and most well-known methods for parameter inference is least squares fitting (LSF). LSF attempts to find the function best describing a set of observations by minimizing the least square error between each observation and the estimated solution. In general, the method is best suited for linear problems without colinearity and with constant variance. In biology, adaptations of LSF have been used for a variety of use cases, including the inference of parameters in S-systems [Liu et al., 2012, Dattner et al., 2019] or biochemical kinetics [Mendes and Kell, 1998].

Genetic algorithms (GA) on the other hand work by assigning *fitness* (value of the objective function) to different, at the beginning randomly generated, samples. The fittest samples are selected and modified by either recombining them with other samples or by randomly mutating them. The process of assigning fitness, selection, and modification is then repeated until samples with sufficient fitness are produced [Mitchell, 1998].

Based on probability theory, Bayesian inference combines prior knowledge with the likelihood of parameters generating the desired output. Respective methods attempt to estimate the parameters and their probability distribution by maximizing the likelihood function. For example, Bayesian inference finds its application in Markov Chain Monte Carlo (MCMC) algorithms. In general, MCMC works by randomly sampling parameter values proportional to a known function. The exact implementation is algorithm-dependent. Recently, Valderrama-Bahamóndez and Fröhlich [2019] studied the performance of different MCMC techniques for parameter inference in ODE-based models.

Kalman filtering is another approach originating from the field of control theory. Kalman filters produce parameter estimates by iteratively interpreting measurements over time and comparing them to their own predictions. These filters often find applications in robotics and navigation. In 2010, Lillacci and Khammash [2010] proposed an algorithm to infer parameters of ODE-driven systems through Kalman filters and proofed their concept on the heat shock response in E. coli and a synthetic gene regulation system. Similarly, Dey et al. [2018] combined Kalman Filters with MCMC to create a robust algorithm for parameter inference in biomolecular systems.

With the growing popularity of deep learning, various attempts have been made to use neural networks for parameter inference. Green and Gair [2020] trained a neural network to closely approximate the posterior distribution of gravitational waves, thereby replacing the more often used MCMC algorithm. At the same time, the concept of physics informed neural networks (PINN) has been introduced by Raissi et al. [2017]. The general idea is to train neural networks on sparse data while enforcing additional constraints modeled through ordinary- or partial differential equations (ODE or PDE). While the first version of PINNs was found to be error-prone by many authors [Wang et al., 2020, Antonelo et al., 2021], the method has since been improved and applied by several researchers. For example, Lagergren et al. [2020] suggested an extension of PINNs that allows for the discovery of underlying biological dynamics even if the exact underlying PDE or ODE is not known. Similarly, Yazdani et al. [2020] suggested a deep learning algorithm that allows for parameter inference using PINNs in systems biology. Additions to make PINNs more suitable for control theory were proposed by Antonelo et al. [2021].

As the most recent method of parameter inference described in this section, PINNs have not been as well studied as other methods. However, preliminary results are promising and show that they have large potential. In contrast to other methods, they allow for the incorporation of previous knowledge of the mechanics underlying different dynamics. In this study, we will use the algorithm proposed by Yazdani et al. [2020] as a foundation to estimate parameters for the previously mentioned computational model of an astrocytic compartment [Oschmann et al., 2017].

### 2.4 Neural Networks

#### 2.4.1 Perceptron

Back in 1958, Frank Rosenblatt proposed the concept of a simple perceptron [Rosenblatt, 1958]. While still very limited in its functionality, the perceptron was able to learn to distinguish between linearly separable classes. To that end, the perceptron took the weighted sum of different inputs. A simple thresholding function (zero if the weighted sum is below `T`, one otherwise) then decided which class the input belongs to. The perceptron was able to learn the needed weights automatically by minimizing the error between actual and sought-after output. Today’s neural networks work very similarly, basically consisting of a multitude of perceptrons.

Mathematically speaking, a single neuron (perceptron) inside a neural network works as follows: Each neuron gets *n* different inputs, denoted as 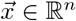. The neuron saves information about the different input weights, denoted as 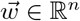, and about its bias, denoted as *b* ∈ ℝ. Figure 3 shows an example of such a perceptron. Weights and bias get optimized throughout the learning process. The relationship between the neuron inputs 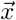 and the neuron output *ŷ* is given through

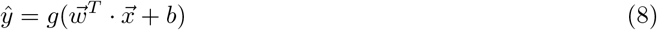

where *g* : ℝ → ℝ is called activation function. Activation functions are functions that map any real single output to a value within a reasonable range. Typical examples include the functions ReLu, sigmoid, and tanh. Figure 4 depicts different activation functions.

**Figure 3.**
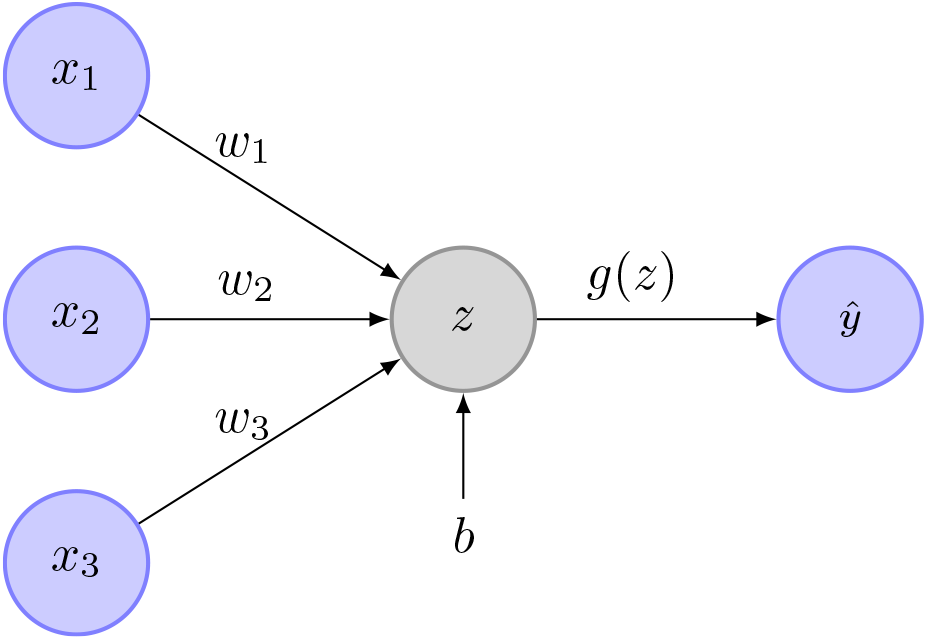
Example image of a perceptron. In this case, *x*_1_, *x*_2_, *x*_3_ are the inputs. The weights are *w*_1_, *w*_2_ and *w*_3_. *b* is the perceptron bias. *z* is the weighted sum of the network inputs and the bias. *g*(*z*) is the activation function. *ŷ* is the perceptrons output.

**Figure 4.**
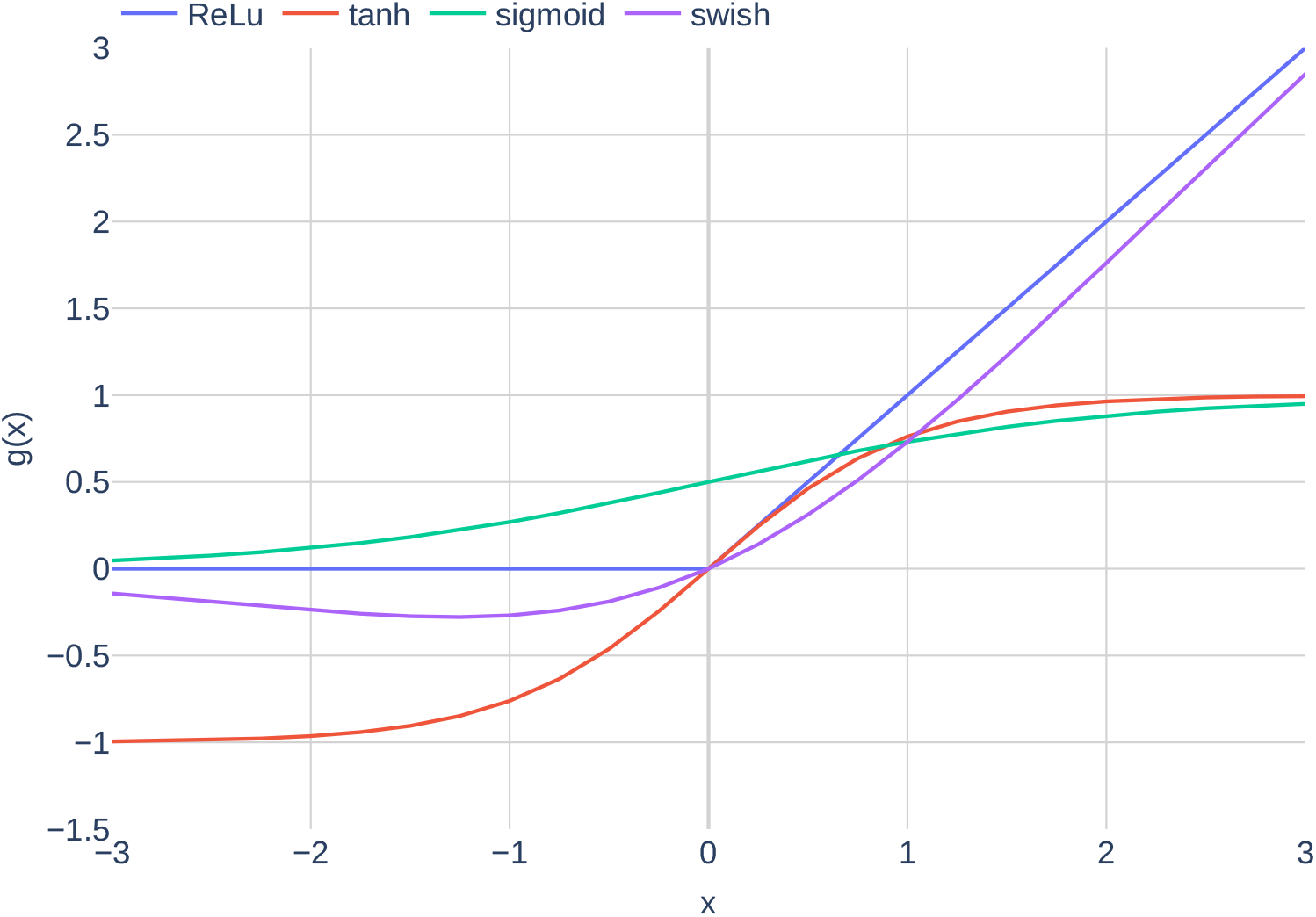
Plots of the different activation functions `ReLu`, `tanh`, `sigmoid` and `swish`.

#### 2.4.2 Full Neural Networks

Usually, a neural network consists of multiple layers *L* of neurons. Inside each layer is a fixed amount of neurons *N*_*l*_. While neurons inside the same layer are not connected with each other, each neuron of layer *l* is connected with each neuron of layer *l* − 1 and with each neuron of layer *l* + 1. The output equation for a single perceptron (Equation 8) can be written in matrix form for each layer, resulting in

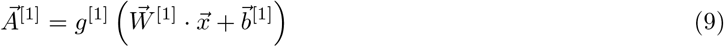

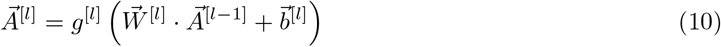

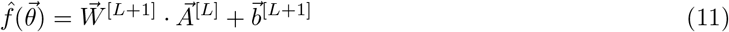

where 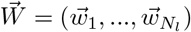 is a matrix containing all input weights, 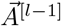 are the output values of the previous layer, 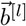 are the biases and *g*^[*l*]^ is a function that applies the chosen activation function component-wise. Figure 5 shows an example of a fully connected network. Note that in this specific example, no activation function is used between the last neural network layer and the output layer. Depending on the desired type of output, this can vary.

**Figure 5.**
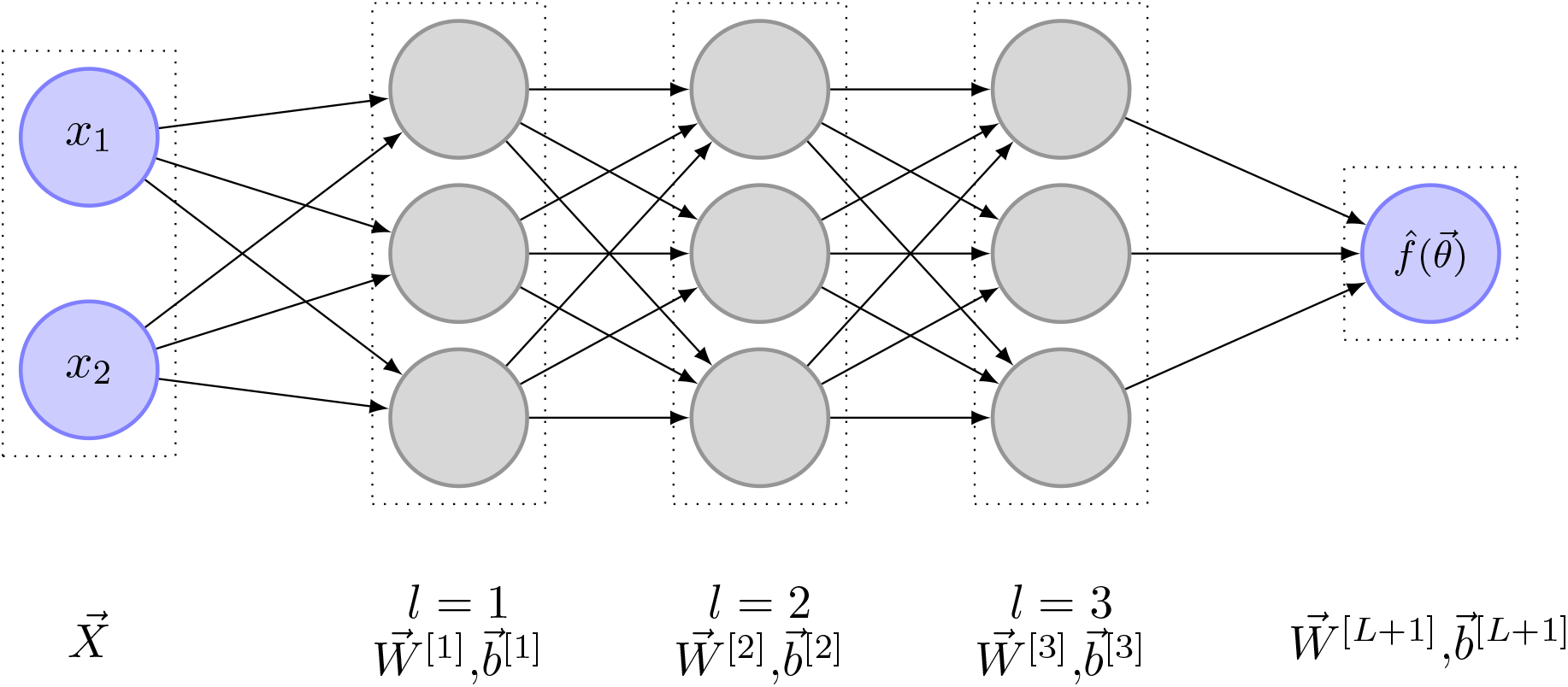
Fully connected neural network with *L* = 3

To learn how weights and biases have to be changed to get the best possible results, the concept of backpropagation is applied. Backpropagation works as follows: First, a loss function ℒ measuring the wrongness of the network is defined. Typical loss functions include mean squared error for regression tasks or cross-entropy for classification tasks. Next, the gradient of the loss with respect to the network weights 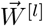 and biases 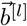 is computed. For simplicity, the combination of all weights and all biases is usually written in vector form 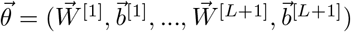, which means that the gradient of the loss function can be written as 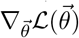. Generally, there is a multitude of different methods to update the network weights given 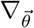, the simplest one being stochastic gradient descent (`SGD`). With `SGD`, the network parameters are updated using

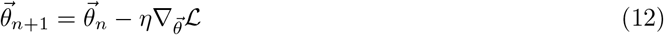

where η is the learning rate and *n* is the current iteration. A more modern adaptation of `SGD` is called `Adam` [Kingma and Ba, 2014]. In contrast to `SGD, Adam` is an adaptive gradient descent algorithm that maintains a learning rate per-parameter and is, therefore, less sensitive to the set learning rate η. Furthermore, it uses the first and second moments of the gradient to speed up convergence where possible.

#### 2.4.3 Systems biology informed deep learning for inferring parameters by Yazdani et al. [2020]

Yazdani et al. [2020] suggested a deep learning model for inferring parameters and hidden dynamics in biological models governed by ODEs. Using only a few, incomplete and noisy measurements, they were able to accurately estimate unknown model parameters.

In their algorithm, Yazdani et al. [2020] assumed a computational model with *s* states 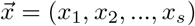 of which *m, m* ≤*s*, states are observable. Each state is described through one ODE. Therefore, the system of ODEs can be described by

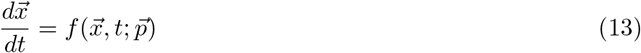

where 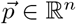 are the *n* unknown model parameters. Using neural networks, they then attempt to learn a surrogate function 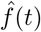 that maps measurement times to the state variables.

In addition to the usual neural network layers (input layer, hidden layers, output layer), they extended the network by three additional layers. The first two layers are added in between the input- and hidden layers. The first one is an input scaling layer, that scales the timestamps to be between zero and one. Second, a feature layer is added. This layer transforms the scaled input time to a function that already roughly describes the function the network is supposed to learn. For example, if the state variables oscillate heavily, sin(*t*) might be used as a feature transform. The last layer is added behind the output layer and is responsible for scaling the output states to be approximately of magnitude 𝒪(1). A schematic drawing of the different network layers can be seen in Figure 6.

**Figure 6.**
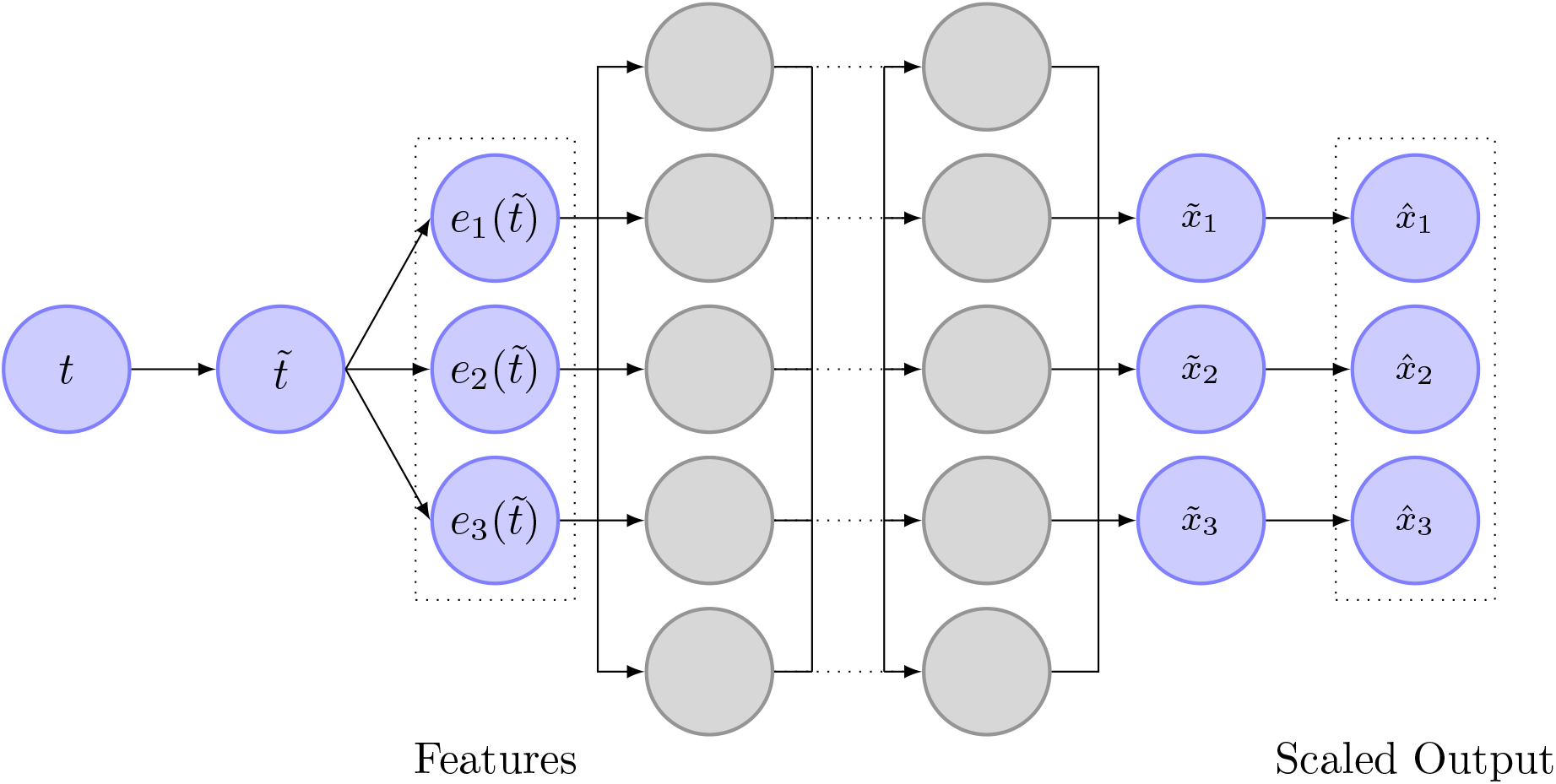
Image depicting the structure of the neural network as it is used by Yazdani et al. [2020]. The neural network uses measurement timestamps as input. In the first layer, the timestamp is normalized to be between zero and one. In the next layer, the scaled time is transformed according to prior knowledge about the state variables. The normal, fully connected layers are depicted next as gray circles. The output of the network is then scaled to ensure that 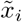 is approximately of magnitude 𝒪(1).

As mentioned earlier, neural networks learn by attempting to reduce a loss function. In this algorithm, the loss function is defined as

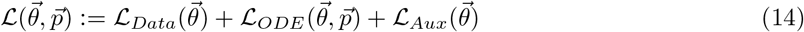

The different loss terms have the following meaning:

- ℒ_*Data*_: The weighted mean squared error (MSE) between the observed states 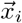 and their respective state outputs 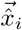 of the neural network.
- ℒ_*Aux*_: The weighted MSE between initial and end values of the original states 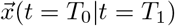 and initial and end values of the neural network output 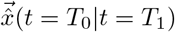
- ℒ_*ODE*_ : The weighted MSE between the gradient of the learned function with respect to time 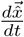 and the gradients given by the computational model, 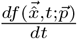. The term 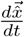 is computed through automatic differentiation.

Using these loss terms, the neural network is able to learn both, an approximation of the function *f* and the unknown parameters 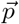.

Using this algorithm, Yazdani et al. [2020] were able to infer hidden dynamics and parameters from noisy data in a standard yeast glycolysis model, in a cell apoptosis model, and even in an event-driven ultradian endocrine model. While inference on the last model worked best when event times were known, parameter inference was still reasonably successful without. However, it is important to note that the suggested setup does not allow for the generalization of inference or measurements when the computational model is event-driven. A more detailed description of the algorithm can be found in the original article [Yazdani et al., 2020]. More details regarding the implementation will be given in the next section.

## 3 Methods, Part 1

In this section, we detail the implementation of the Oschmann et al. [2017] model and the parameter inference algorithm originally developed by Yazdani et al. [2020].

### 3.1 Tools

All code was written in `Python` 3.8.1. The well-known libraries `numpy, scipy`, and `pandas` were used to aid with different aspects of the implementation. The plotting library `plotly` was used for result visualization.

While the original deep learning paper referred to in this manuscript, [Yazdani et al., 2020] used the machine learning library `TensorFlow` in combination with `DeepXDE` [Lu et al., 2021], we chose to use `PyTorch 1.8.1` instead. In contrast to `Tensorflow, PyTorch` is more object-oriented (OOP) and usually more intuitive to understand and modify. Runtime experiments were performed on a local computer with Ubuntu 20.04, an AMD 6 core CPU, and a high-end NVIDIA graphics card.

### 3.2 Model by Oschmann et al. [2017]

In this section, we shortly detail changes made to the original Oschmann et al. [2017] model. Furthermore, we explain how the ODEs from the Oschmann et al. [model are integrated and where the parameter sets used originate from.

#### 3.2.1 Conceptual Changes to the Model

We made two minor changes to the computational model of an astrocytic compartment. First, we noticed that other computational models only consider charge fluxes between intra- and extracellular space when computing membrane voltage [Farr and David, 2011, Witthoft and Em Karniadakis, 2012]. Since fluxes between the ER and the cytosol do not change the total charge of the intracellular space, we removed currents originating from the mGluR-dependent pathway from the membrane voltage ODE in Equation 7, resulting in a new ODE of the form

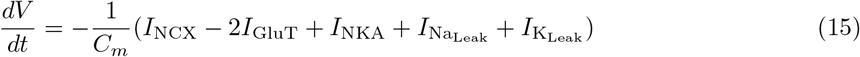

where *C*_*m*_ is the membrane capacitance.

Second, we modified Equation 1 to incorporate the two times positive valence of Ca^2+^, resulting in:

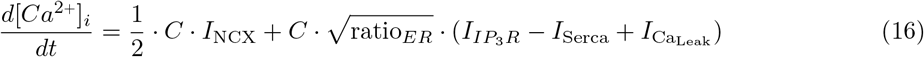

In this equation, *C* is a constant accounting for the ratio between the area of the internal Ca^2+^ storage and the volume of the intracellular space.

#### 3.2.2 Integration Method

As mentioned earlier, the Oschmann et al. [model consists of seven highly nonlinear ODEs that describe the behavior of different molecules within an astrocytic compartment. Using a glutamate stimulation train and a time frame as input, the computational model integrates the ODEs and gives concentrations ([*Ca*^*2+*^]_*i*_, [*Ca*^*2+*^]_*e*_, [*K*^*+*^]_*i*_, [*Na*^*+*^]_*i*_, [*IP*_*3*_]_*i*_), open probability of *IP*_*3R*_ channels (*h*) and membrane voltage (*V*_*m*_) as output at each timestep. The integration is done using the `scipy` function `solve_ivp`.

While `solve_ivp` allows for many different integration methods, we chose the implicit multi-step variable order method BDF [Shampine and Reichelt, 1997]. This decision is based on the observation that the described system of ODEs is stiff. Another stiff solver offered by `scipy` is Radau [Hairer and Wanner, 1996]. However, BDF is known to perform better if evaluating the ODEs in itself is expensive, as is the case in the computational model at hand. We used a relative tolerance of 1e^−9^ and an absolute tolerance of 1e^−6^.

#### 3.2.3 Parameter Configuration

As part of this work, we tested different parameter sets. The first parameter set included the parameters as they were in the original paper (parameter set `Paper`). The second parameter set slightly differed from the first one and included parameters according to the doctorate thesis by Oschmann [2018] (parameter set `Thesis`). The third parameter set is seen as the default parameter set and is used unless otherwise indicated (parameter set `Default`; based on a personal communication between Franziska Oschmann and Kerstin Lenk, 08.11.2018). The differences in parameter sets are listed in Table 1. A simple configuration mechanism that allows for modifying, loading, and saving different parameter sets is provided.

**Table 1.** Different values for the three parameter sets `Default, Paper`, and `Thesis`

| Parameter | Unit | Default | Paper | Thesis | Description |
| --- | --- | --- | --- | --- | --- |
| **Initial Values** |  |  |  |  |  |
| $[Ca^{2+}]_e$ | mM | 0.01963 | 0.025 | 0.019 | $Ca^{2+}$ in ER |
| $[Na^+]_{tot}$ | mM | 165 | 160 | 160 | Tot. $Na^+$ available |
| $[Na^+]_{out}$ | mM | 150 | 145 | 145 | $Na^+$ in extracellular space |
| $V_m$ | V | -0.08588 | -0.085 | -0.085 | Membrane voltage |
| **Parameters** |  |  |  |  |  |
| $I_{GluTmax}$ | $\frac{pA}{\mu m^2}$ | 0.75 | 0.68 | 0.75 | Max. current of $I_{GluT}$ |
| $I_{NCXmax}$ | $\frac{pA}{\mu m^2}$ | 0.001 | 0.1 | 0.1 | Max. current of $I_{NCX}$ |
| $g_{NaLeak}$ | $\frac{\mu S}{\mu m^2}$ | 13 | 6.5 | 13 | Conductance of $Na^+$ leak |
| $g_{KLeak}$ | $\frac{\mu S}{\mu m^2}$ | 162.46 | 79.1 | 162.46 | Conductance of $K^+$ leak |
| $\nu_\beta$ | $\frac{mM}{s}$ | 1e-4 | 5e-5 | 5e-5 | Max. production of $IP_3$ by PLC $\beta$ |
| $r_L$ | $\frac{1}{s}$ | 0.055 | 0.11 | 0.11 | $Ca^{2+}$ leak rate between ER and intracellular space |
| $r_C$ | $\frac{1}{s}$ | 3 | 6 | 6 | Max. CICR rate |
| $\nu_{ER}$ | $\frac{mM}{s}$ | 0.0045 | 0.004 | 0.004 | Max. $Ca^{2+}$ uptake by SERCA |

### 3.3 Adaptation of the Deep Learning Model by Yazdani et al. [2020]

In this section, we detail the methods and equations used to do parameter inference using the algorithm by Yazdani et al. [2020]. We show how the algorithm has to be adapted for the astrocytic compartment model, discuss implementation details not mentioned in the original paper, and highlight changes.

#### 3.3.1 Configuration of the Neural Network

Figure 7 shows a schematic drawing of the neural network algorithm as proposed by Yazdani et al. [2020] implemented for the Oschmann et al. [model. As mentioned previously, the Oschmann et al. [model consists of seven different ODEs. Therefore, the neural network has seven output nodes. If not otherwise indicated in parameter inference experiments, the neural network itself consists of 4 network layers with 150 nodes each. Weights and biases are initialized with random values from a truncated normal distribution, called Glorot normal distribution, centered around zero [Glorot and Bengio, 2010].

**Figure 7.**
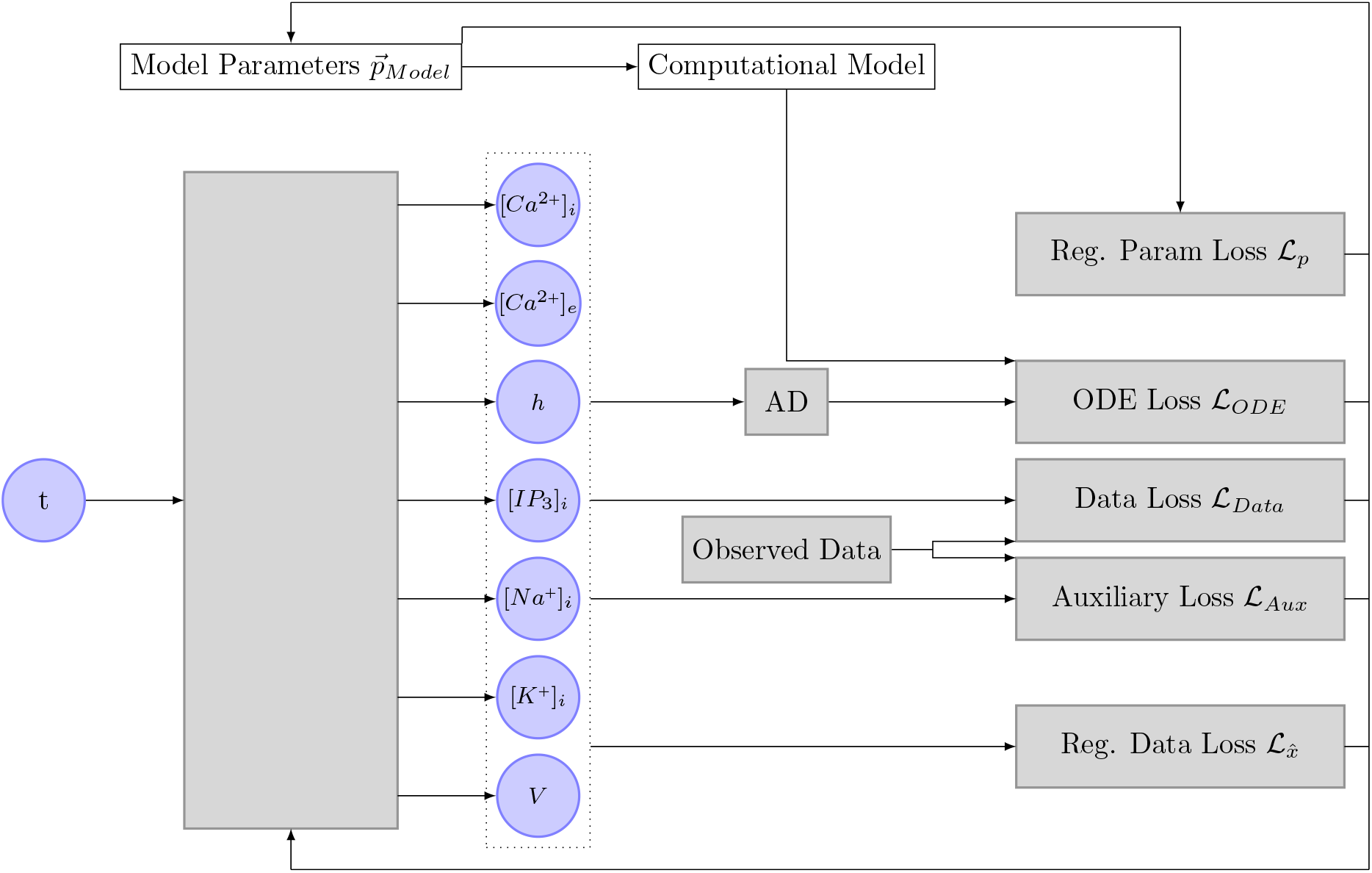
This Figure shows the implementation of the algorithm initially proposed by Yazdani et al. [2020] in the context of this thesis. The neural network takes the input time of a measurement as an input and outputs the seven different state variables. These state variables, together with the inferred parameters, the computational model, and the observed data are used to compute the different loss functions. The gradient of these loss functions is then used to optimize the inferred parameter and the neural network. AD stands for automatic differentiation.

we used the activation function `swish` [Ramachandran et al., 2017], which is defined as

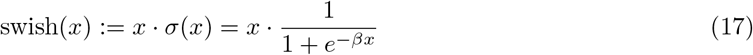

with *β* = 1. This activation function was introduced by Google in 2017 and has been shown to perform better than the more commonly known activation functions `ReLu` and `sigmoid`. The performance improvement is mostly attributed to the unboundedness of the function. The previously shown Figure 4 includes a plot of the activation function `swish`. In contrast to the original authors [Yazdani et al., 2020], we decided to shuffle the data and create batches of size *N*. In general, shuffling of input data is considered to be good practice. Furthermore, the usage of a fixed batch size circumvents that the learning rate has to be adapted according to the size of the data set.

#### 3.3.2 Input- and Feature Transform

As in the original paper [Yazdani et al., 2020], we added an input scaling and a feature transform layer. The input time *t* was linearly scaled to be between zero and one. Setting *T*_0_ to be the smallest time in the measurement data and *T*_1_ to be the largest time, the scaled time was therefore defined as

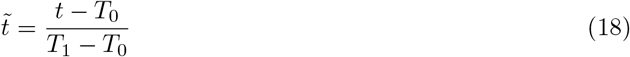

It is important to note that the time should be scaled as part of the neural network. Scaling the time beforehand, for example, to seemingly decrease complexity, leads to incorrect derivatives when automatic differentiation is applied to the neural network.

The goal of the feature transform layer is to add prior knowledge about the time response of the different state variables to the neural network, thereby accelerating learning. For the computational model at hand [Oschmann et al., 2017], we chose the feature transform

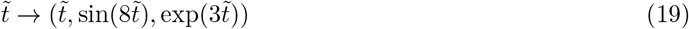

based on the observation that some state variables ([*Na*^*+*^]_*i*_, [*K*^*+*^]_*i*_, *V*) behave like step functions and the repeated exponential growth of the intracellular Ca^2+^ concentration ([*Ca*^*2+*^]_*i*_).

#### 3.3.3 Output Transform

In the code accompanying the original paper [Yazdani et al., 2020], the output transform of the network is implemented as follows:

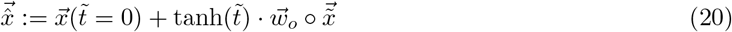

where 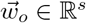 is a vector accounting for the different orders of magnitude, 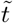 is the scaled time and is the Hadamard product (component-wise multiplication). While this output transform works, it has one major underlying problem. It requires the initial state 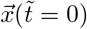 to be known exactly. Since tanh 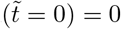, the gradient of the data- and auxiliary loss 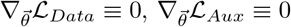 with respect to the neural network parameters will always be zero. It follows that the network can not learn from the observed data at 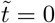. For some state variables, it might not be possible to observe the initial state, leaving the network with an uncorrectable error. We, therefore, implemented the simpler and computationally less expensive output transform function

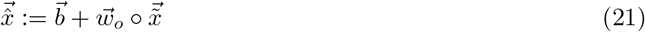

where 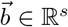 is a vector allowing for prior knowledge about the starting conditions to be incorporated into the network. In contrast to the previous transform function, however, 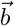 can be noisy or set to 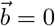 without limiting the network’s ability to learn. Further, all data is prioritized equally, independent of time.

Both transform functions have the disadvantage that 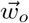 has to be set manually. The weights 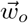 used throughout this study are based on the mean values of the different state variables and are listed in Table 2. The mean values are shown in the Appendix in Table **??** and Table **??**.

**Table 2.** This table lists the state variable-related weights used for the deep learning algorithm.

| State Variable | $\vec{w}_o$ | $\vec{w}_{\text{Data}}$ | $\vec{w}_{\text{ODE}}$ |
| --- | --- | --- | --- |
| $[Ca^{2+}]_i$ | 1e-04 | 1e+04 | 1e+04 |
| $[Ca^{2+}]_e$ | 1e-02 | 1e+02 | 1e+03 |
| $h$ | 1e-01 | 1e+01 | 1e+02 |
| $[IP_3]_i$ | 1e-04 | 1e+04 | 1e+04 |
| $[Na^+]_i$ | 1e+01 | 1e-01 | 1e-01 |
| $[K^+]_i$ | 1e+02 | 1e-02 | 1e-02 |
| $V$ | -1e-01 | 1e+01 | 1e-01 |

#### 3.3.4 Loss Function

In the following, we shortly explain changes and additions made to the originally used loss function [Yazdani et al., 2020], before giving the exact loss formulas used throughout this study.

##### Mean Squared Error

In the original paper, the authors use the following definition of weighted MSE:

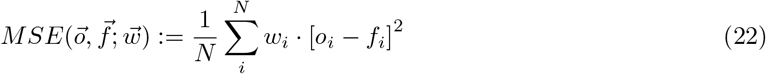

where 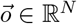 is the expected output and 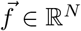 is the computed output. The vector 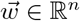 is used to scale the different state variables to approximately the same order of magnitude.

In practice, we found that setting appropriate weights is more intuitive when using the following definition:

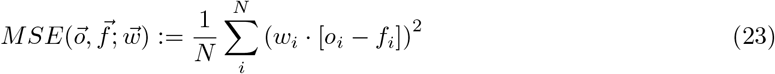

The weights used in this manuscript are listed in Table 2. Column 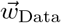 is used when computing the MSE of the observed data. Column 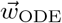 is used when computing the MSE of the automatically differentiated network output in comparison to the ODEs computed by the Oschmann et al. [model.

##### ODE Loss

As mentioned earlier, ℒ_*ODE*_ is the weighted MSE between the gradient of the neural network with respect to time and the gradients given by the computational Oschmann et al. [model. The assumption is that ℒ_*ODE*_ is minimized when the learned dynamics and the inferred parameters are correct. To compute ℒ_*ODE*_, the Oschmann et al. [model is fed with the output of the neural network 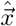 and the current parameter assumptions at each iteration. Similar to the neural network outputs, Yazdani et al. [2020] suggested scaling the model parameters to be approximately of scale 𝒪(1). The scalings used throughout this study are listed in Table 3. The gradient of the neural network 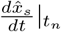 is computed using the automatic differentiation function `autograd.grad` from the machine learning library `PyTorch`.

**Table 3.** Parameter values (`Default`), their scaling, and feasible parameter ranges that are used throughout this study.

| Parameter | Unit | Original | Scaling | Range | Description |
| --- | --- | --- | --- | --- | --- |
| $I_{\text{NKA}}$ | | | | | |
| $K_{\text{NKAmN}}$ | $mM$ | 10 | 1 | $[0, \infty]$ | Half saturation of $\text{Na}^+$ |
| $I_{\text{NCX}}$ | | | | | |
| $I_{\text{NCXmax}}$ | $\frac{pA}{\mu m^2}$ | 0.001 | 0.01 | $[0, \infty]$ | $\text{Ca}^{2+}$ max. current |
| $I_{\text{Serca}}$ | | | | | |
| $\nu_{ER}$ | $\frac{mM}{s}$ | 0.0045 | 0.001 | $[0, \infty]$ | $\text{Ca}^{2+}$ max. uptake |
| $K_{ER}$ | $mM$ | 0.0001 | 0.001 | $[0, \infty]$ | $\text{Ca}^{2+}$ affinity |

##### Auxiliary Loss

In physics informed deep learning, the idea of auxiliary loss origins from the concept of Dirichlet boundary conditions. For example, when attempting to learn the solution to the stationary heat equation, one might want to enforce the temperature next to known heat sources. However, in the field of computational biology, the auxiliary loss ℒ_*Aux*_ might not be suitable as it requires the state variables 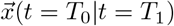 to be known at the beginning and the end of the experimental data. To ensure the algorithm can still be used and still delivers good results when this data is not available, we created a flag σ_Aux_ with which the auxiliary loss can be disabled.

Since we shuffle the data and only use batches of size *N* = 32, the learning batch will often not contain the timestep *t* = 0. To circumvent this problem, we added the data point *t* = 0 manually for each learning step.

##### Regularization Loss

In their original paper, Yazdani et al. [2020] suggest speeding up the convergence process by first training the network on the supervised losses ℒ_*Data*_ and ℒ_*Aux*_ only, before adding the unsupervised learning of the computational model parameters. While this method does indeed speed up the convergence of the network, we found it to lead to one significant problem: The neural network learned the output of the observed state variables without considering the implications for unobserved state variables, leading to infeasible predictions which interfered with the evaluation of the computational model once ℒ_*ODE*_ was added.

To counteract this behavior, we added a soft regularization to the state variables, constraining their feasible range. The regularization mechanism is expressed through a function ℜ defined as:

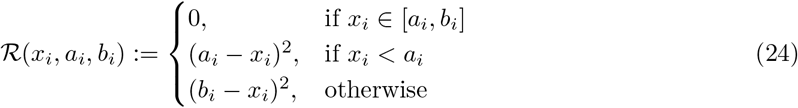

where *x*_*i*_, *i*∈ { 1, …, *s*} is the considered variable, *a*_*i*_ is the lower range boundary and *b*_*i*_ is the upper range boundary. In words, ℜ evaluates to zero if the state variable is within range. Otherwise, ℜ returns the square distance between the closest range boundary and the current value. This regularization function is used in an additional loss function 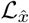. The exact formulation is given in the following section. We added the same mechanism for the inferred network parametersℒ_*p*_, thereby allowing for the incorporation of prior knowledge and avoiding biologically illogical minimas.

For experimental purposes, the ODE loss and the regularization losses can be enabled or disabled through the respective flags σ_*ODE*_, 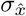 and σ_*p*_. The feasible ranges for the state variables are listed in Table 4, and the feasible ranges for parameters in Table 3.

**Table 4.** Feasible ranges for the different state variables. The ranges are used to compute the regularization loss.

| State Variable | Minimum | Maximum |
| --- | --- | --- |
| $[Ca^{2+}]_i$ | 0 mM | 1e-02 mM |
| $[Ca^{2+}]_e$ | 0 mM | 1e-01 mM |
| $h$ | 0 | 1 |
| $[IP_3]_i$ | 0 mM | 1e-02mM |
| $[Na^+]_i$ | 5e+00mM | 4e+01mM |
| $[K^+]_i$ | 5e+01mM | 103mM |
| $V$ | -2e-01V | 0V |

##### Weighting

Although not explicitly mentioned in the paper, the code by Yazdani et al. [2020] shows that the different loss terms ℒ_*Data*_, ℒ_*Aux*_, and ℒ_*ODE*_ are not only weighted to account for different orders of magnitude but also give varying weight to the different loss functions. In my own implementation, we chose to weight the data loss with 98% and the auxiliary and ODE loss with 1% each.

##### Complete Loss Function

Taken all together, the changed loss function now reads

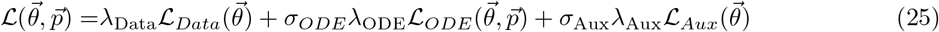

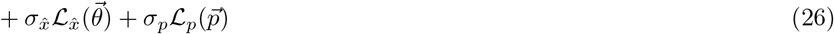

Assuming a batch size of *N* and *S* different state variables of which the first *M* are observable, the different loss terms are defined as

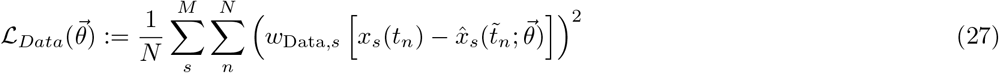

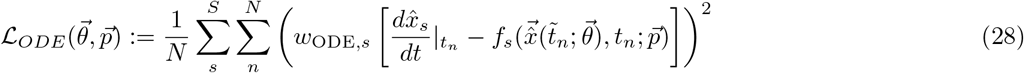

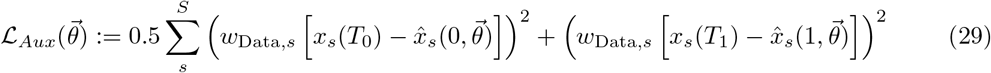

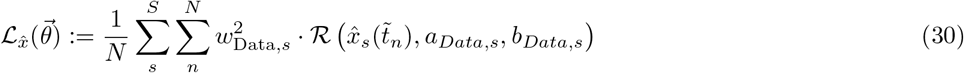

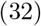

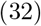

Again, special care has to be taken regarding the timestamp: While the network learns the output with respect to scaled time, automatic differentiation and computational model relay on unscaled time.

#### 3.3.5 Stabilization of the Learning Process

To stabilize the learning process, we employed two methods not initially considered in the original paper [Yazdani et al., 2020]. First, we included the possibility of automatic learning rate reduction. Second, we extended the update step of the neural network with gradient clipping.

##### Learning Rate Reduction

If the learning rate η of a neural network optimization is too large, a network might fail to learn because it keeps overshooting the minimal region. At the same time, if the learning rate is too small, the network might take too long to converge to an appropriate solution. A solution to that problem is learning rate reduction strategies. In this manuscript, we decided to use a learning rate reduction strategy that reduced the learning rate once it has not decreased for a fixed number of epochs. The respective number is called `patience`. The learning rate reduction is implemented using the learning rate model called `ReduceLROnPlateau` implemented in the library `PyTorch`. Unless otherwise indicated, we reduced the learning rate by a factor of 0.5 if the learning rate had not decreased for 5000 epochs.

##### Gradient Clipping

Figure 8 depicts the gradient norms computed during 60000 epochs of network training with the deep learning algorithm described in this section. It can be seen that most gradient norms are within a reasonable range. However, occasionally occurring highly inaccurate network predictions cause far larger gradient norms that disturb the learning process or, in some cases, even cause overflows that render the currently used neural network useless.

**Figure 8.**
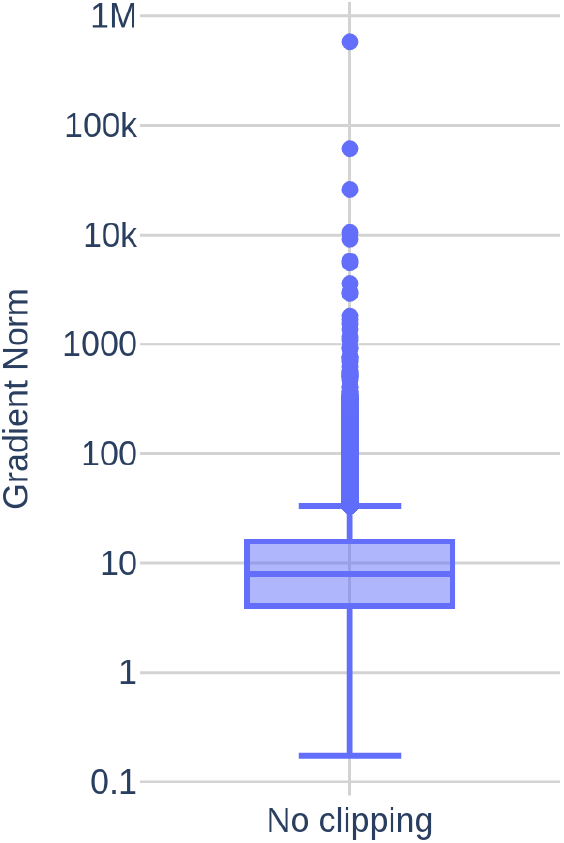
Gradient norms computed during 60000 epochs of network training.

Gradient clipping is a mechanism often employed to avoid these predictions disturb the training process too much. The basic idea is to scale the norm of 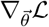 to a maximum value *c* if it is larger than *c*. In my experiments, we found *c* = 10 to work best. The gradient clipping is done through the function `clip_grad_norm_` from the `PyTorch` library.

#### 3.3.6 Complete Algorithm

Algorithm 1 gives an overview of the deep learning algorithm described in this section. The algorithm starts by loading all observed data and by initializing the necessary models. After that, the learning process begins. For `n_epochs`, the algorithm loads the whole data set in batches of size *N* = 32 and feeds them into the neural network to predict 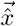. Together with the neural network parameters and the inferred parameters, the predicted data is used to compute the different loss terms and eventually the total loss ℒ and its gradient 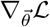. If gradient clipping is enabled, the norm of 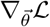 is clipped as described in Section 3.3.5. Afterwards, the neural network parameters and the inferred parameters are changed according to the chosen optimization technique (`Adam` or `SGD`). The variable `mean_loss` is used to compute the mean loss value in the current epoch and to reduce the learning rate as necessary as described in Section 3.3.5. At the end of the algorithm, all generated data and the created neural network model are saved.

### 3.4 Inference Setup

In this section, we specify the used artificial data sets and define the term accuracy. An overview of the different neural network parameters and configurations used is given in Table 5.

**Table 5.** Overview of the different neural network parameters used.

| Parameter | Value | Description |
| --- | --- | --- |
| Network |  |  |
| num layers | 4 | Number of hidden neural network layers |
| num nodes | 150 | Number of nodes per hidden neural network layer |
| $N$ | 32 | Batch size |
| $c$ | 100 | Maximum norm for clip gradient |
| $T_0$ | 0s | Used for scaling of $t$ |
| $T_1$ | 50s | Used for scaling of $t$ |
| Learning Rate |  |  |
| $\eta$ | 0.001 | Learning rate |
| reduction factor | 0.5 |  |
| patience | 5000 |  |
| min lr | 1e-6 | Minimal learning rate |

#### 3.4.1 Data Sets

we generated results using two different data sets:

1. The data set `Parameter Study` consists of 600 data points from 50s of simulation with the computational astrocyte model. The timestamps are spaced evenly and the data is assumed to be noise free. We used this data set to test different neural network configurations.
2. The data set `Noise` is identical to the previous data set. However, in comparison to `Parameter Study`, we added 10% Gaussian noise to the data, resulting in a more realistic data set.

Unless otherwise indicated, we assumed that the glutamate stimulation causing the Ca^2+^ signals is known. The concentration of the glutamate stimulus over time is shown in Figure 9. To study the stability of the deep learning algorithm, we experimented with different amounts of observed state variables, and the data sets were reduced accordingly.

**Figure 9.**
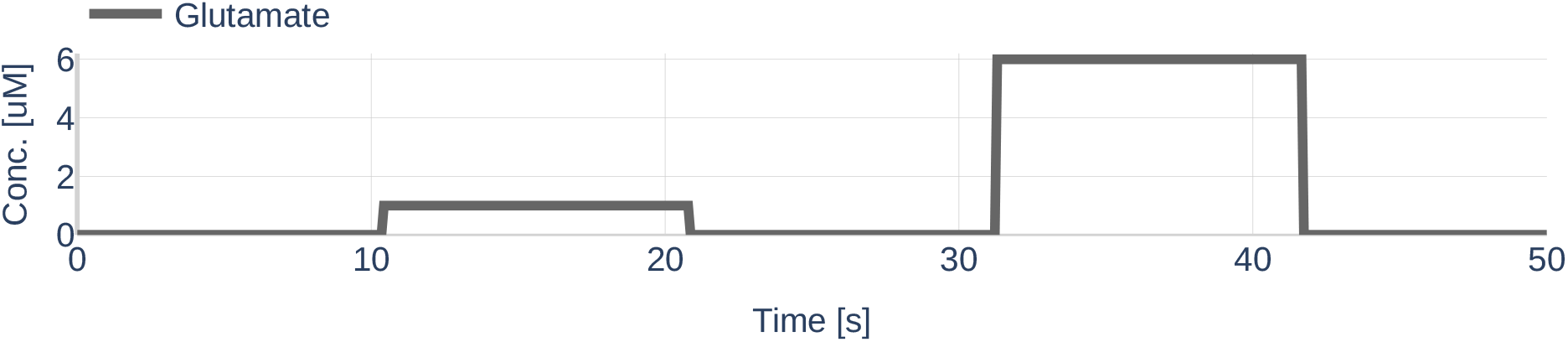
Concentration for the glutamate stimulus used to simulate the astrocytic compartment. The value range was taken from the paper by De Pittà et al. [2009].

##### Algorithm 1

Overview over the deep learning algorithm described in this section

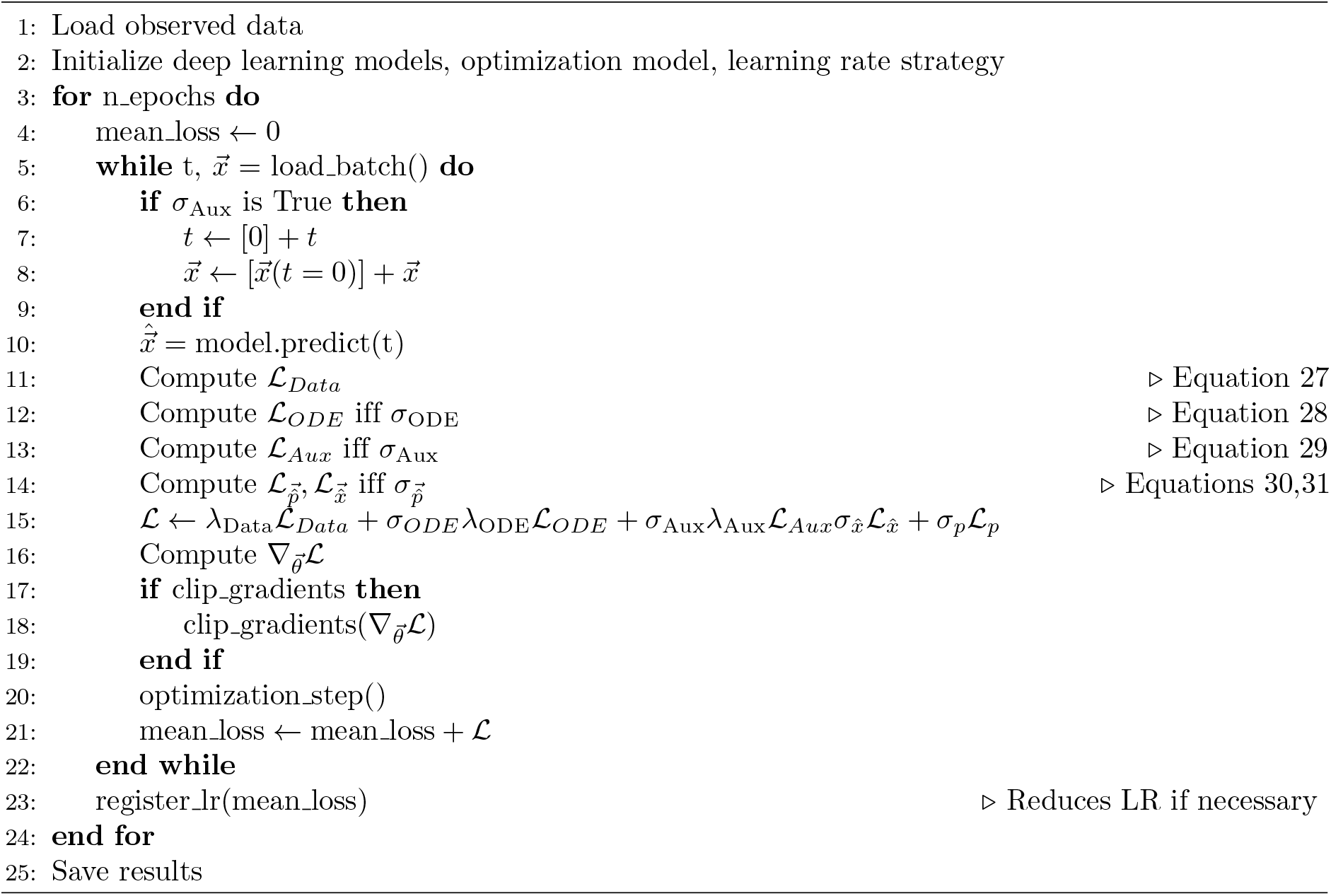

During testing, each data set was randomly split 80/20 into a training- and a validation set. The neural network was only trained on the training set, accuracy reports were made on the validation set. Figures were created by predicting data on complete data sets.

#### 3.4.2 Accuracy

We measured two different kinds of accuracy: The first type describes the accuracy of the dynamics of the different state variables 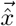. A state variable at time *t* is assumed to be inferred correctly if there is not more than 5% deviation from the original, noise-free, value.

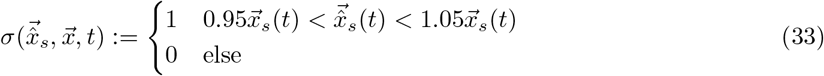

The reported accuracy scores 𝒜_obs_ and 𝒜_all_ then describe the mean accuracy overall measurement times of the observed or overall existing state variables, respectively. Therefore,

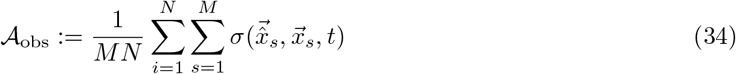

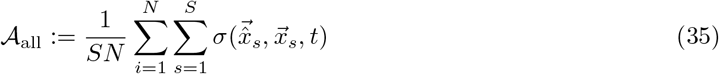

where *S* is number of different state variables, *M* is the number of the observed state variables and *N* is the number of different measurement times *t*.

The second type is concerned with the accuracy of inferred parameters. The accuracy of an inferred parameter is defined as

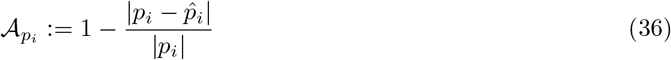

where 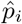 is the inferred parameter and *p*_*i*_ the corresponding real value. Reported is the mean accuracy 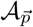 of all inferred parameters.

## 4 Results, Part 1

In this section, we show the dynamics resulting from the ODEs of the Oschmann et al. [model and discuss the influence of the different types of currents.

### 4.1 Model by Oschmann et al. [2017]

First, we describe the dynamics and currents resulting from the Oschmann et al. [model and highlight the influence of the conceptual changes and of the different parameter sets.

#### 4.1.1 Dynamics

First, we studied the temporal evolution of the state variables ([*Ca*^2+^]_*i*_, [*Ca*^2+^]_*e*_, *h*, [*IP*_3_]_*i*_, [*Na*^+^]_*i*_, [*K*^+^]_*i*_, *V*_*m*_) given a specified glutamate stimulus. The influence of the differently made conceptual changes and the different parameter sets will be discussed in the following sections. The results are depicted as colored full lines in Figure 10. The behavior of [*Ca*^2+^]_*i*_ can be described as a repeated pattern of rapid increases and decreases in concentration. The amplitude and the frequency are higher when a glutamate stimulus is present. The increase in [*Ca*^2+^]_*i*_ is always correlated with a drop of Ca^2+^ in the ER. When [*Ca*^2+^]_*i*_ decreases, the [*Ca*^2+^]_*e*_ raises back to its initial value.

**Figure 10.**
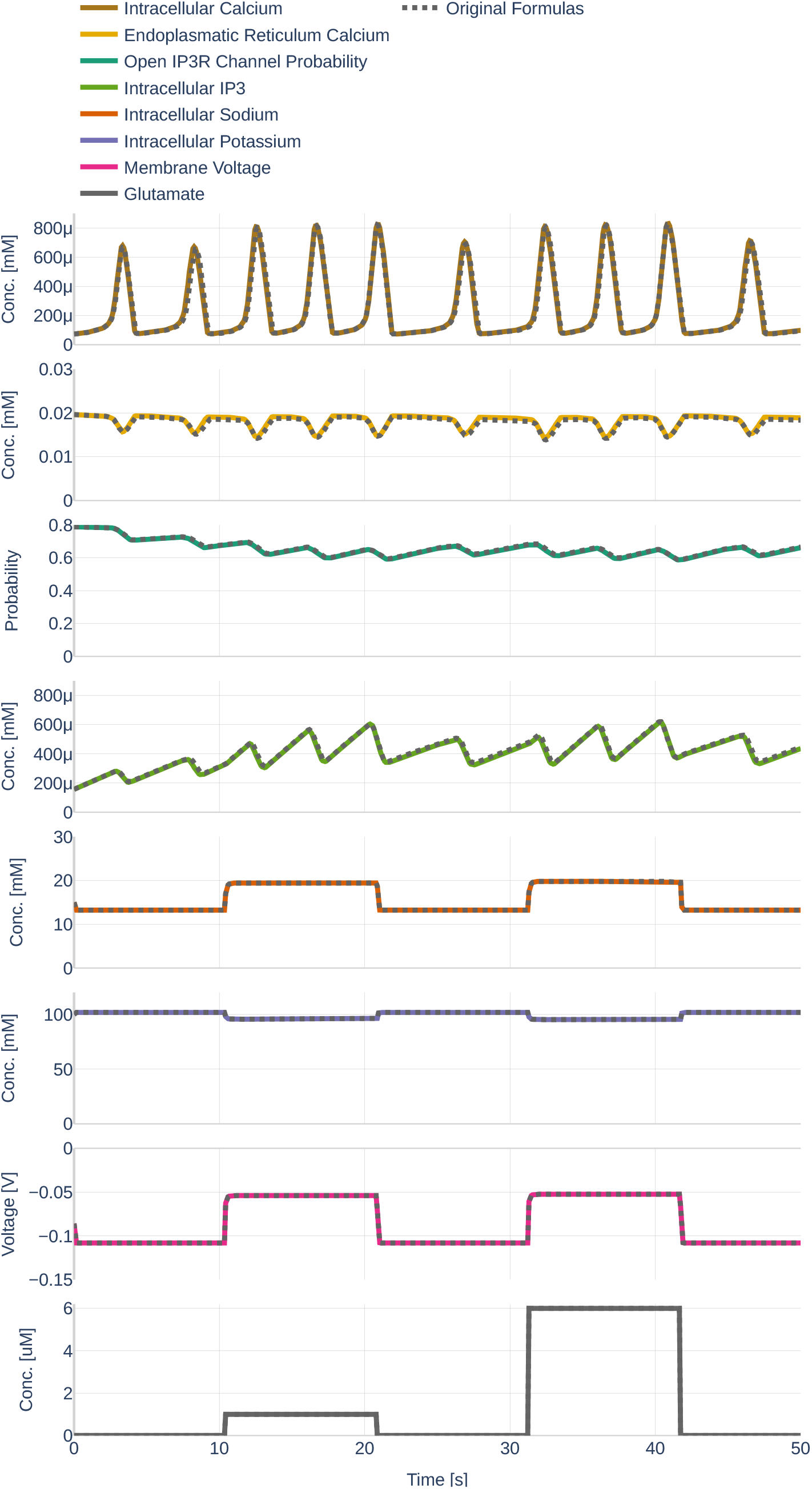
The behavior of the different state variables [*Ca*^2+^]_*i*_, [*Ca*^2+^]_*e*_, *h*, [*IP*_3_]_*i*_, [*Na*^+^]_*i*_, [*K*^+^]_*i*_, *V* over time. Black dashed lines (where visible) indicate the behavior of the state variable before the changes described in Section 3.2.1 were made. Note the differently scaled axes.

As assumed, an increase in [*IP*_3_]_*i*_ is correlated with an increase in the open probability of *IP*_3*R*_ channels. However, the average open probability increases over time while [*IP*_3_]_*i*_ decreases. The presence of a glutamate stimulus results in a higher frequency of *IP*_3_ accumulation- and degradation. While the state variables described so far fluctuate over time, the *V*_*m*_, the [*K*^+^]_*i*_, and [*Na*^+^]_*i*_ only change within the first seconds after a change in glutamate stimulus, therefore appearing like step functions. The reaction of *V*_*m*_ and [*K*^+^]_*i*_ to an increase in glutamate can be described as exponential decay; the reaction of the [*Na*^+^]_*i*_ as exponential saturation.

Figure 11 depicts the different currents of the GluT-driven pathway. The NKA current, the Na^+^- and K^+^ leak current, as well as the GluT current, resemble step functions, similar to the previously observed Na^+^, K^+^, and voltage membrane dynamics. Furthermore, it can be seen that the NCX current is significantly smaller than the other GluT pathways currents. It stands out that the Na^+^ leak current is negative, while the K^+^ leak current is positive, indicating that the K^+^ leak points inward rather than outward as would be expected from the schematics shown in the original paper by Oschmann et al. [2017]. By running the code written by Dr. Oschmann, we observed that the original code suffers from the same problem.

**Figure 11.**
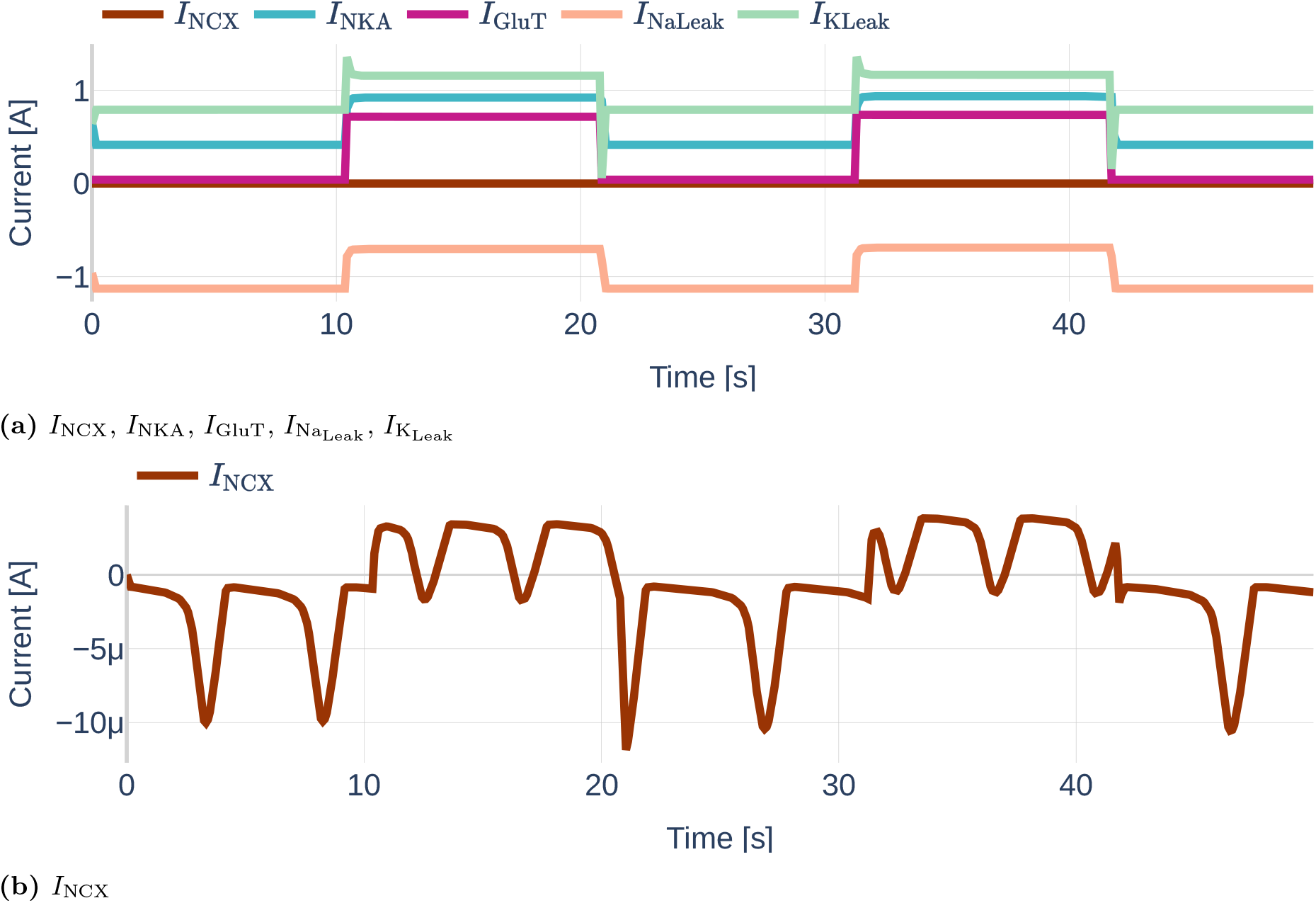
Dynamics of the GluT-pathway related currents *I*_NCX_, *I*_NKA_, 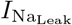, 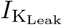 and *I*_GluT_ (a). Due to the different orders of magnitude, *I*_NCX_ is plotted a second time in (b). Note the different scales of the y axes. The used glutamate stimulation is shown in Figure 9.

Similarly, Figure 12 shows the dynamics of the mGluR pathway-driven currents. It can be seen that both, *I*_Serca_ and 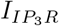 heavily oscillate. Increases and decreases of the SERCA current correlate positively with increases and decreases of the *IP*_3_*R* current. The Ca^2+^ leak current slightly decreases linearly during the raise in SERCA and *IP*_3_*R* current, dips shortly when *I*_Serca_ reaches its maximum and then recovers back to its initial value.

**Figure 12.**
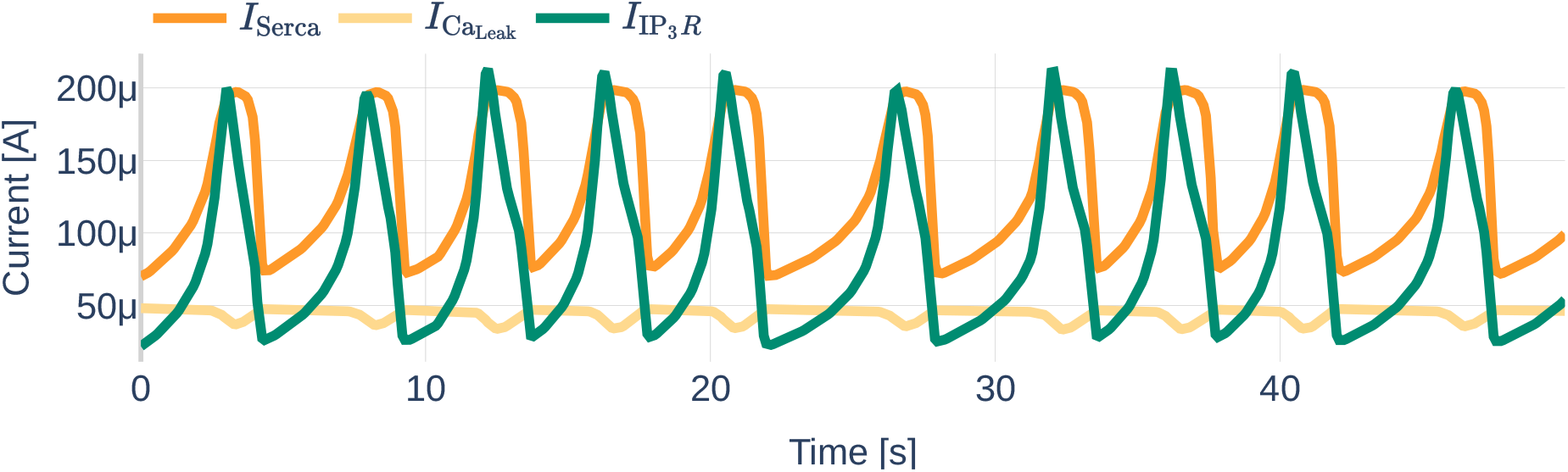
Dynamics of the mGluR-pathway related currents *I*_Serca_, 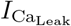 and 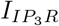. The used glutamate stimulation is shown in Figure 9.

#### 4.1.2 Influence of Conceptual Changes

Second, we studied the influence of the conceptual changes described in Section 3.2.1. The black dotted lines in Figure 10 represent the respective results of the original, unchanged computational model. Other than for the [*Ca*^2+^]_*i*_ and [*Ca*^2+^]_*e*_, the changes are barely visible. This corresponds with the computation of the mean absolute and the mean relative deviation with respect to the original model listed in Table 6. The changes of Equation 15 regarding the computation of 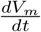 barely affected the membrane voltage. However, adding the valence of Ca^2+^ to 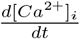 in Equation 16 affected the [*Ca*^2+^]_*i*_ significantly. Correspondingly, significant changes were also observed for [*Ca*^2+^]_*e*_, the intracellular *IP*_3_ concentration and the open probability *h* - although the effect was less pronounced.

**Table 6.** Absolute and relative deviation of the state variables with respect to the computational astrocyte model as described in the paper by Oschmann et al. [2017].

| State Variable | Absolute Deviation | Relative Deviation |
| --- | --- | --- |
| $[Ca^{2+}]_i$ | 1.479e-05 mM | 5.24% |
| $[Ca^{2+}]_e$ | 3.411e-04mM | 1.94% |
| $h$ | 3.502e-03 | 0.538% |
| $[IP_3]_i$ | 7.488e-06 mM | 1.87% |
| $[Na^+]_i$ | 1.084e-03 mM | 0.00711% |
| $[K^+]_i$ | 4.162e-04 mM | 0.000432% |
| $V_m$ | 2.748e-06 V | 0.0038% |

#### 4.1.3 Influence of Different Parameter Sets

In this section, the influence of the different parameter sets described in Section 3.2.3 is examined. The dynamics resulting from the three different parameter sets `Paper`, `Thesis`, and `Default` are shown in Figure 13. While Ca^2+^ levels, *IP*_3_ concentrations, and open probability oscillate heavily in the parameter set `Default`, their behavior is more linear for the parameter sets `Paper`and `Thesis`. The [*Ca*^2+^]_*i*_ mimics a step function that increases whenever a glutamate stimulus is present, thereby behaving similarly to the [*K*^+^]_i_ and [*Na*^+^]_*i*_. During the absence of a glutamate stimulus, the [*Ca*^2+^]_*e*_ decreases linearly, only to linearly increase again during the presence of a stimulus. Increases are more pronounced for the parameter set `Thesis`. The open probability of *IP*_3_*R* channels and the *IP*_3_ concentration show opposite behavior to the [*Ca*^2+^]_*e*_. At the same time, Na^+^ levels, K^+^ levels, and *V*_*m*_ are barely affected by the change in the parameter set.

**Figure 13.**
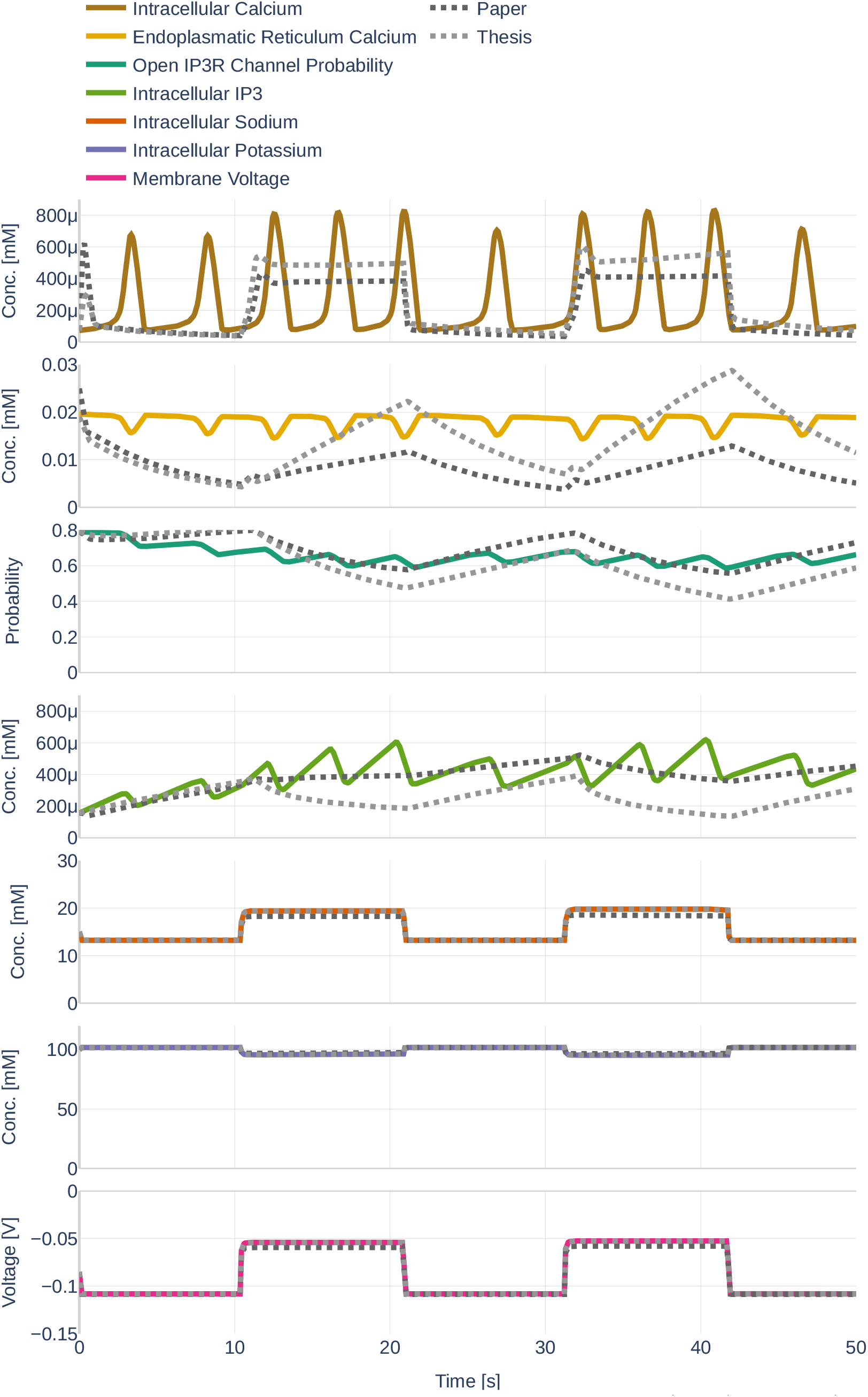
Dynamics resulting from the different parameter sets `Paper` (black), `Thesis` (gray) and `Default` (colored).The used glutamate stimulation is shown in Figure 9.

### 4.2 Learning the Dynamics and their Gradients

Before starting with the parameter inference experiments, we ensured that the network size (number of layers and number of nodes per layer) is large enough to represent the dynamics of all seven ODEs. To that end, we trained the network on the data set `Parameter Set` and assumed that all data can be observed and that all parameters are known. The learned data can be seen in Figure 14. The network learns the dynamics (dotted black line) perfectly in comparison to the underlying dynamics. Figure 15 then depicts both, the gradient of the learned network function 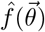 and the gradient returned by the ODEs if the output of the neural network is fed into the computational model. The colored lines indicate the gradients computed during the initial simulation. It is apparent that the network is successful at learning the gradients for 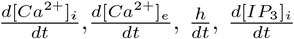. However, large errors occur for the ODEs computed by the computational model for 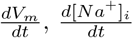 and 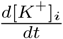.

**Figure 14.**
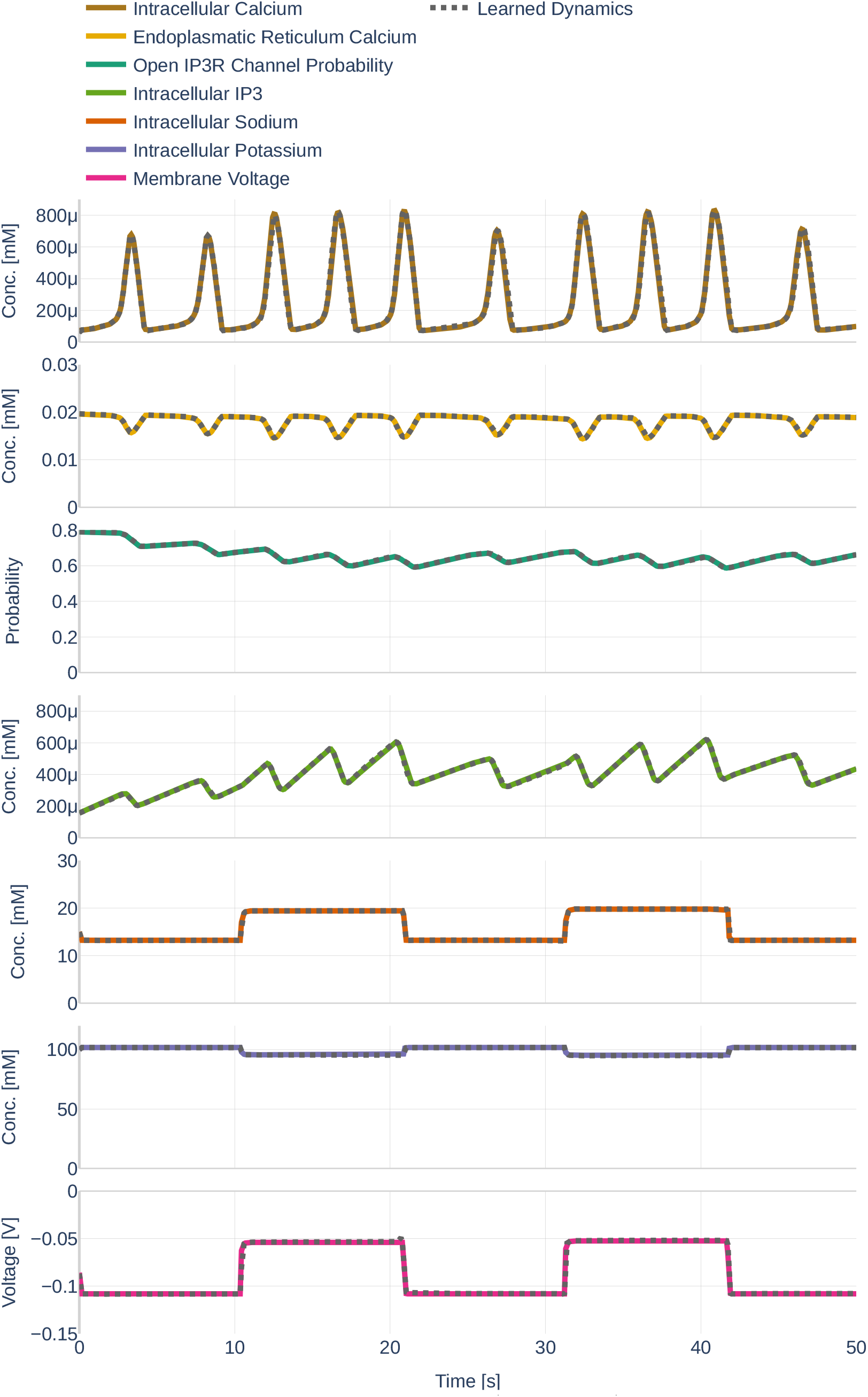
Dynamics as learned by the neural network (dotted lines) in comparison to the dynamics outputted by the Oschmann et al. [model (colored lines). The used glutamate stimulation is shown in Figure 9.

**Figure 15.**
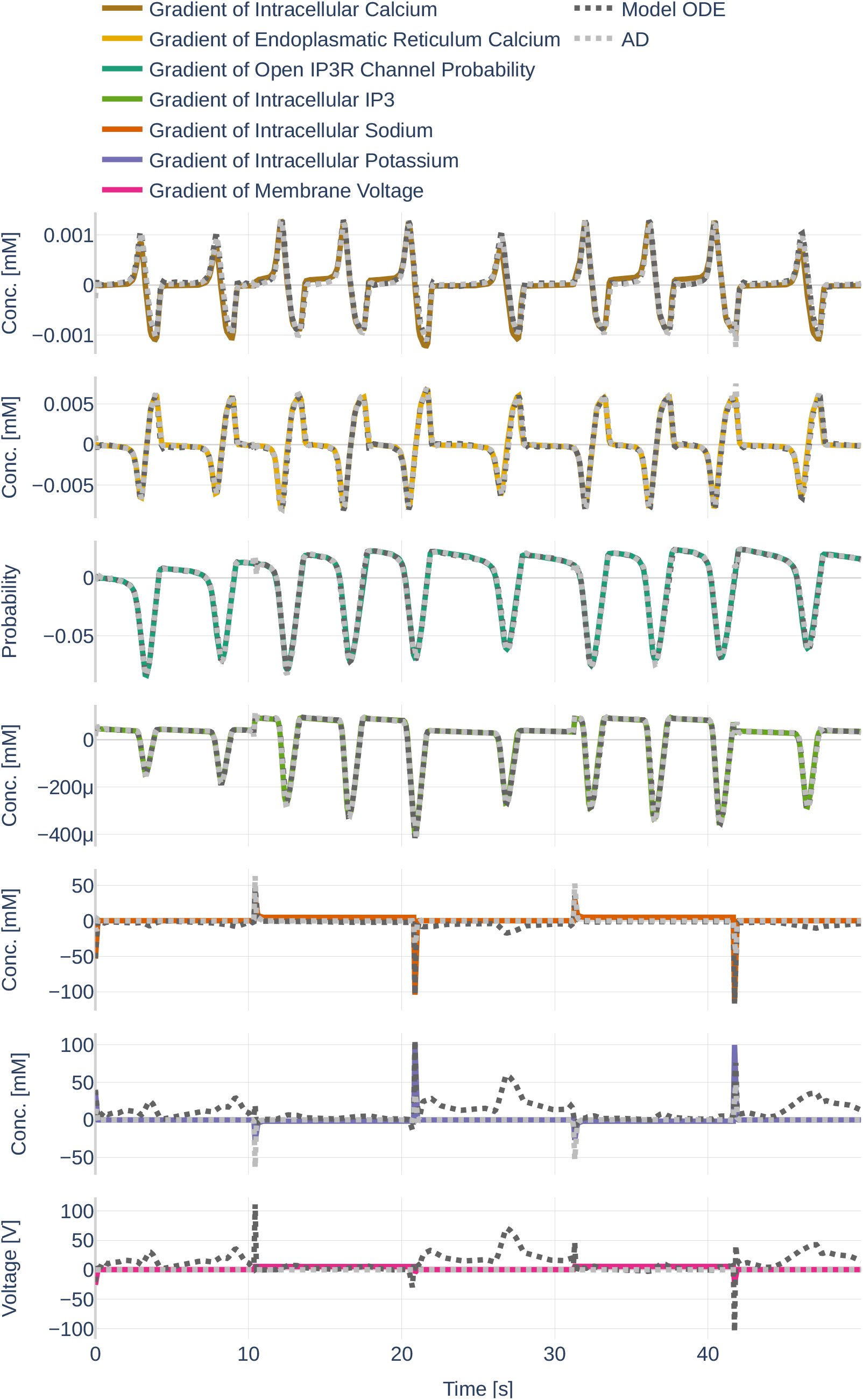
Gradients as they are learned by the neural network (light gray, dotted lines). Furthermore, it shows the gradients outputted by the Oschmann et al. [model if it is fed the neural network output as input (dark gray, dotted lines) and the gradients as they originally occur (colored lines). The used glutamate stimulation is shown in Figure 9.

### 4.3 Parameter Inference and Influence of Changes made to the Original Algorithm by Yazdani et al. [2020]

In this section, we show the results of three different parameter inference experiments.

#### 4.3.1 Precision (Repeatability)

As part of the first parameter inference experiments, we studied the stability of the algorithm. To that end, we run the algorithm with fixed configurations six times and observed if the network infers the same parameter each time. The network was trained on the data set `Parameter Study` and all but the dynamics of 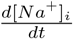 and 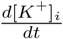 were observed. For each run, we inferred the parameter *K*_NKA*mN*_ (Table 3). To ensure that the inference result is start point independent, we started three times with the assumption *K*_NKA*mN*_ (*t* = 0) = 1 and three times with the assumption *K*_NKA*mN*_ (*t* = 0) = 20. The results can be seen in Figure 16. Other than `Repetition 4`, each run inferred a value around *K*_NKA*mN*_ = 8*mM*, which corresponds to an accuracy of 80%. The exact inferred values are listed in Table 7. `Repetition 4` shows a significant drop in accuracy between epoch 45000 and 50000. However, the accuracy starts raising again afterwards. It is therefore likely that the network would achieve the same accuracy as the other runs after more iterations.

**Figure 16.**
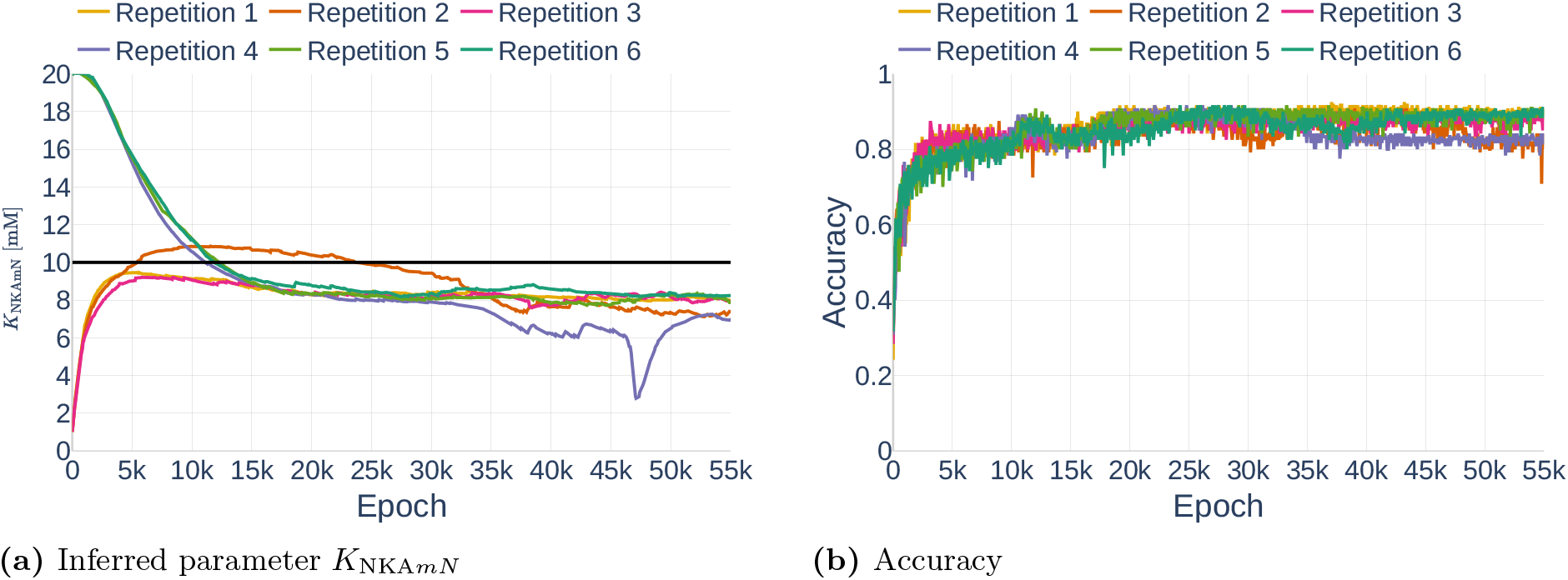
Inferred parameter for *K*_NKA*mN*_ (a) and the accuracy 𝒜_all_ (b) over different epochs. The experiment is performed with a gradient clipping value of *c* = 10 and no learning rate reduction. The black line in (a) indicates the original parameter value.

**Table 7.**
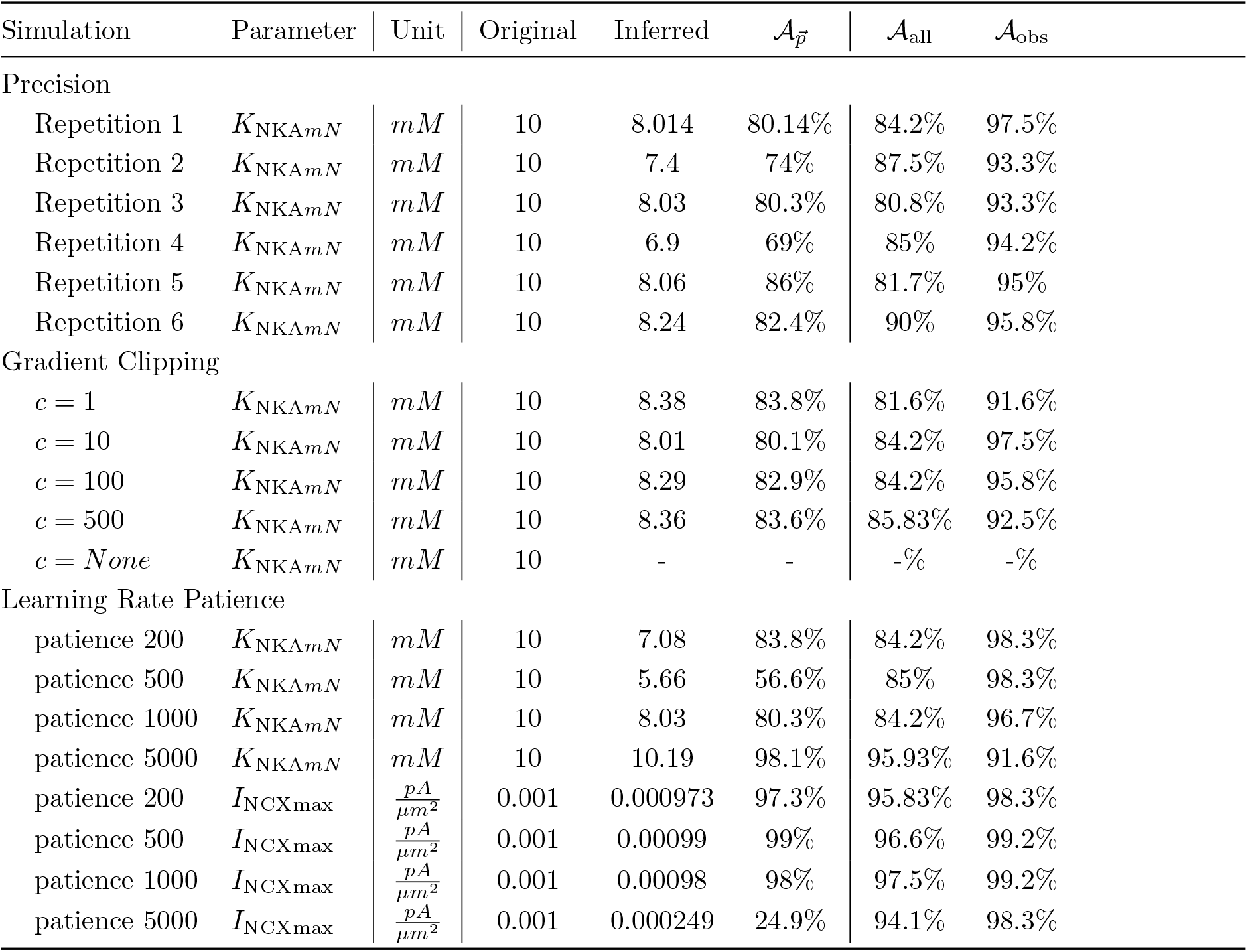
Inferred parameter values and accuracies for the different parameter inference experiments.

| Simulation | Parameter | Unit | Original | Inferred | $\mathcal{A}_{\vec{p}}$ | $\mathcal{A}_{\text{all}}$ | $\mathcal{A}_{\text{obs}}$ |
| --- | --- | --- | --- | --- | --- | --- | --- |
| Precision |  |  |  |  |  |  |  |
| Repetition 1 | $K_{\text{NKAmN}}$ | $mM$ | 10 | 8.014 | 80.14% | 84.2% | 97.5% |
| Repetition 2 | $K_{\text{NKAmN}}$ | $mM$ | 10 | 7.4 | 74% | 87.5% | 93.3% |
| Repetition 3 | $K_{\text{NKAmN}}$ | $mM$ | 10 | 8.03 | 80.3% | 80.8% | 93.3% |
| Repetition 4 | $K_{\text{NKAmN}}$ | $mM$ | 10 | 6.9 | 69% | 85% | 94.2% |
| Repetition 5 | $K_{\text{NKAmN}}$ | $mM$ | 10 | 8.06 | 86% | 81.7% | 95% |
| Repetition 6 | $K_{\text{NKAmN}}$ | $mM$ | 10 | 8.24 | 82.4% | 90% | 95.8% |
| Gradient Clipping |  |  |  |  |  |  |  |
| $c = 1$ | $K_{\text{NKAmN}}$ | $mM$ | 10 | 8.38 | 83.8% | 81.6% | 91.6% |
| $c = 10$ | $K_{\text{NKAmN}}$ | $mM$ | 10 | 8.01 | 80.1% | 84.2% | 97.5% |
| $c = 100$ | $K_{\text{NKAmN}}$ | $mM$ | 10 | 8.29 | 82.9% | 84.2% | 95.8% |
| $c = 500$ | $K_{\text{NKAmN}}$ | $mM$ | 10 | 8.36 | 83.6% | 85.83% | 92.5% |
| $c = \text{None}$ | $K_{\text{NKAmN}}$ | $mM$ | 10 | - | - | -% | -% |
| Learning Rate Patience |  |  |  |  |  |  |  |
| patience 200 | $K_{\text{NKAmN}}$ | $mM$ | 10 | 7.08 | 83.8% | 84.2% | 98.3% |
| patience 500 | $K_{\text{NKAmN}}$ | $mM$ | 10 | 5.66 | 56.6% | 85% | 98.3% |
| patience 1000 | $K_{\text{NKAmN}}$ | $mM$ | 10 | 8.03 | 80.3% | 84.2% | 96.7% |
| patience 5000 | $K_{\text{NKAmN}}$ | $mM$ | 10 | 10.19 | 98.1% | 95.93% | 91.6% |
| patience 200 | $I_{\text{NCXmax}}$ | $\frac{pA}{\mu m^2}$ | 0.001 | 0.000973 | 97.3% | 95.83% | 98.3% |
| patience 500 | $I_{\text{NCXmax}}$ | $\frac{pA}{\mu m^2}$ | 0.001 | 0.00099 | 99% | 96.6% | 99.2% |
| patience 1000 | $I_{\text{NCXmax}}$ | $\frac{pA}{\mu m^2}$ | 0.001 | 0.00098 | 98% | 97.5% | 99.2% |
| patience 5000 | $I_{\text{NCXmax}}$ | $\frac{pA}{\mu m^2}$ | 0.001 | 0.000249 | 24.9% | 94.1% | 98.3% |

#### 4.3.2 Gradient Clipping

Next, we studied the effect of gradient clipping values. Using the data set `Parameter Study` and assuming all but the dynamics of 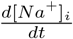 and 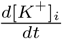 observed, we run the algorithm for the gradient clipping values *c* = 1, *c* = 10, *c* = 100, *c* = 500 and *c* = *None* (indicating no gradient clipping). The results are shown in Figure 17 and the exact inferred values are listed in Table 7. The simulation for *c* = *None* started showing inconsistent behavior after epoch 20000 and finally predicted NaN-Values shortly before epoch 30000, therefore being unable to make further predictions or improvements. In general, it can be seen that the network experienced large fluctuations for *c* = 100 but made stable progress for *c* = 10 and *c* = 1. However, the plot of the accuracy 𝒜_all_ shows that the learning process for *c* = 1 was slower than the progress for *c* = 10. As before, all simulations with a fixed gradient clipping value inferred approximately a value of *K*_NKA*mN*_ = 8*mM*.

**Figure 17.**
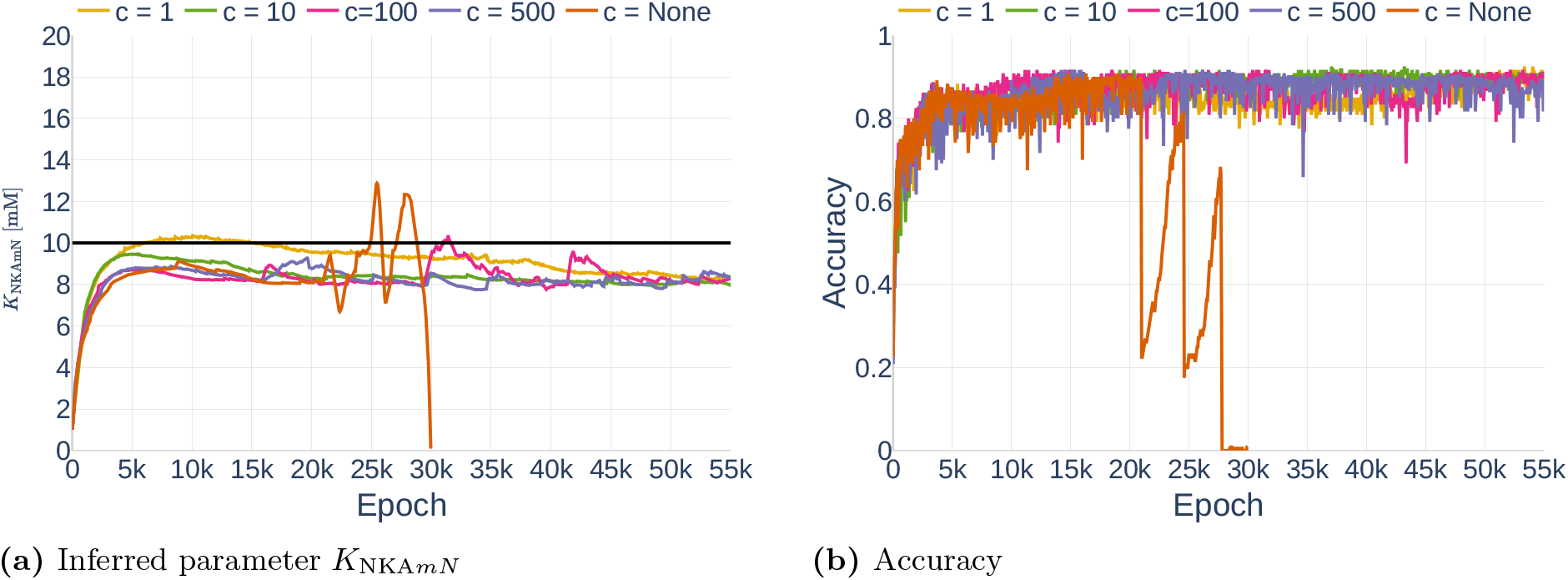
Effect of different gradient clipping values *c* on the inferred parameter (a) and the accuracy (b). The black line in (a) indicates the original parameter value.

#### 4.3.3 Unstable Learning Process and the Problem with Patience

As explained in Section 3.3.5, the property `patience` of a learning reduction algorithm describes how long it takes before the learning rate gets reduced if the loss does not decrease. In this section, we show the problem with setting the `patience` correctly. Figure 18 shows the inference of parameters *K*_NKA*mN*_ and *I*_NCX*max*_. All simulations were performed with the data set `Parameter Study` and with all but 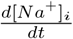 and 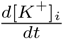 assumed observed. The gradient clipping value was set to *c* = 10. The inference of *I*_NCX*max*_ is very stable for a `patience` between 200 and 1000. However, the network has problems inferring the correct parameter with a higher `patience` of 5000, starting to deviate from a good inference of approximately the correct value after epoch 25000. In comparison, a `patience` of 200 is too small for the inference *K*_NKA*mN*_, leading the network to stop learning too early. The inferred values are listed in Table 7.

**Figure 18.**
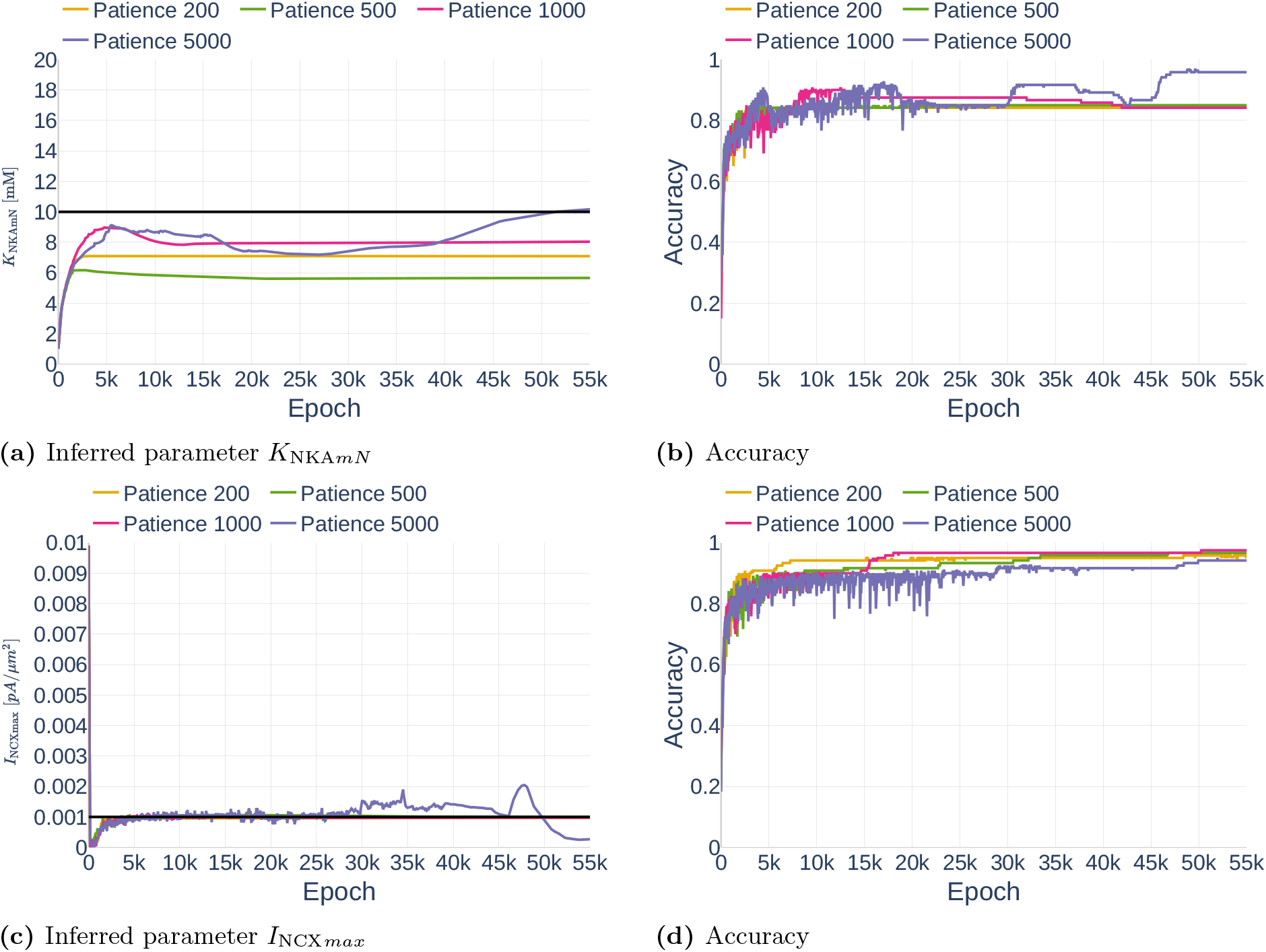
Effect of different amounts of `patience` regarding the learning rate reduction for *K*_NKA*mN*_ (a and b) and *I*_NCX*max*_ (c and d). The black line in (a) and (c) indicate the original parameter value.

## 5 Methods, Part 2

In this section, we describe several methods that aim at stabilizing the inference of parameters in the Oschmann et al. [model. In Section 6.1, we explain a change to the Oschmann et al. [model that aims at stabilizing the *V*_*m*_ problem shown in Section 4.2. Based on a paper by Wang et al. [2020], we show methods to improve gradient pathologies during the inference process in Section 6.2. Last, Section 6.3 proposes the addition of control inputs to the neural network as was originally done by Antonelo et al. [2021].

### 5.1 Adapted Leak Currents and their subsequent Changes in Neural Network Parameters

As was observed in Section 4.2, the gradients of [*Na*^+^]_*i*_, [*K*^+^]_*i*_, and *V*_*m*_ returned by the computational Oschmann et al. [model are extremely sensitive to small errors in the input states. In part, this is due to the way the leak currents are computed. The original model computes the leak currents as

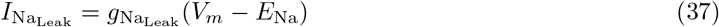

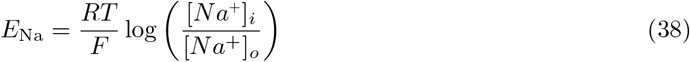

and

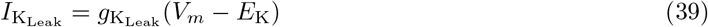

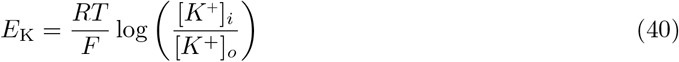

where *F* is the Faraday constant, *R* is the molar gas constant and *T* is the current temperature. This way of computing *E*_*K*_ and *E*_Na_ introduces a high level of sensitivity to the computations of the leak current, especially as the outer- and inner concentrations of both K^+^ and Na^+^ are dependent on the amount of [*Na*^+^]_*i*_ and [*K*^+^]_*i*_. To decouple this sensitivity, we replaced the dynamic computation of *E*_Na_ and *E*_*K*_ with constants, as is regularly done in other computational astrocyte models [Farr and David, 2011, Flanagan et al., 2018]. The used constants are equal to the known reversal potentials of Na^+^ and K^+^ and are listed in Table 10.

**Table 8.**
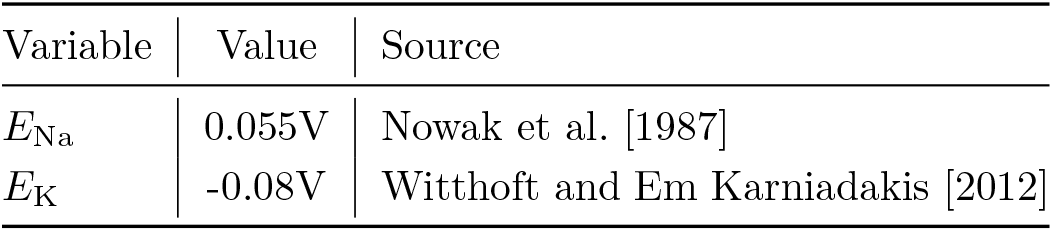
The used reversal potentials for *Na*^+^ and *K*^+^ and their sources

**Table 9.**
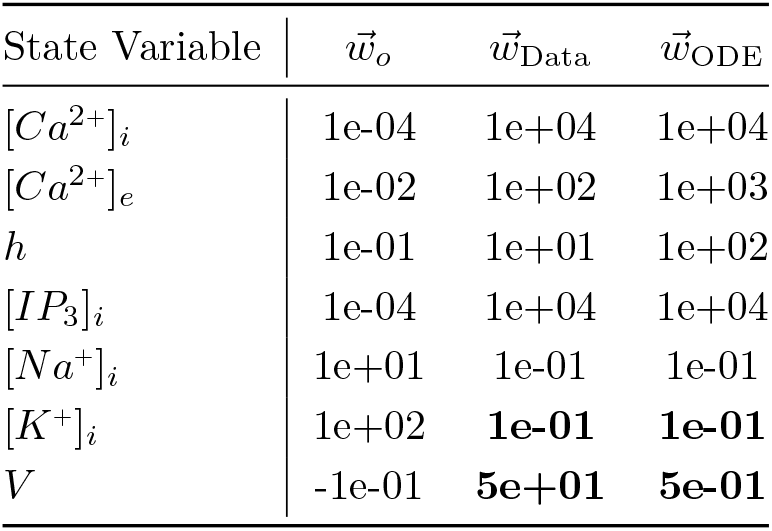
This table lists the state variable related weights used for the deep learning algorithm with new leak computation. Values that changed in comparison to the previous weights (Table 2) are indicated in bold.

**Table 10.**
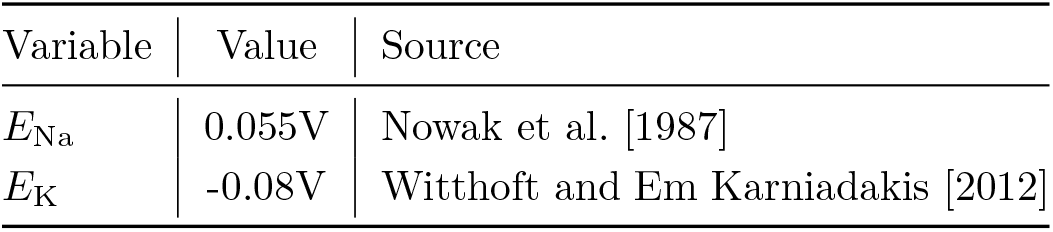
The used reversal potentials for *Na*^+^ and *K*^+^ and their sources

While the new leak computation does not change the general behavior of the simulation, it does change the order of magnitude of the computed gradients. The changed gradients require the usage of adapted weights for the parameter inference algorithm. These weights are listed in Table 11.

**Table 11.**
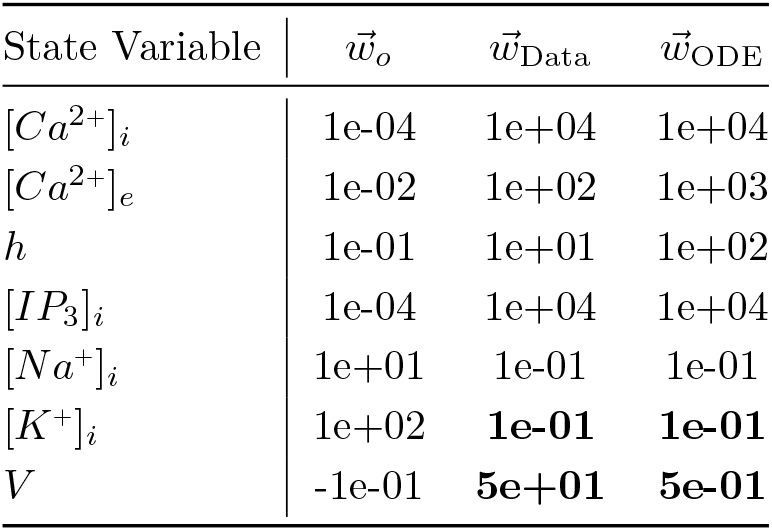
This table lists the state variable related weights used for the deep learning algorithm with new leak computation. Values that changed in comparison to the previous weights (Table 2) are indicated in bold.

### 5.2 Gradient Pathologies in PINNs

While we trained my neural network with the configurations described in Section 3.3, it became obvious that the training process was not as stable and fast as expected. Wang et al. [2020] discovered and addressed one major mode of failure in PINNs. According to them, numerical stiffness might lead to unbalanced learning gradients during the back-propagation step in model training. They solved the problems in two ways. First, they suggested an algorithm that outbalances different loss terms. Second, they changed the model architecture to include a transformer network. In the following sections, we shortly describe their propositions and then explain how we adapted them for my model.

#### 5.2.1 Learning Rate Annealing

##### Implementation

To give different amounts of importance to different loss terms, loss terms are usually weighted. Assuming the loss functions consist of an ODE loss ℒ_*ODE*_ and *M* different data loss terms, such as different kinds of measurements or boundary conditions, the total loss can be written as

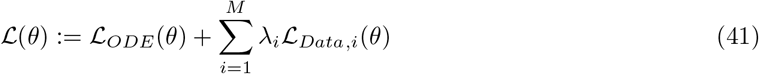

where λ_*i*_ is the weight of ℒ_*Data,i*_. Based on the optimization method `Adam` [Kingma and Ba, 2014] explained earlier, the authors suggested scaling the weights according to the ratio between the largest and average gradient of the different loss terms. Let

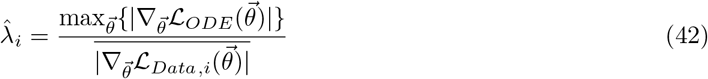

<u>where</u> 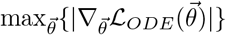 is the largest absolute parameter gradient of the ODE loss and 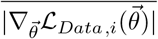 denotes the mean absolute parameter gradient of the different data loss terms. Due to the possibly high variance of 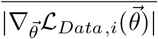, it was suggested to not directly use 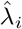 for weighting but rather to compute a running average using the equation

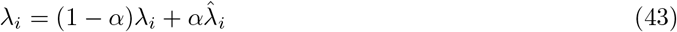

with *α* ∈ [0.5, 0.9]. Assuming `SGD` optimization (Equation 12) is used, the optimization step becomes

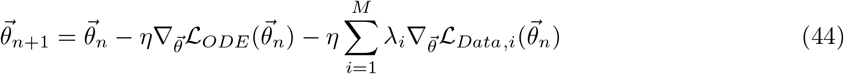

where *n* stands for the *n*-th iteration and ∇ is the learning rate. Wang et al. [2020] suggest to use a learning rate of *η* = 1e^−3^.

##### Adaption for Parameter Inference Deep Learning

In contrast to the examples by [Wang et al., 2020], the computational model at hand [Oschmann et al., 2017] consists of multiple ODEs. Furthermore, observations usually only exist for a subset of the given ODEs. This leads to the question of how the learning rate annealing algorithm should be adapted for my model. We tested three different strategies (`A`, `B`, `C`) further described below. To reduce the computational effort of computing 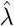, we only performed an update step every 50th epoch. The different strategies are visualized in Figure 23.

**Figure 19.**
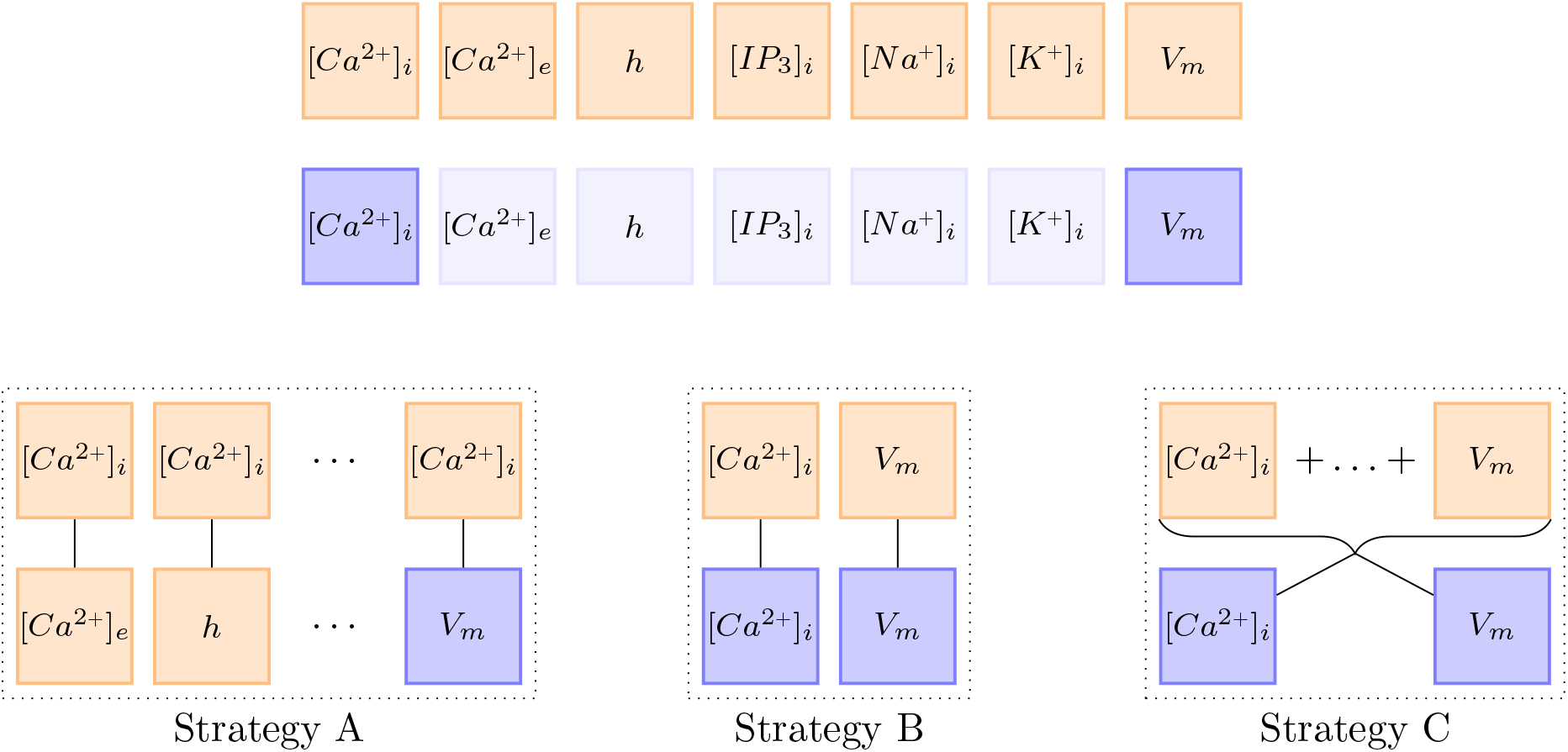
This figure visualizes the three different λ update strategies. Orange boxes stand for the gradient with respect to the network parameters of the ODE loss 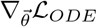. Blue boxes stand for the gradient with respect to the network parameters of the data loss 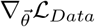. Almost transparent blue boxes indicate that the respective data was not observed and is therefore not considered in the loss functions.

**Figure 20.**
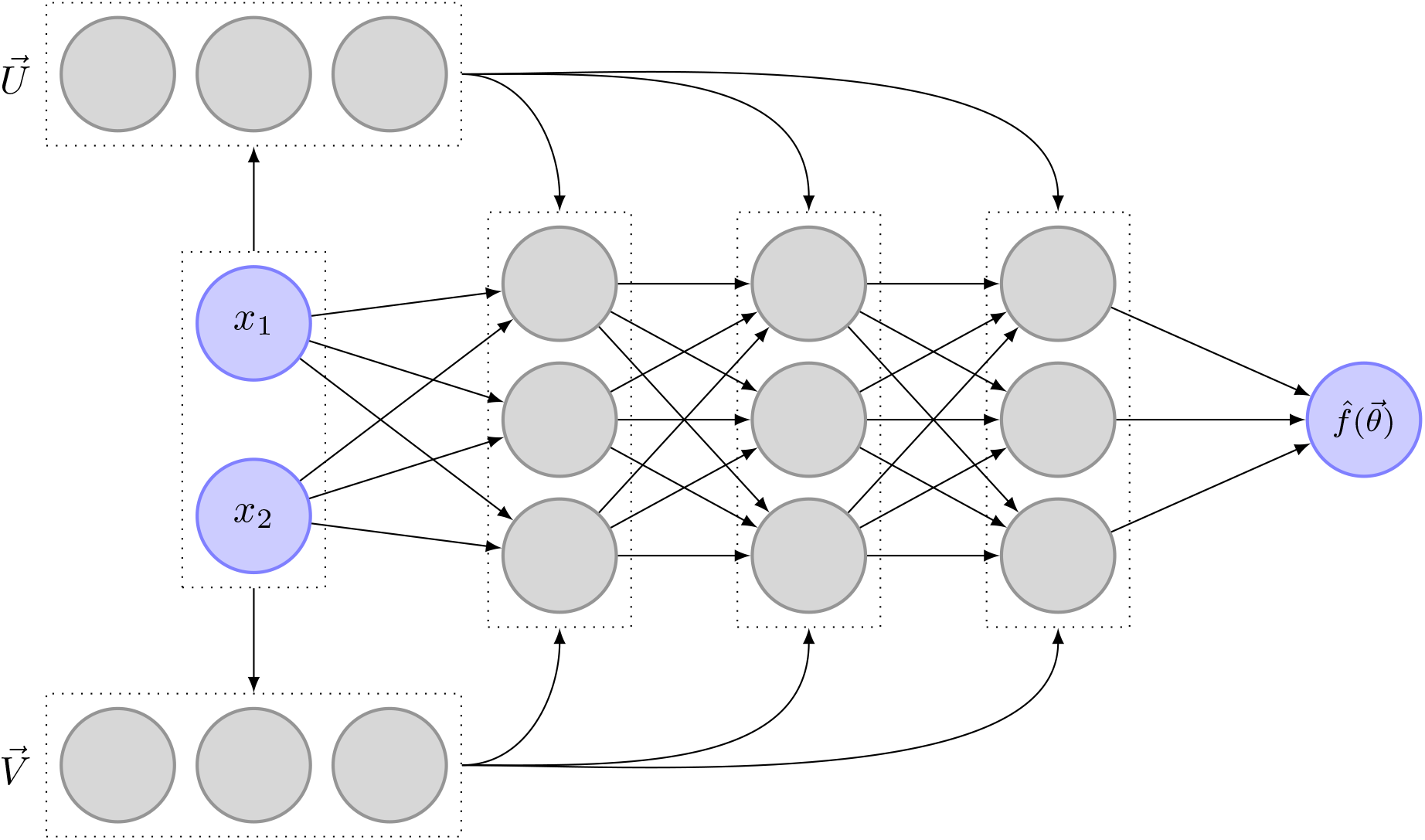
This figure shows the extension of a transitional neural network with two additional, fully connected layers 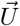 and 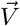. This addition is based on the idea of transformer networks and was adapted to PINNs in a recent paper by Wang et al. [2020]

**Figure 21.**
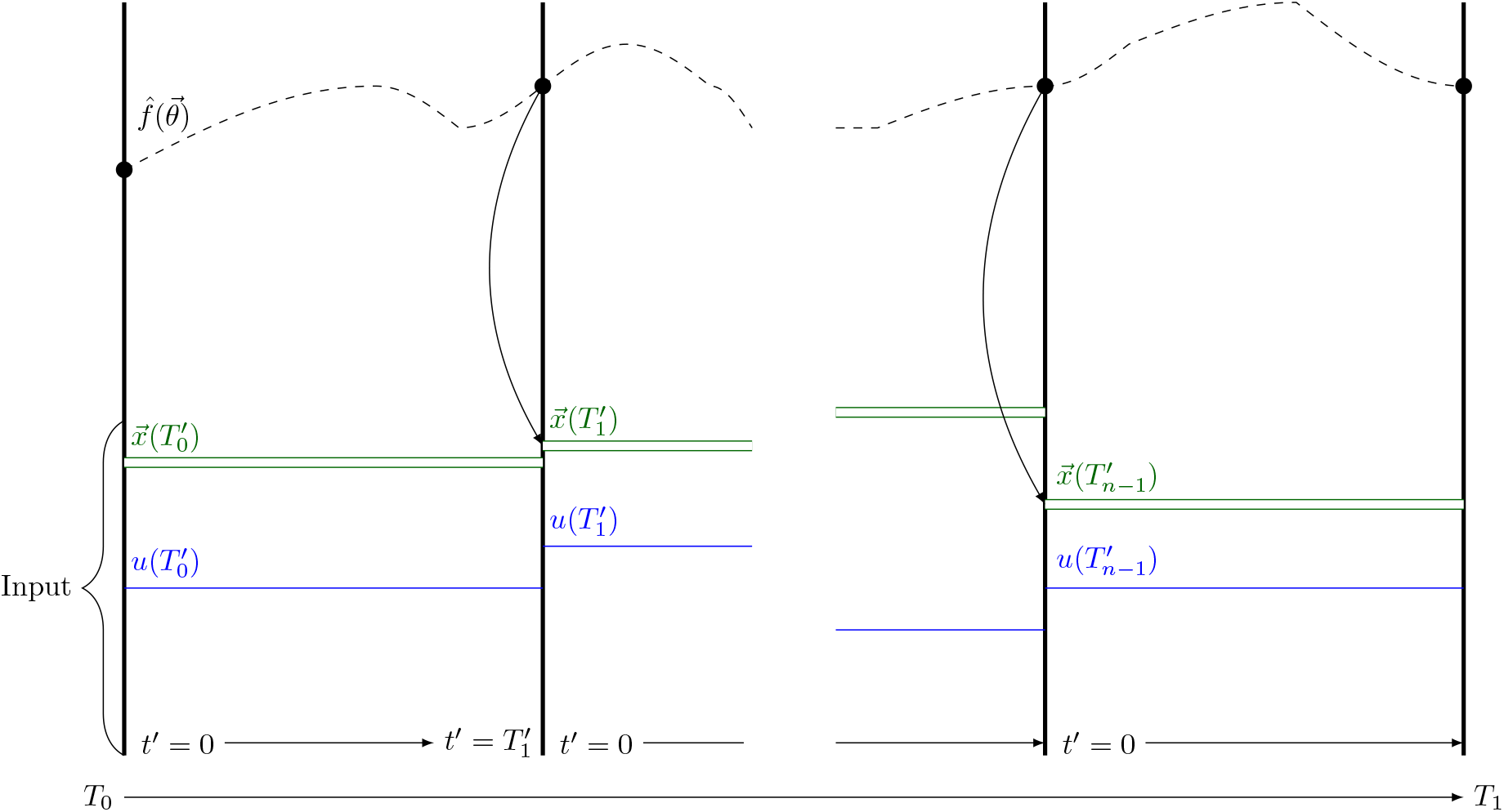
Schematic of data propagation in a PINC based on a figure in the original paper by Antonelo et al. [2021]. *u* is the control input of the different intervals 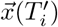 represents the corresponding initial conditions. 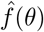 is the function learned by the neural network.

**Figure 22.**
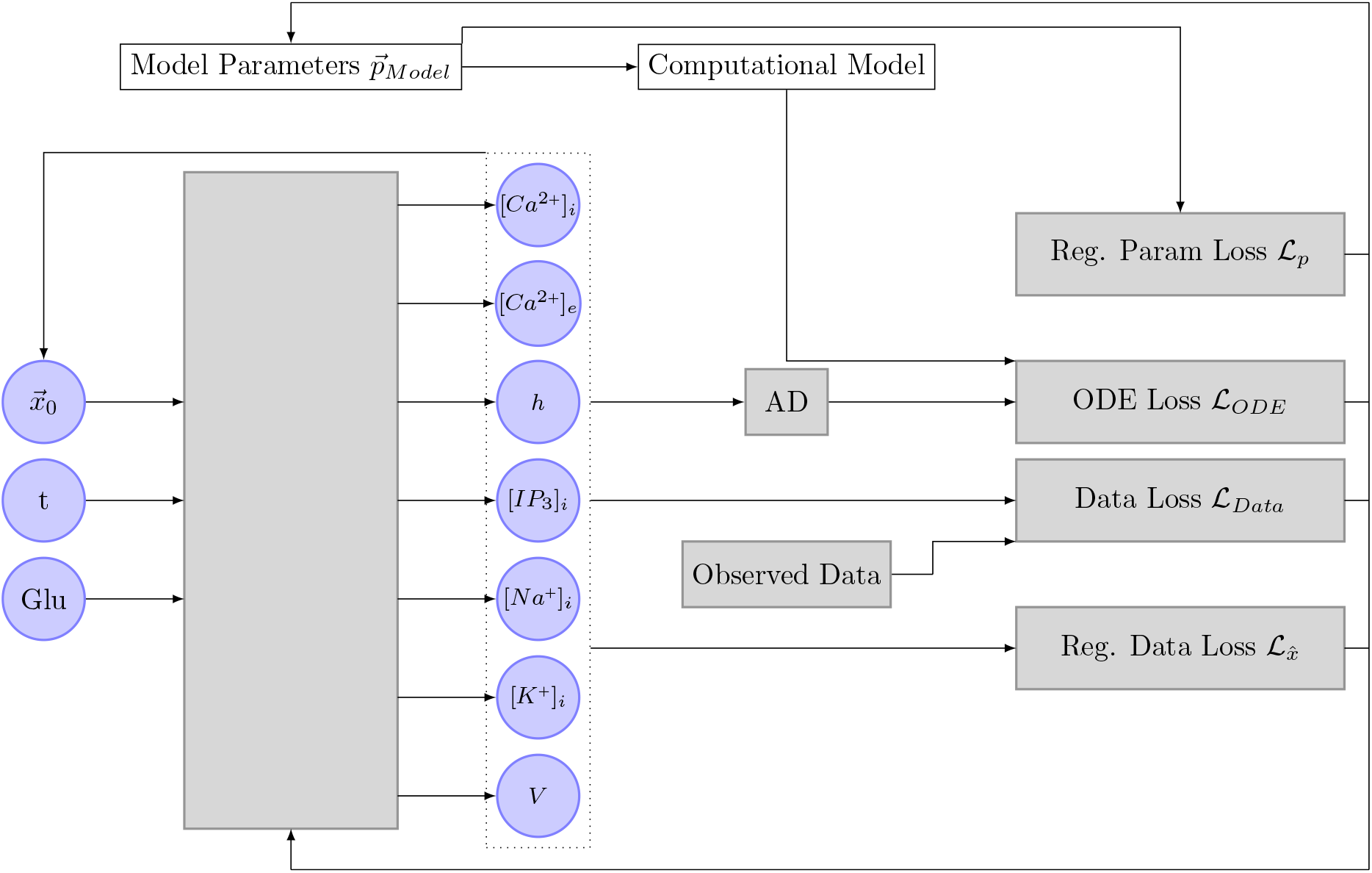
This schematic shows the adaptation of the algorithm by Antonelo et al. [2021] to the Oschmann et al. [model and in combination with the deep learning algorithm initially developed by Yazdani et al. [2020]. The input of the neural network is expanded with a control input (`Glu`) and initial conditions 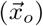. The initial conditions of each interval are predicted by the neural network itself and replace the previously used concept of ℒ_*Aux*_. AD stands for automatic differentiation.

**Figure 23.**
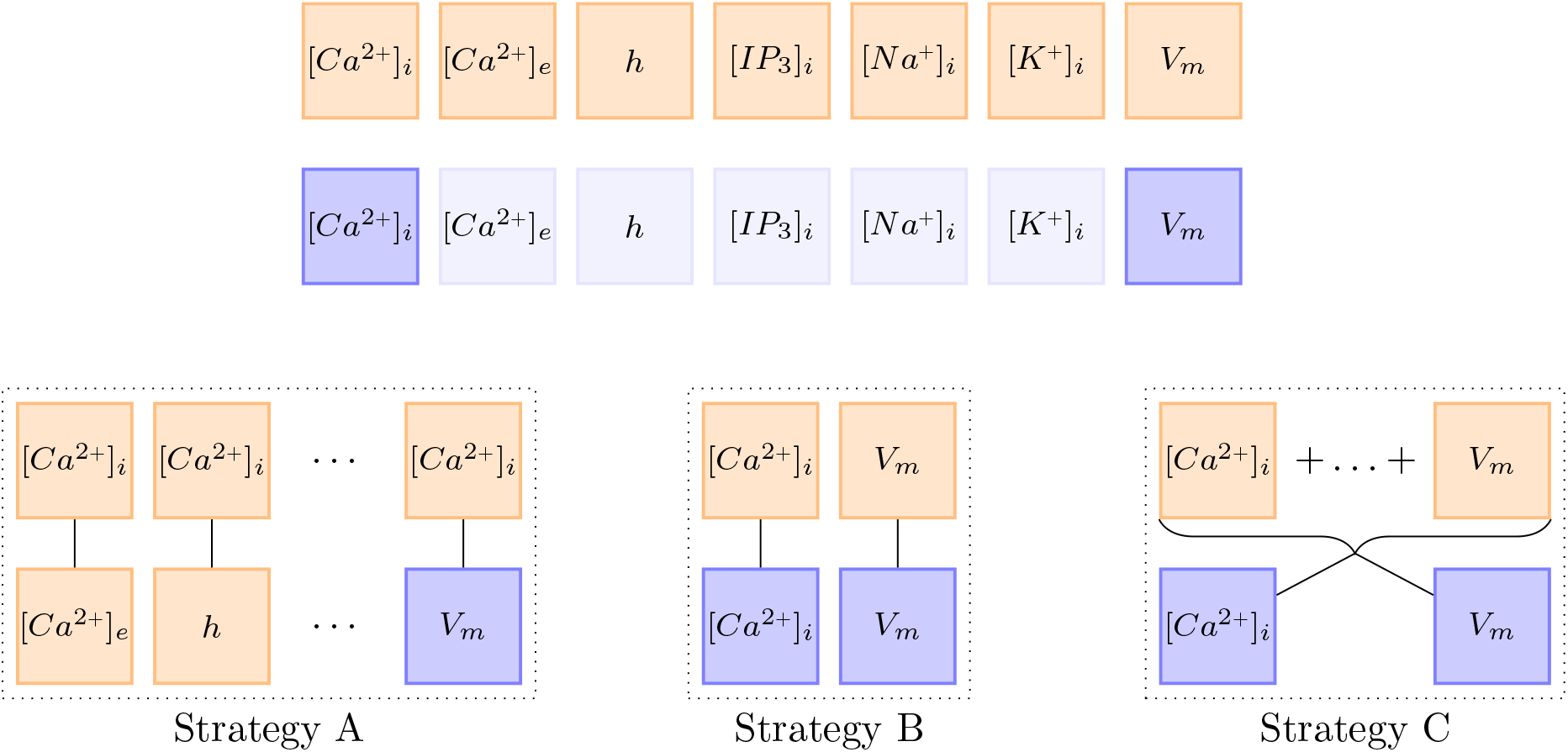
This figure visualizes the three different *λ* update strategies. Orange boxes stand for the gradient with respect to the network parameters of the ODE loss 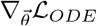. Blue boxes stand for the gradient with respect to the network parameters of the data loss 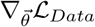. Almost transparent blue boxes indicate that the respective data was not observed and is therefore not considered in the loss functions.

##### Strategy A

First, we matched the weight of the first ODE loss gradient 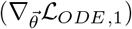 against all other loss gradients, both ODE loss and data loss, separately. This idea was motivated by the observation that it is not about balancing the ODE loss with the data loss, but about balancing all terms with each other. By setting 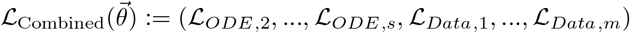 the total loss can be defined as

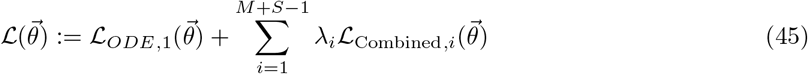

Then, 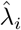 becomes

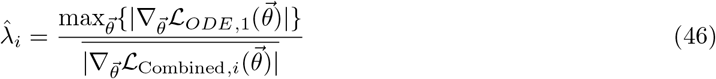

and is used in combination with the moving average Equation 63 to compute λ_*i*_.

##### Strategy B

For my second strategy, we assumed that the different ODEs are already balanced out well enough through the loss weighting described in Section 3.3.4. Therefore, the weighting only has to be adjusted between an ODE loss and its respective data loss. If an ODE does not have a counterpart, we do not change the weighting. Assuming no regularization and auxiliary losses, the total loss can be written in the form 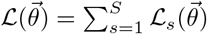 with

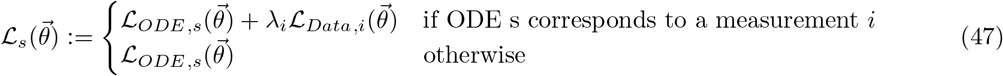

The weights are then computed using

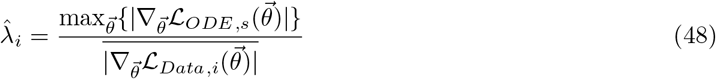

together with the moving average Equation 63.

##### Strategy C

For my last strategy, we made the same assumption as for Strategy `B`, but rather than balancing each ODE against its counterpart, we took the ratio between the largest gradient of the sum of all ODE losses and the mean gradient of the different data losses. In this form, the total loss is written as

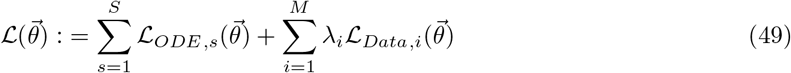

and the temporary weight becomes

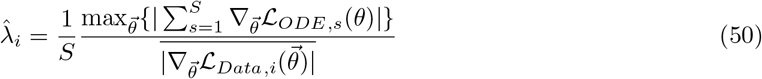

#### 5.2.2 Improved Fully Connected Architecture

The second improvement by Wang et al. [2020] concerned the architecture of the neural network itself and was based on the idea of a Transformer [Vaswani et al., 2017]. Transformers are often used in natural language processing or sequence transduction tasks and offer an alternative to the more commonly known recurrent or convolutional neural networks. Broadly speaking, a transformer considers possible multiplicative connections between different input nodes and strengthens the influence of input nodes on later network layers.

In the context of their paper, Wang et al. [2020] adapted the idea of transformer networks to PINNs by adding two additional network layers 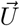 and 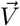. Just as the first fully connected neural network layer, 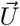 and 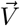 are directly connected to the input layer. They consist of the same number of nodes as all the other network layers. In form of equations, 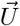 and 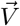 are defined through

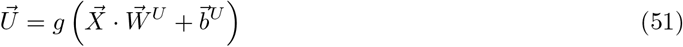

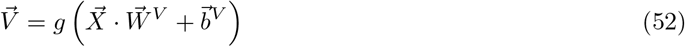

where *g* is the activation function, 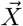 the input layer, 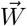 and 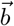 are the layers parameter. To enhance the network’s performance, 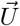 and 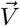 are multiplied component-wise to the output of the normal network layers described in Equation 10. The forward propagation equations, therefore, change to

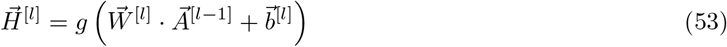

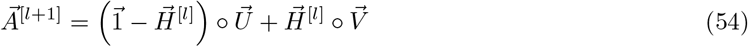

where ° denotes component-wise multiplication. Note that this change does not affect Equation 11 for the output layer of the neural network. Figure 24 shows the addition of 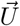 and 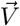 to a fully connected neural network. In the context of this manuscript, we followed the original implementation by Wang et al. [2020] exactly.

**Figure 24.**
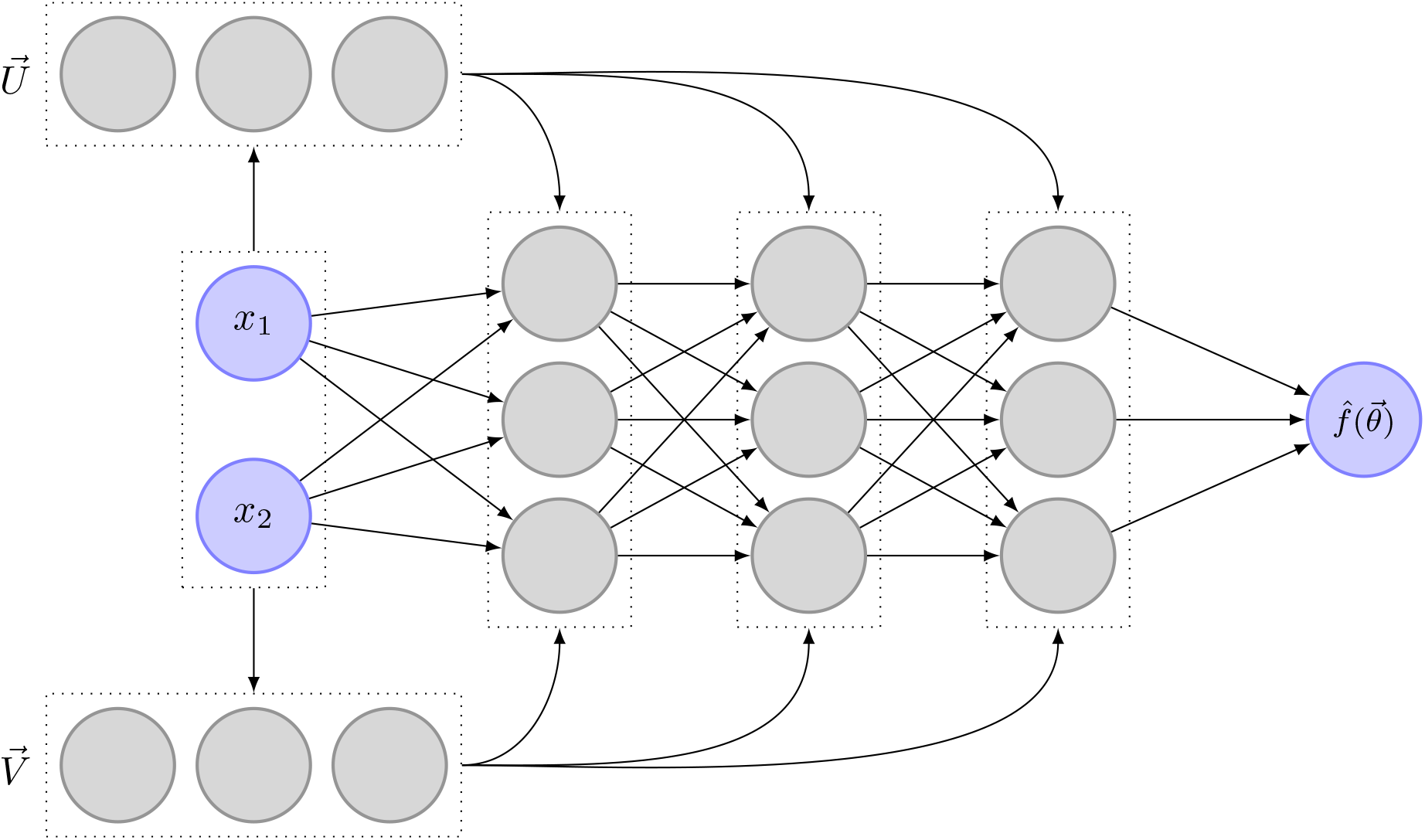
This figure shows the extension of a transitional neural network with two additional, fully connected layers 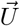 and 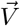. This addition is based on the idea of transformer networks and was adapted to PINNs in a recent paper by Wang et al. [2020]

### 5.3 Control Input

In the context of ODEs, PINNs attempt to learn the relationship between a continuous time input t and several state variables 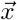. One of the major drawbacks of this method is that external events, such as a glutamate release by a neighboring neuron, can not be taken into account. Therefore, the glutamate level has either to be known or inferred at every point in time. While this might be possible under some preconditions, it is not feasible and further prohibits the use of multiple, different measurement sets to train one specific model.

The same problem is often faced in the context of control theory. While processes in for example the oil, gas, or robotics industry can often be modeled through differential equations, they usually have some dependence on external control inputs. To counteract this problem, Antonelo et al. [2021] recently proposed an adapted PINN algorithm that allows for control inputs. The concept is called *Physics-Informed Neural Nets-based Control* (PINC) and will be detailed further in the next Section 6.3.1. Section 6.3.2 then details how we adapted and implemented the concept of PINC to further improve on the parameter inference algorithm proposed by Yazdani et al. [2020] implemented in the context of this manuscript.

#### 5.3.1 Original Implementation

Inspired by multiple shooting and collocation methods, Antonelo et al. [2021] changed the original PINN algorithm [Raissi et al., 2017] in two significant ways. The first change is concerned with the input time *t*. Rather than attempting to learn how the state variables change over the whole time horizon, they suggested letting the network learn how the state variables have behaved since the last change in control input *u*(*t*). To that end, they subdivided the time interval [*T*_0_, *T*_1_] into multiple, smaller subintervals.

Assuming the control input is given by a piecewise constant function *u*(*t*), they split the time intervals at the points of discontinuity 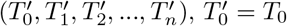 and 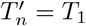, of *u*(*t*). Then, the input to the neural network is changed from *t* to *t*^*′*^ where *t*^*′*^ indicates how much time has passed since the beginning of the current subinterval.

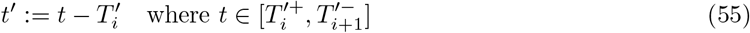

Second, they added the control input *u*(*t*) and the initial conditions of the state variables of the current time interval 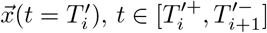 as input nodes to the neural networks. If the initial conditions 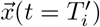 are not known, one can instead use the output of the neural network for the last time point of the previous control input 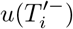. Figure 25 shows how the data propagation works in a PINC.

**Figure 25.**
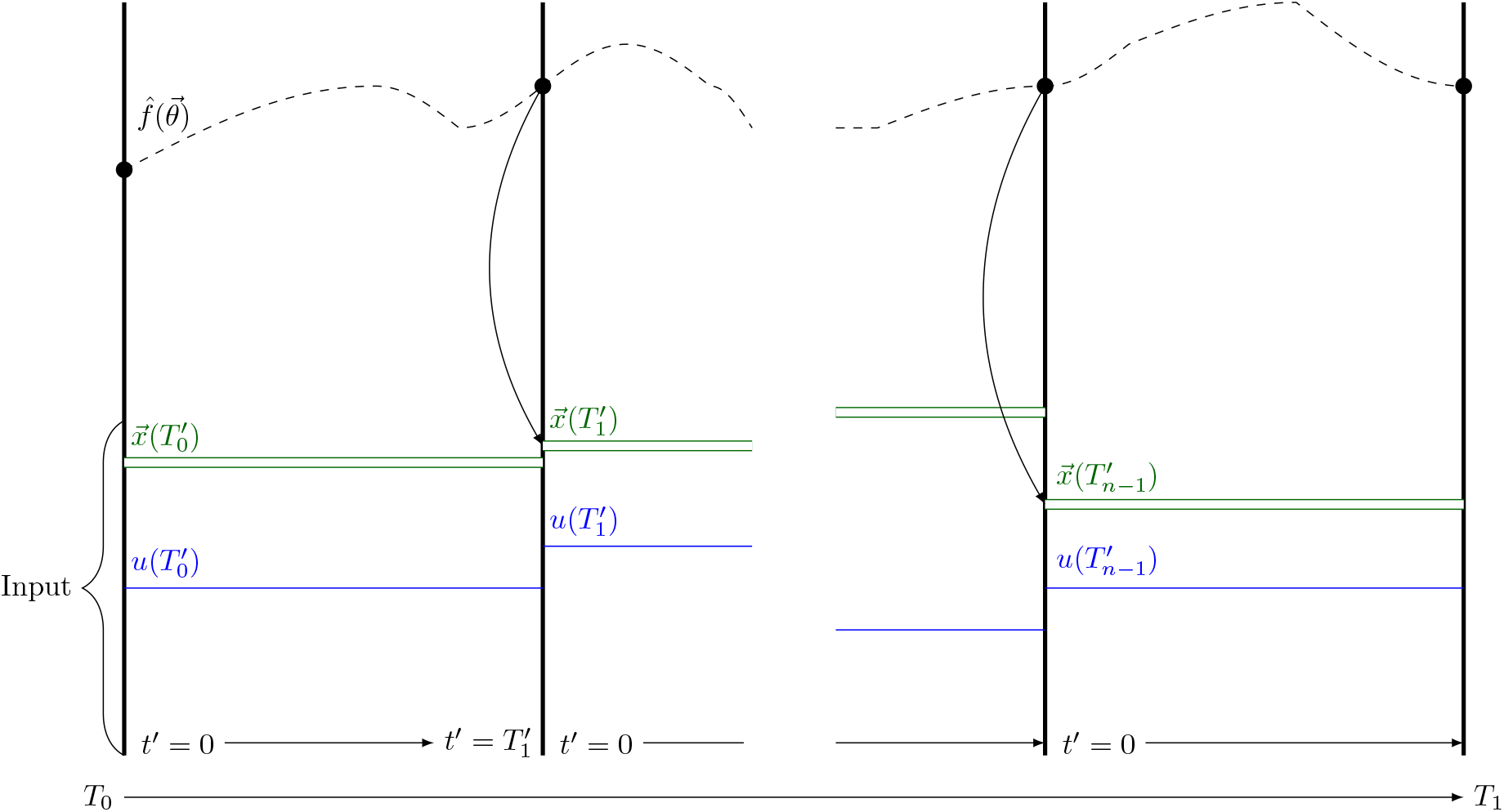
Schematic of data propagation in a PINC based on a figure in the original paper by Antonelo et al. [2021]. *u* is the control input of the different intervals. 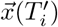 represents the corresponding initial conditions. 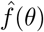 is the function learned by the neural network.

#### 5.3.2 Adaptation for Parameter Inference Deep Learning

Based on the original implementation of PINC by Antonelo et al. [2021], we adapted the parameter inference algorithm by Yazdani et al. [2020] to allow for control inputs. To that end, we added the possibility to automatically detect glutamate stimulation intervals, extended the neural network architecture, and adapted the learning process. An overview of the extended algorithm is given in Figure 26. The different changes are explained further in the following sections.

**Figure 26.**
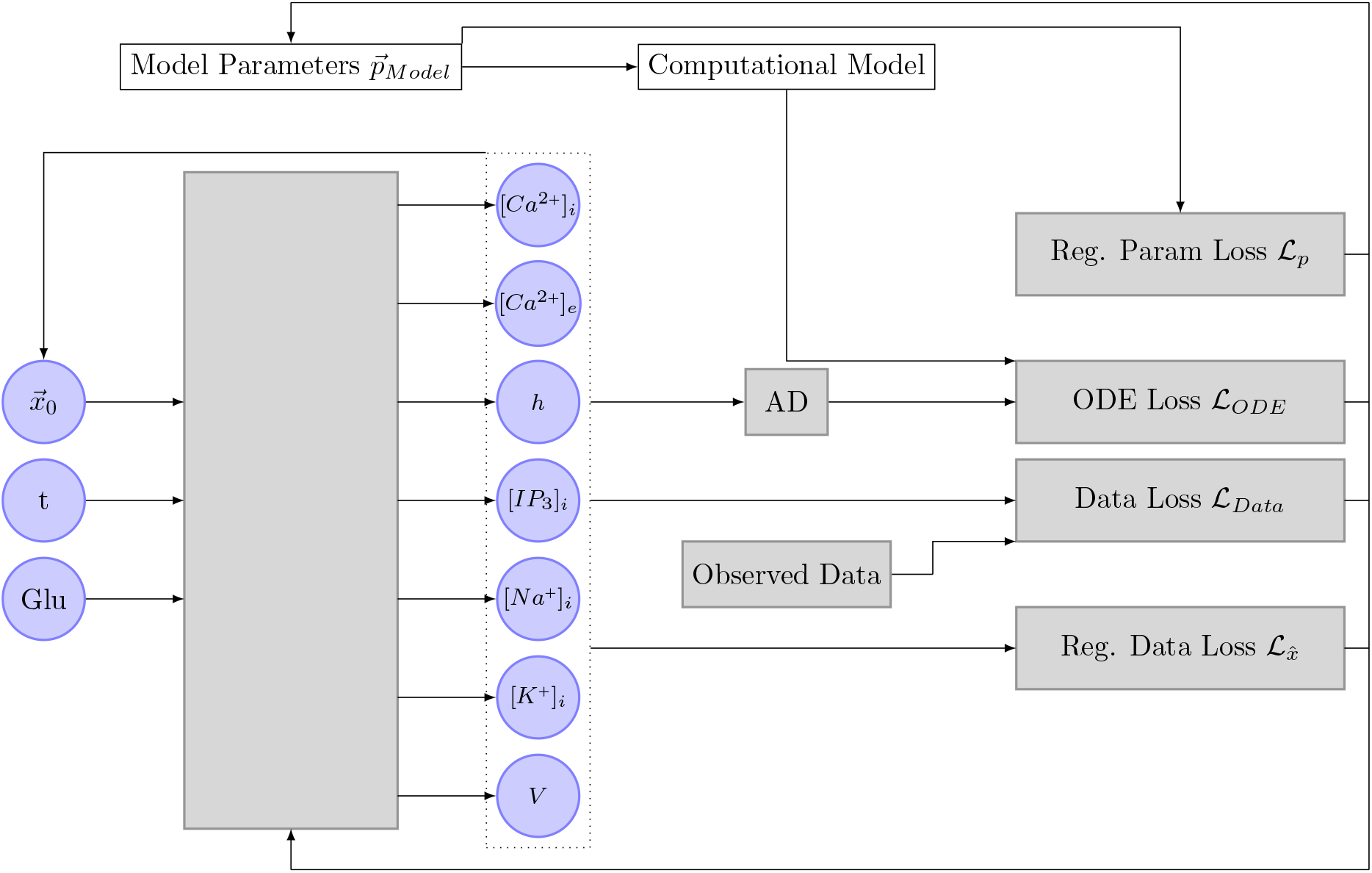
This schematic shows the adaptation of the algorithm by Antonelo et al. [2021] to the Oschmann et al. [model and in combination with the deep learning algorithm initially developed by Yazdani et al. [2020]. The input of the neural network is expanded with a control input (`Glu`) and initial conditions 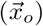. The initial conditions of each interval are predicted by the neural network itself and replace the previously used concept of ℒ_*Aux*_. AD stands for automatic differentiation.

##### Interval Detection

As a first step, changes in glutamate stimulation had to be detected. To that end, we assigned a `interval number` to each data point. The exact value of this `interval number` is technically unimportant, as long as each interval has a unique identifier. For simplicity, we chose ascending numbers. While the `interval number` does not get fed into the network, it is an important identifier to feed the correct initial conditions into the network. Furthermore, it simplifies the process of knowing the time frame of each stimulation interval. The `interval number` for each data point was determined by comparing the glutamate stimulation at consecutive time steps with each other. If no change larger than ϵ was detected, the data point was assigned the current interval number. Otherwise, we increased the current `interval number` before assigning it to the current data point and proceeding.

In this manuscript, we always assumed that the glutamate stimulation is noise free. Therefore, we chose ϵ = 0 for all inference experiments. However, noise could easily be incorporated by setting larger values of ϵ.

##### Input and Feature Transform

Next, we had to change the input- and feature transform layer of the neural network. To that end, we added eight input nodes to the input and feature transform layer of the neural network (one for the value of the glutamate stimulation and seven for the initial conditions).

Furthermore, we extended the scaling of the input time t by the shifting mechanism explained in Equation 75:

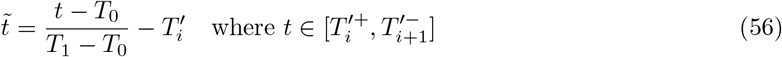

Initial conditions were scaled with the inverse of 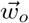 described earlier (Table 2). The glutamate stimulation was scaled linearly to be between one and two.

##### Training Process

Due to the addition of the initial values to the input layer, the training process had to be extended with a mechanism that gauges the initial conditions of each interval before the actual training step. We implemented two different versions. The simpler version, `Version A`, assumes that the initial states of every interval are known and expands the neural network input accordingly. `Version B` is more complex and only assumes that the initial values at *x*(*t* = 0) are known. At each epoch, the algorithm starts by predicting the initial values of every interval. Since the dynamics of the state variables are assumed to be continuous, this can be done as an iterative process. Starting at interval *i* = 1 and using the initial conditions of the interval i −1, the neural network is used to predict the end state of the interval *i* −1. At each training step, the loaded network inputs are then concatenated with the appropriate, predicted initial states. In theory, a combination of `Version A` and `Version B` would be possible. In that combination, `Version B` would be used for unobserved state variables and `Version A` for the observed state variables. The incorporation of 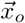 into the neural network replaces the concept of auxiliary loss.

###### Algorithm 2

Overview over the deep learning algorithm adapted for control inputs

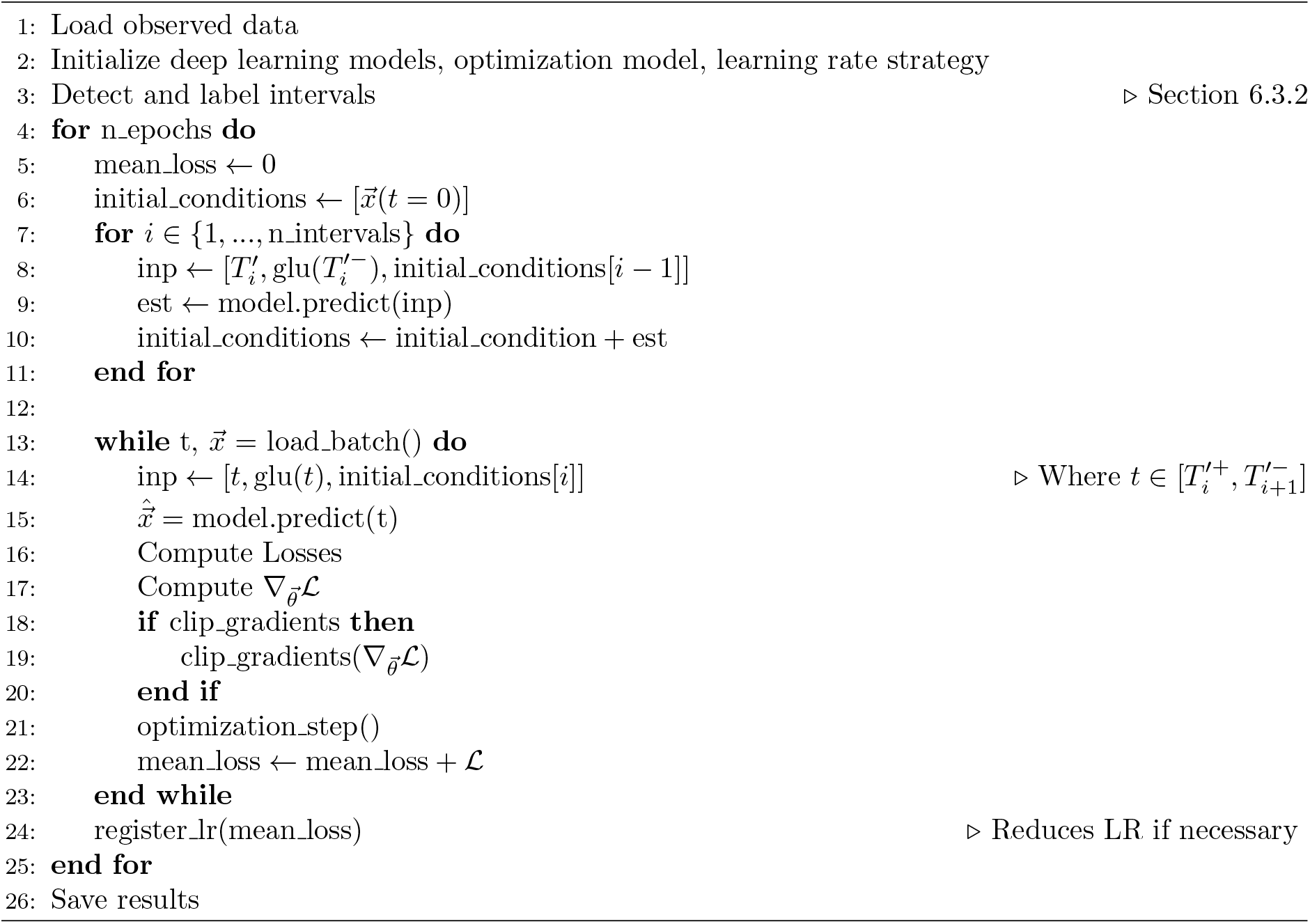

## 6 Methods, Part 2

In this section, we describe several methods that aim at stabilizing the inference of parameters in the Oschmann et al. [model. In Section 6.1, we explain a change to the Oschmann et al. [model that aims at stabilizing the *V*_*m*_ problem shown in Section 4.2. Based on a paper by Wang et al. [2020], we show methods to improve gradient pathologies during the inference process in Section 6.2. Last, Section 6.3 proposes the addition of control inputs to the neural network as was originally done by Antonelo et al. [2021].

### 6.1 Adapted Leak Currents and their subsequent Changes in Neural Network Parameters

As was observed in Section 4.2, the gradients of [*Na*^*+*^]_*i*_, [*K*^*+*^]_*i*_, and *V*_*m*_ returned by the computational Oschmann et al. [model are extremely sensitive to small errors in the input states. In part, this is due to the way the leak currents are computed. The original model computes the leak currents as

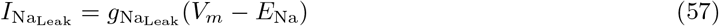

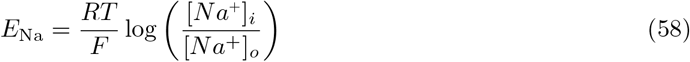

and

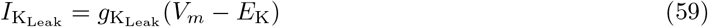

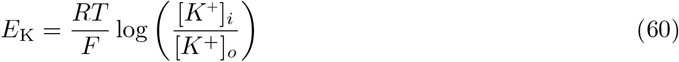

where *F* is the Faraday constant, *R* is the molar gas constant and *T* is the current temperature. This way of computing *E*_*K*_ and *E*_Na_ introduces a high level of sensitivity to the computations of the leak current, especially as the outer- and inner concentrations of both K^+^ and Na^+^ are dependent on the amount of [*Na*^+^]_*i*_ and [*K*^+^]_*i*_. To decouple this sensitivity, we replaced the dynamic computation of *E*_Na_ and *E*_K_ with constants, as is regularly done in other computational astrocyte models [Farr and David, 2011, Flanagan et al., 2018]. The used constants are equal to the known reversal potentials of Na^+^ and K^+^ and are listed in Table 10.

While the new leak computation does not change the general behavior of the simulation, it does change the order of magnitude of the computed gradients. The changed gradients require the usage of adapted weights for the parameter inference algorithm. These weights are listed in Table 11.

### 6.2 Gradient Pathologies in PINNs

While we trained my neural network with the configurations described in Section 3.3, it became obvious that the training process was not as stable and fast as expected. Wang et al. [2020] discovered and addressed one major mode of failure in PINNs. According to them, numerical stiffness might lead to unbalanced learning gradients during the back-propagation step in model training. They solved the problems in two ways. First, they suggested an algorithm that outbalances different loss terms. Second, they changed the model architecture to include a transformer network. In the following sections, we shortly describe their propositions and then explain how we adapted them for my model.

#### 6.2.1 Learning Rate Annealing

##### Original Implementation

To give different amounts of importance to different loss terms, loss terms are usually weighted. Assuming the loss functions consist of an ODE loss ℒ_*ODE*_ and *M* different data loss terms, such as different kinds of measurements or boundary conditions, the total loss can be written as

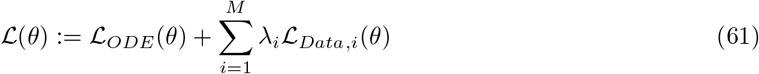

where *λ*_*i*_ is the weight of ℒ_*Data,i*_. Based on the optimization method `Adam` [Kingma and Ba, 2014] explained earlier, the authors suggested scaling the weights according to the ratio between the largest and average gradient of the different loss terms. Let

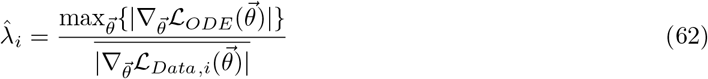

<u>where</u> 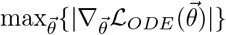 is the largest absolute parameter gradient of the ODE loss and 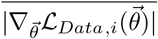 denotes the mean absolute parameter gradient of the different data loss terms. Due to the possibly high variance of 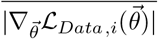, it was suggested to not directly use 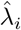 for weighting but rather to compute a running average using the equation

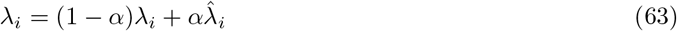

with *α* ∈ [0.5, 0.9]. Assuming `SGD` optimization (Equation 12) is used, the optimization step becomes

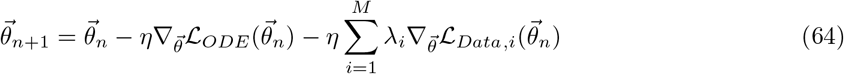

where *n* stands for the *n*-th iteration and ∇ is the learning rate. Wang et al. [2020] suggest to use a learning rate of *η* = 1e^−3^.

##### Adaption for Parameter Inference Deep Learning

In contrast to the examples by [Wang et al., 2020], the computational model at hand [Oschmann et al., 2017] consists of multiple ODEs. Furthermore, observations usually only exist for a subset of the given ODEs. This leads to the question of how the learning rate annealing algorithm should be adapted for my model. We tested three different strategies (`A`, `B`, `C`) further described below. To reduce the computational effort of computing 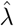, we only performed an update step every 50th epoch. The different strategies are visualized in Figure 23.

##### Strategy A

First, we matched the weight of the first ODE loss gradient 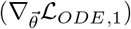 against all other loss gradients, both ODE loss and data loss, separately. This idea was motivated by the observation that it is not about balancing the ODE loss with the data loss, but about balancing all terms with each other. By setting 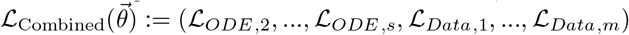 the total loss can be defined as

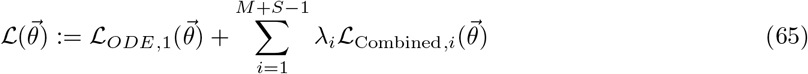

Then, 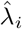 becomes

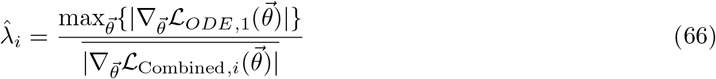

and is used in combination with the moving average Equation 63 to compute *λ*_*i*_.

##### Strategy B

For my second strategy, we assumed that the different ODEs are already balanced out well enough through the loss weighting described in Section 3.3.4. Therefore, the weighting only has to be adjusted between an ODE loss and its respective data loss. If an ODE does not have a counterpart, we do not change the weighting. Assuming no regularization and auxiliary losses, the total loss can be written in the form 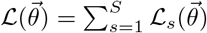 with

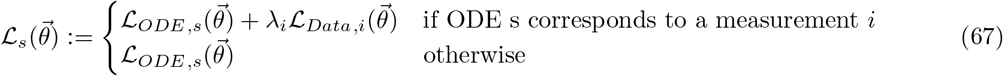

The weights are then computed using

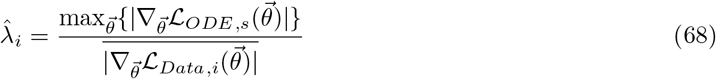

together with the moving average Equation 63.

##### Strategy C

For my last strategy, we made the same assumption as for Strategy `B`, but rather than balancing each ODE against its counterpart, we took the ratio between the largest gradient of the sum of all ODE losses and the mean gradient of the different data losses. In this form, the total loss is written as

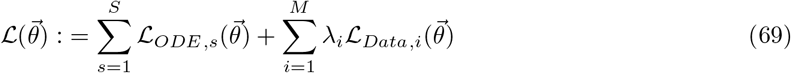

and the temporary weight becomes

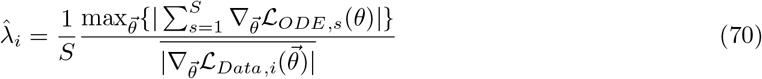

#### 6.2.2 Improved Fully Connected Architecture

The second improvement by Wang et al. [2020] concerned the architecture of the neural network itself and was based on the idea of a Transformer [Vaswani et al., 2017]. Transformers are often used in natural language processing or sequence transduction tasks and offer an alternative to the more commonly known recurrent or convolutional neural networks. Broadly speaking, a transformer considers possible multiplicative connections between different input nodes and strengthens the influence of input nodes on later network layers.

In the context of their paper, Wang et al. [2020] adapted the idea of transformer networks to PINNs by adding two additional network layers 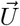 and 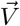. Just as the first fully connected neural network layer, 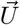 and 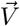 are directly connected to the input layer. They consist of the same number of nodes as all the other network layers. In form of equations, 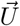and 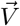 are defined through

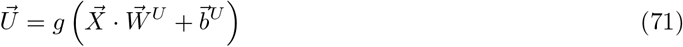

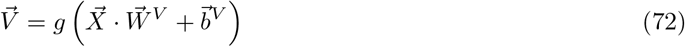

where *g* is the activation function, 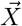 the input layer, 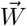 and 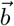 are the layers parameter. To enhance the network’s performance, 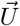 and 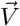 are multiplied component-wise to the output of the normal network layers described in Equation 10. The forward propagation equations, therefore, change to

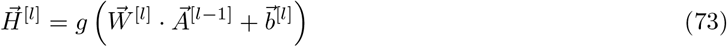

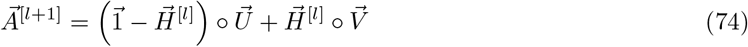

where ° denotes component-wise multiplication. Note that this change does not affect Equation 11 for the output layer of the neural network. Figure 24 shows the addition of 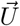 and 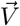 to a fully connected neural network. In the context of this manuscript, we followed the original implementation by Wang et al. [2020] exactly.

### 6.3 Control Input

In the context of ODEs, PINNs attempt to learn the relationship between a continuous time input *t* and several state variables 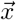. One of the major drawbacks of this method is that external events, such as a glutamate release by a neighboring neuron, can not be taken into account. Therefore, the glutamate level has either to be known or inferred at every point in time. While this might be possible under some preconditions, it is not feasible and further prohibits the use of multiple, different measurement sets to train one specific model.

The same problem is often faced in the context of control theory. While processes in for example the oil, gas, or robotics industry can often be modeled through differential equations, they usually have some dependence on external control inputs. To counteract this problem, Antonelo et al. [2021] recently proposed an adapted PINN algorithm that allows for control inputs. The concept is called *Physics-Informed Neural Nets-based Control* (PINC) and will be detailed further in the next Section 6.3.1. Section 6.3.2 then details how we adapted and implemented the concept of PINC to further improve on the parameter inference algorithm proposed by Yazdani et al. [2020] implemented in the context of this manuscript.

#### 6.3.1 Original Implementation

Inspired by multiple shooting and collocation methods, Antonelo et al. [2021] changed the original PINN algorithm [Raissi et al., 2017] in two significant ways. The first change is concerned with the input time *t*. Rather than attempting to learn how the state variables change over the whole time horizon, they suggested letting the network learn how the state variables have behaved since the last change in control input *u*(*t*). To that end, they subdivided the time interval [*T*_0_, *T*_1_] into multiple, smaller subintervals.

Assuming the control input is given by a piecewise constant function *u*(*t*), they split the time intervals at the points of discontinuity (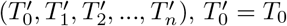 and 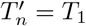, of *u*(*t*). Then, the input to the neural network is changed from *t* to *t*^*′*^ where *t*^*′*^ indicates how much time has passed since the beginning of the current subinterval.

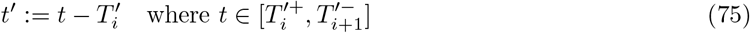

Second, they added the control input *u*(*t*) and the initial conditions of the state variables of the current time interval 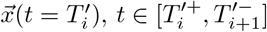 as input nodes to the neural networks. If the initial conditions 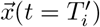 are not known, one can instead use the output of the neural network for the last time point of the previous control input 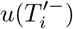. Figure 25 shows how the data propagation works in a PINC.

#### 6.3.2 Adaptation for Parameter Inference Deep Learning

Based on the original implementation of PINC by Antonelo et al. [2021], we adapted the parameter inference algorithm by Yazdani et al. [2020] to allow for control inputs. To that end, we added the possibility to automatically detect glutamate stimulation intervals, extended the neural network architecture and adapted the learning process. An overview of the extended algorithm is given in Figure 26. The different changes are explained further in the following sections.

##### Interval Detection

As a first step, changes in glutamate stimulation had to be detected. To that end, we assigned a `interval number` to each data point. The exact value of this `interval number` is technically unimportant, as long as each interval has a unique identifier. For simplicity, we chose ascending numbers. While the `interval number` does not get fed into the network, it is an important identifier to feed the correct initial conditions into the network. Furthermore, it simplifies the process of knowing the time frame of each stimulation interval. The `interval number` for each data point was determined by comparing the glutamate stimulation at consecutive time steps with each other. If no change larger than *ϵ* was detected, the data point was assigned the current `interval number`. Otherwise, we increased the current `interval number` before assigning it to the current data point and proceeding.

In this manuscript, we always assumed that glutamate stimulation is noise-free. Therefore, we chose *ϵ* = 0 for all inference experiments. However, noise could easily be incorporated by setting larger values of *ϵ*.

##### Input and Feature Transform

Next, we had to change the input- and feature transform layer of the neural network. To that end, we added eight input nodes to the input and feature transform layer of the neural network (one for the value of the glutamate stimulation, and seven for the initial conditions). Furthermore, we extended the scaling of the input time *t* by the shifting mechanism explained in Equation 75:

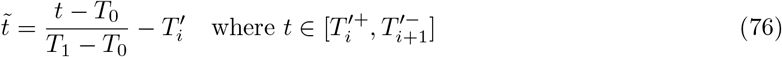

Initial conditions were scaled with the inverse of 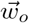 described earlier (Table 2). The glutamate stimulation was scaled linearly to be between one and two.

##### Training Process

Due to the addition of the initial values to the input layer, the training process had to be extended with a mechanism that gauges the initial conditions of each interval before the actual training step. We implemented two different versions. The simpler version, `Version A`, assumes that the initial states of every interval are known and expands the neural network input accordingly. `Version B` is more complex and only assumes that the initial values at *x*(*t* = 0) are known. At each epoch, the algorithm starts by predicting the initial values of every interval. Since the dynamics of the state variables are assumed to be continuous, this can be done as an iterative process. Starting at interval *i* = 1 and using the initial conditions of the interval *i* −1, the neural network is used to predict the end state of the interval *i* −1. At each training step, the loaded network inputs are then concatenated with the appropriate, predicted initial states. In theory, a combination of `Version A` and `Version B` would be possible. In that combination, `Version B` would be used for unobserved state variables and `Version A` for the observed state variables. The incorporation of 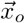 into the neural network replaces the concept of auxiliary loss.

###### Algorithm 3

Overview over the deep learning algorithm adapted for control inputs

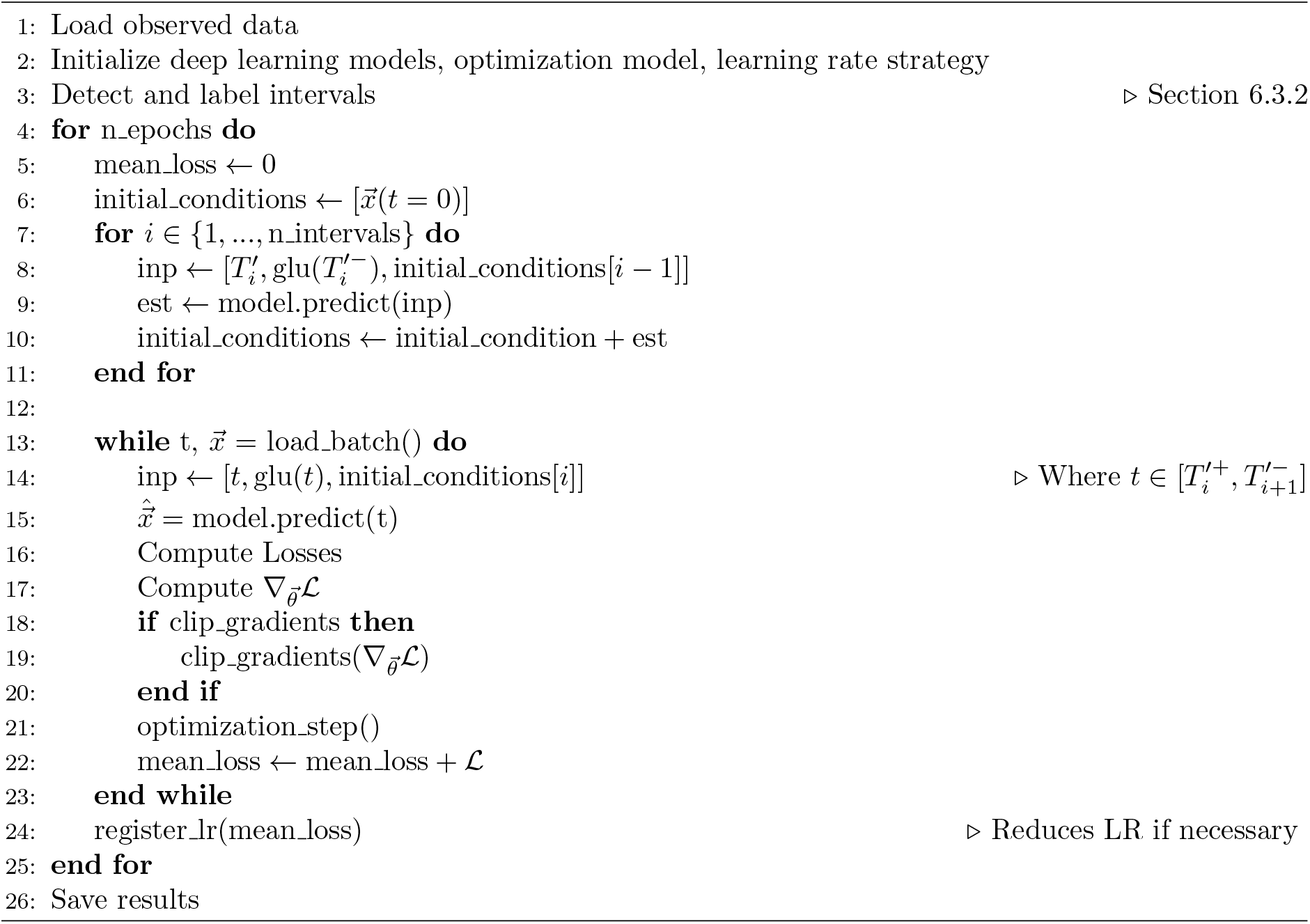

## 7 Results, Part 2

In this section, we show the inference results obtained from the methods described in section 6.

### 7.1 Effect of new Leak Computation

In this section, we visualize the dynamics of the state variables with the new leak computations, show the effect on the learning process of the learned gradients, and infer one parameter using the new leak computation.

#### 7.1.1 Dynamics

Figure 27 shows the changed dynamics after the computation of the leak currents was changed in the Oschmann et al. [model. It can be seen that the dynamics of [*Ca*^2+^]_*i*_, [*Ca*^2+^]_*e*_, *h*, and [*IP*_3_]_*i*_ were barely affected. This stands in contrast to the dynamics of [*Na*^+^]_*i*_, [*K*^+^]_*i*_, and *V*_*m*_. While these dynamics keep resembling step functions, the difference between the different step levels changed. Specifically, the differences in *V*_*m*_ and [*K*^+^]_*i*_ decreased, while the differences in [*Na*^+^]_*i*_ increased.

**Figure 27.**
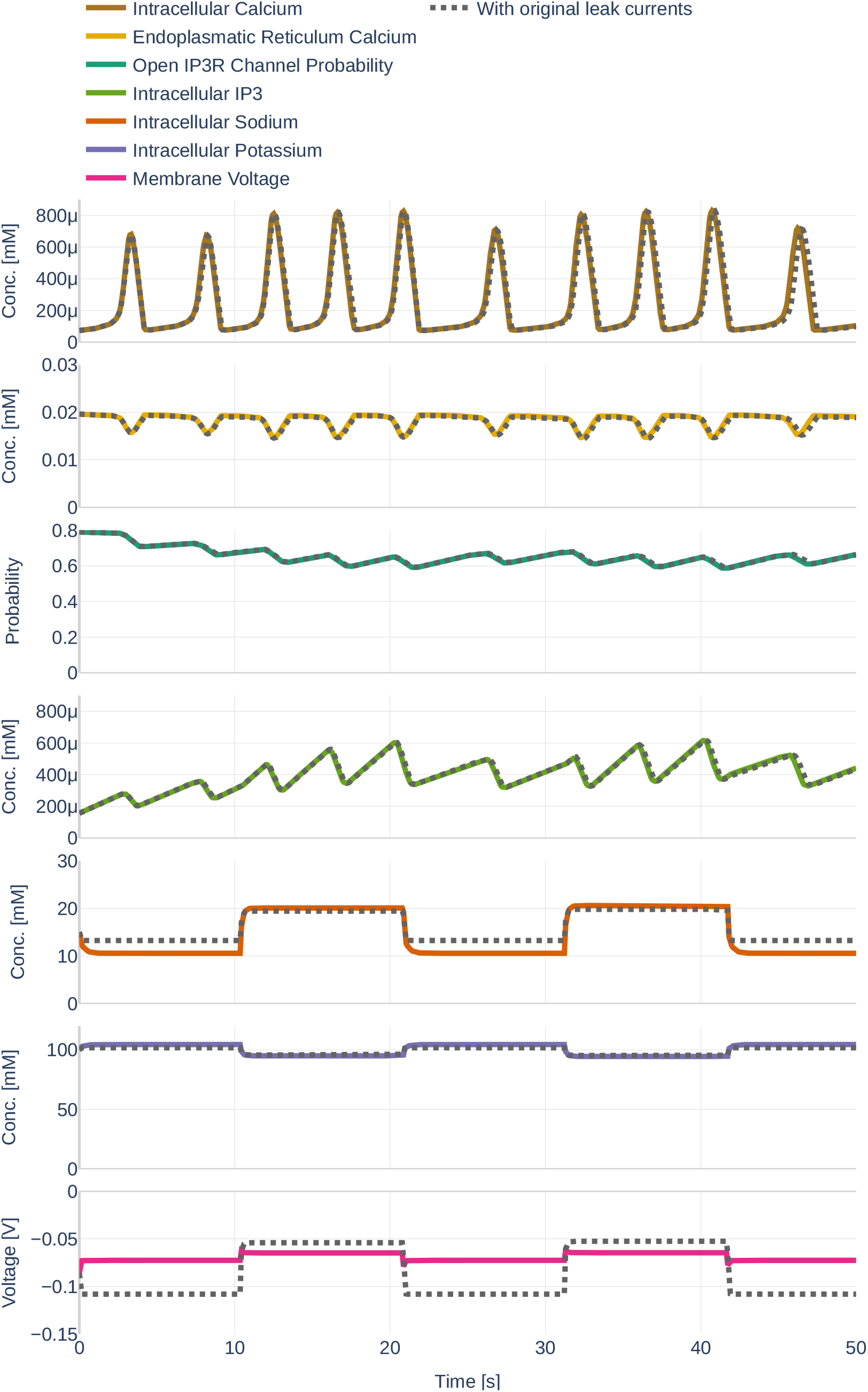
Dynamics of the Oschmann et al. [model when the leak current computation is changed (colored lines) as described in Section 6.1. The previous behavior is depicted as black, dotted lines. The used glutamate stimulation is shown in Figure 9.

#### 7.1.2 Learned Gradients

Next, we repeated the experiment performed in Section 4.2 and trained the neural network on the full data set `Parameter Study`, assuming that all dynamics were observed and that all parameters were known. Once all dynamics were learned accurately, we plotted the gradient of the neural network together with the gradient returned by the ODEs if the output of the neural network is fed into the computational model. The results can be seen in Figure 28. Noticeably, the network was able to learn all network gradients well. The only noteworthy errors occurred around the sharp gradients of [*Na*^+^]_*i*_, [*K*^+^]_*i*_ and *V*_*m*_ caused by changes in glutamate stimulation. Furthermore, the output of the computational model of [*K*^+^]_*i*_ and *V*_*m*_ is less error-prone than it was with the original leak computation.

**Figure 28.**
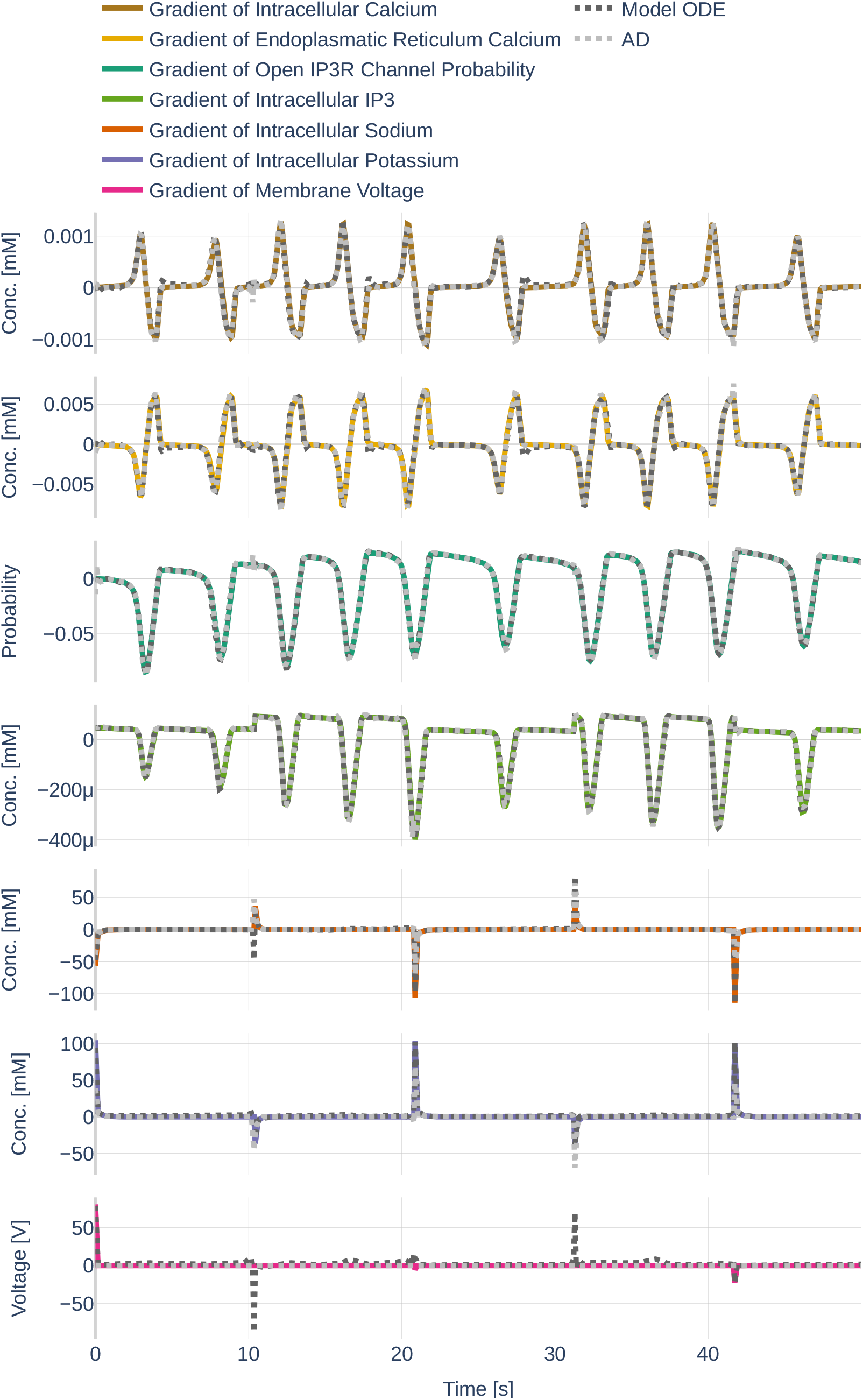
Gradients learned (gray) and gradients returned by the computational Oschmann et al. [model (black) if the learned dynamics are used as an input. The colored lines represent the gradients initially computed during the simulation. The used glutamate stimulation is shown in Figure 9.

#### 7.1.3 Inference

To test if the inference of parameters still works with the new leak computation, we repeated the inference of parameter *K*_NKA*mN*_ on a noiseless data set created with the changed computational model. The first inference experiment was done with the same weighting of the MSE terms as in the previous results section (Table 2). The second inference experiment was done with the new weights listed in Table 11. The results are shown in Figure 29. Generally, it can be seen that the inference of *K*_NKA*mN*_ is quite accurate for the new leak computation with old weights. However, the accuracy plot shows that the network fails to infer the not observed dynamics, indicating that, without learning rate reduction, the inference might have failed to converge and would instead have continued to increase. Interestingly, the network has a period of high accuracy 𝒜_all_ at the time when the parameter inference was approximately correct. Furthermore, the inference process took significantly longer than for the old leak computation. The inference of *K*_NKA*mN*_ with the new weighting of MSE terms did not work well. The inferred parameter only achieves 40% accuracy, the accuracy of the observed dynamics does not increase over 70%. However, the network is more successful in learning the dynamics of the observed state variables if the new leak computation is used. The exact inferred values and accuracies are listed in Table 12.

**Figure 29.**
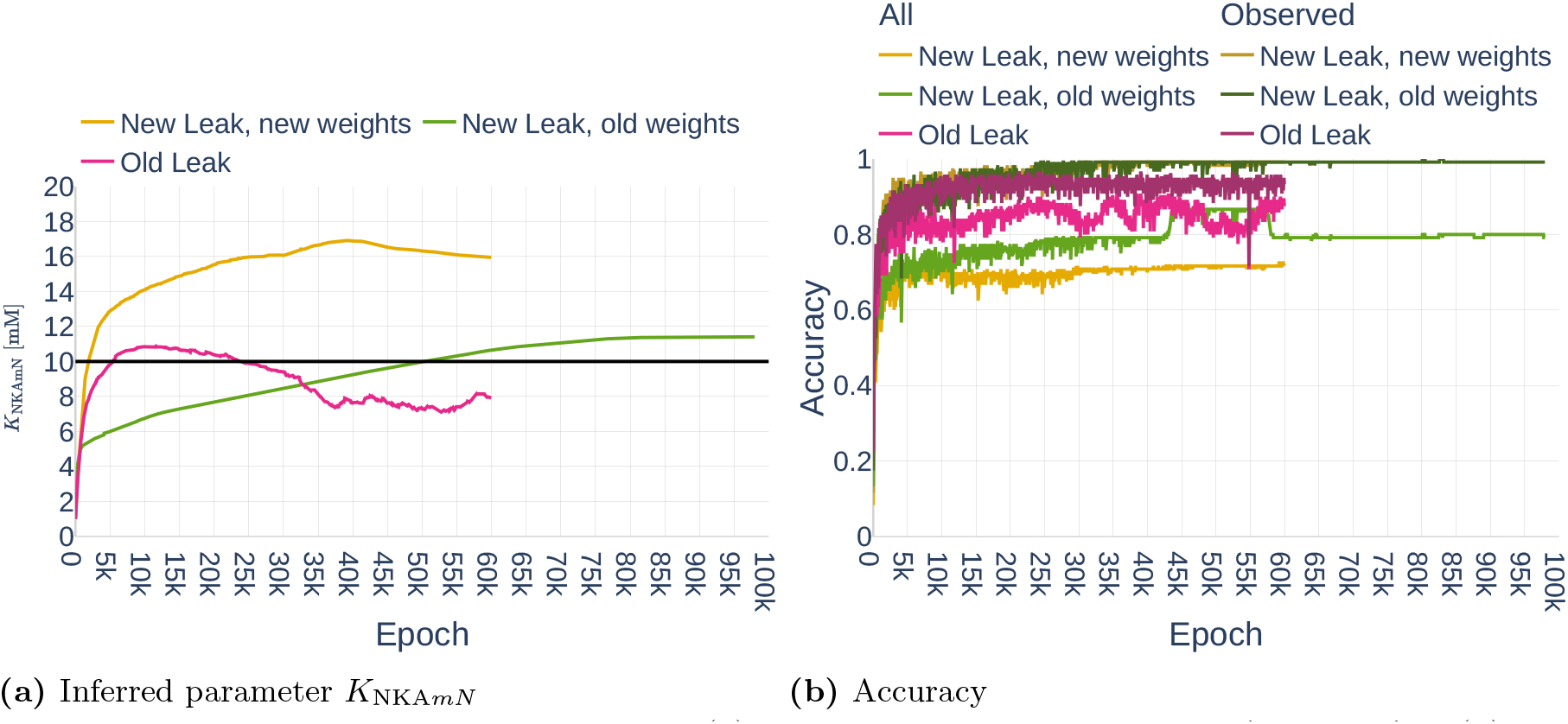
Inference of parameter *K*_NKA*mN*_ (a) and the respective accuracies 𝒜_*all*_ and 𝒜_*obs*_ (b) using the original leak computation in comparison to the inference of the same parameter using the new leak computation with two sets of weights. *old* weights refer to the weights used beforehand and listed in Table 2. *new* weights refer to the weights computed for the new leak computation and are listed in Table 11. The black line in (a) indicates the original value.

**Table 12.** Inferred parameter values for the different parameter inference experiments in this section. The original values and their scaling can be seen in Table 3. The abbreviation *Trans*. stands for Transformer, the abbreviation *LR* for learning rate.

| Simulation | Parameter | Unit | Inferred | $\mathcal{A}_{\bar{p}}$ | $\mathcal{A}_{\text{all}}$ | $\mathcal{A}_{\text{obs}}$ |
| --- | --- | --- | --- | --- | --- | --- |
| Inference with new leak |  |  |  |  |  |  |
| New leak, old weights | $K_{\text{NKAmN}}$ | $mM$ | 11.4 | 89.6% | 79.1% | 99.1% |
| New leak, new weights | $K_{\text{NKAmN}}$ | $mM$ | 15.95 | 40.5% | 71.6% | 99.1% |
| Old leak | $K_{\text{NKAmN}}$ | $mM$ | 7.9 | 79% | 75.5% | 93% |
| LR Annealing |  |  |  |  |  |  |
| Strategy A | $I_{\text{NCXmax}}$ | $\frac{pA}{\mu m^2}$ | 0.006 | -% | 60 % | 82.5% |
| Strategy B | $I_{\text{NCXmax}}$ | $\frac{pA}{\mu m^2}$ | 0.001084 | 91.6% | 90% | 97.5% |
| Strategy C | $I_{\text{NCXmax}}$ | $\frac{pA}{\mu m^2}$ | 0.001085 | 91.5% | 91.6% | 99.2% |
| Improved Architecture |  |  |  |  |  |  |
| Strategy B, Trans. | $I_{\text{NCXmax}}$ | $\frac{pA}{\mu m^2}$ | 0.00103 | 97% | 84.2% | 92.5% |
| Strategy C, Trans. | $I_{\text{NCXmax}}$ | $\frac{pA}{\mu m^2}$ | 0.000975 | 97.5% | 82.5% | 92.5% |
| PINC, Strategy B |  |  |  |  |  |  |
| Version A | $I_{\text{NCXmax}}$ | $\frac{pA}{\mu m^2}$ | 0.000981 | 98.1% | 87.5% | 96.6% |
| Version A, Trans. | $I_{\text{NCXmax}}$ | $\frac{pA}{\mu m^2}$ | 0.00111 | 89% | 83.3% | 98.3% |
| Version B | $I_{\text{NCXmax}}$ | $\frac{pA}{\mu m^2}$ | 0.00104 | 96% | 89.2% | 96.6% |
| Version B, Trans. | $I_{\text{NCXmax}}$ | $\frac{pA}{\mu m^2}$ | 0.009 | -% | 37.5 % | 51.7% |
| PINC, Strategy C |  |  |  |  |  |  |
| Version A | $I_{\text{NCXmax}}$ | $\frac{pA}{\mu m^2}$ | 0.00104 | 96% | 92.5% | 100% |
| Version A, Trans. | $I_{\text{NCXmax}}$ | $\frac{pA}{\mu m^2}$ | 0.00105 | 95% | 89.2% | 97.5% |
| Version B | $I_{\text{NCXmax}}$ | $\frac{pA}{\mu m^2}$ | 0.000978 | 97.8% | 88.3% | 95.8% |
| Version B, Trans. | $I_{\text{NCXmax}}$ | $\frac{pA}{\mu m^2}$ | 0.00098 | 98% | 88.3% | 97.5% |
| Noisy data, NCX |  |  |  |  |  |  |
| No noise | $I_{\text{NCXmax}}$ | $\frac{pA}{\mu m^2}$ | 0.00104 | 96% | 89.16% | 96.6% |
| Noise, repetition 1 | $I_{\text{NCXmax}}$ | $\frac{pA}{\mu m^2}$ | 0.0011 | 89% | 71.6% | 78.3% |
| Noise, repetition 2 | $I_{\text{NCXmax}}$ | $\frac{pA}{\mu m^2}$ | 0.0011 | 89% | 76.6% | 79.16% |
| Noise, repetition 3 | $I_{\text{NCXmax}}$ | $\frac{pA}{\mu m^2}$ | 0.00240 | -% | 75.0% | 80.8% |
| Noisy data, SERCA |  |  |  |  |  |  |
| No noise | $\nu_{ER}$ | $\frac{mM}{s}$ | 0.00449 | 99.7% | 85% | 92.5 % |
| Noise, repetition 1 | $\nu_{ER}$ | $\frac{mM}{s}$ | 0.0045 | 100% | 72.5% | 75.8% |
| Noise, repetition 2 | $\nu_{ER}$ | $\frac{mM}{s}$ | 0.00449 | 99.7% | 74.2% | 76.6% |
| Noise, repetition 3 | $\nu_{ER}$ | $\frac{mM}{s}$ | 0.0045 | 100% | 75.8% | 79.2% |

### 7.2 Gradient Pathologies

In this section, we show the effect of the strategies originally suggested by Wang et al. [2020]. The first subsection shows the inference of *I*_NCXmax_ using three different learning rate annealing strategies `Strategy A`, `Strategy B` and `Strategy C`. The second subsection shows the results for the combination of Transformers with `Strategy B` and `Strategy C`. All results are computed on the noise-free data set `Parameter Study` and all but the dynamics of [*Na*^+^]_*i*_ and [*K*^+^]_*i*_ were assumed to be observed. The leak computation was reset to the original version.

#### 7.2.1 Learning Rate Annealing with Different Strategies

Figure 30 shows the results for the inference of parameter *I*_NCXmax_ with activated learning rate annealing. The experiments were performed without decreases in learning rate and with a gradient clipping value of *c* = 100. It can easily be seen that `Strategy A` does not work. This strategy leads to large oscillations and inaccurate inference results. Furthermore, the achieved overall accuracy 𝒜_all_ is extremely low at 60%. Therefore, we did not consider `Strategy A` further after this point. However, `Strategy B` and `Strategy C` yielded good results. The exact inferred values are shown in Table 12. The learning process of both, `Strategy B` and `C`, showed a steady decrease in total loss and a steady increase in accuracy.

**Figure 30.**
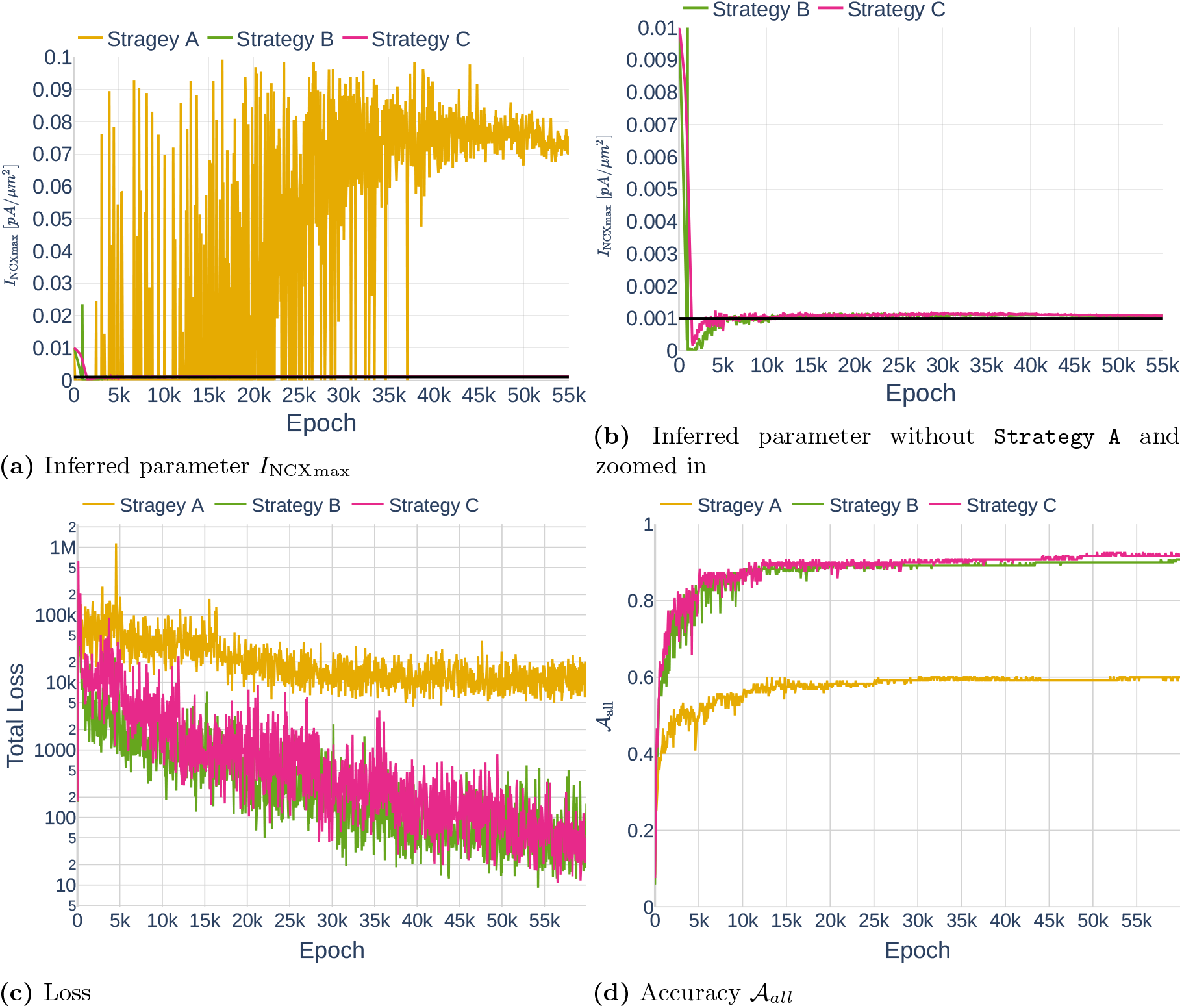
Comparison of different *λ* strategies when inferring parameter *I*_NCXmax_ (a; b shows a cutout for `Strategy B` and `C`) together with the respective loss values (c) and accuracies 𝒜_*all*_ (d). The black lines in (a) and (b) indicate the original value.

#### 7.2.2 Improved Fully Connected Architecture

Next, we added two Transformer layers to the fully connected network architecture as suggested by Yazdani et al. [2020]. The results can be seen in Figure 31. Although the transformer networks also succeed at inferring the parameter *I*_NCXmax_, it can be seen that the inference oscillates more heavily than for the networks without a transformer. Due to this high oscillation level, we considered the inferred parameter to be the average inferred parameter over the last 1000 epochs. The inferred values are listed in Table 12. It can be seen that the achieved parameter accuracies are higher than for the parameter inference experiments without Transformers. However, the accuracy of all dynamics 𝒜_All_ is lower for the Transformer architecture than for the normal one.

**Figure 31.**
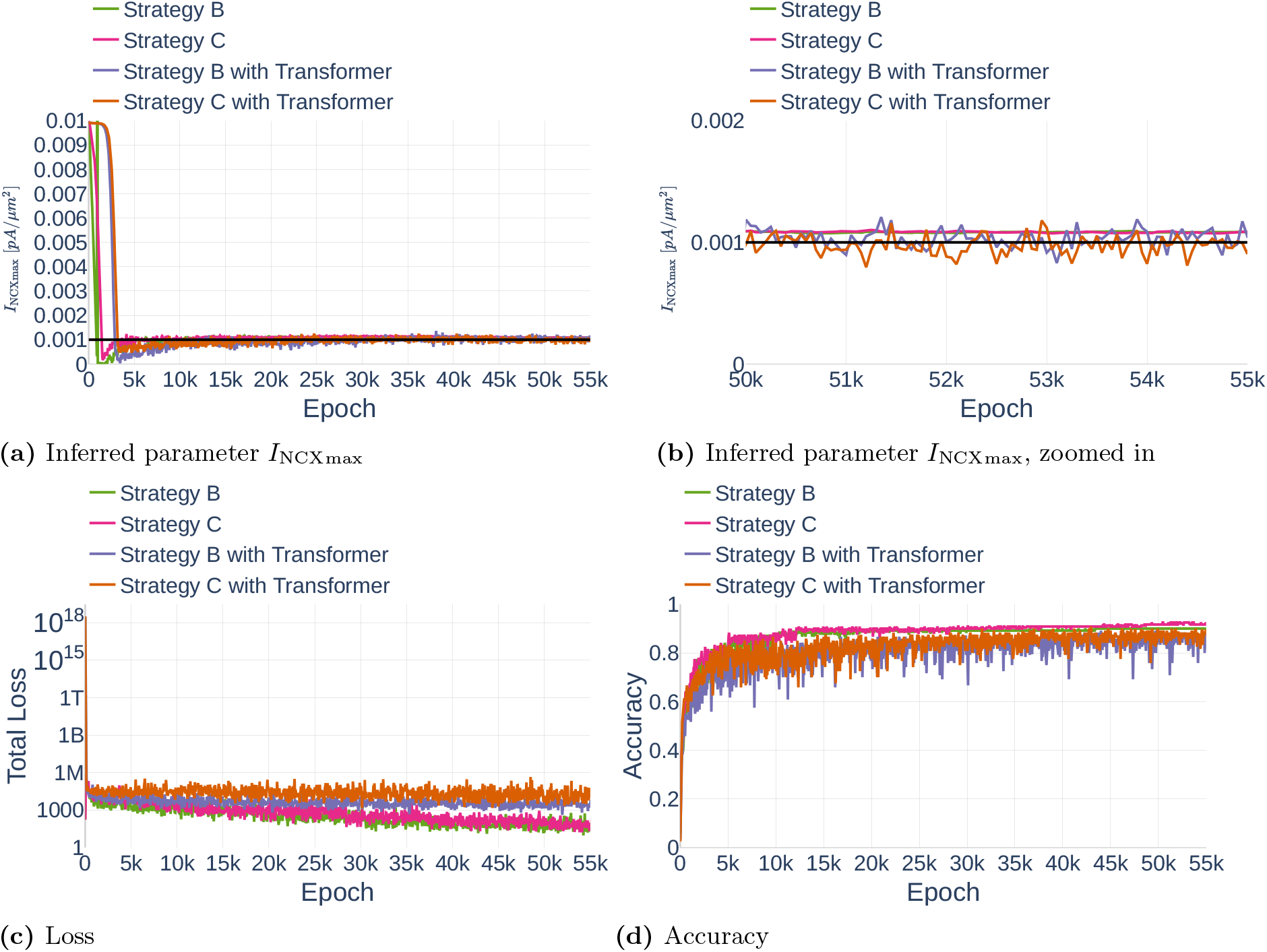
Comparison of different *λ* strategies when inferring parameter *I*_NCXmax_ with and without the addition of Transformers to the neural network (a; b shows a cutout). The respective loss values (c) and accuracies 𝒜_*all*_ (d) are shown. The black lines in (a) and (b) indicate the original value.

### 7.3 Control Inputs

In this section, we report the results for parameter inference using PINC and highlight one possible problem regarding the combination of learning with noise and PINC.

#### 7.3.1 Parameter Inference

Using the concept of PINC originally suggested by Antonelo et al. [2021], we tested the inference of *I*_NCXmax_ in combination with the different strategies to avoid gradient pathologies. The results of PINC in combination with `Strategy B` are shown in Figure 32. Similarly, Figure 33 shows the results for `Strategy C`. In the plots, `Version A` stands for the inference using PINC under the assumption that the initial states of each interval are known. For `Version B`, only the initial states of the first interval are assumed to be known, the initial states of the following intervals are predicted and continuously updated. The inference of *I*_NCXmax_ was successful for all but `Strategy B` in combination with a Transformer network. As can be seen in the plot of the loss value, the computed *λ* weights in this version likely became too large. As a consequence, the network was unable to learn. Another interesting observation is that when using `Strategy C` with `Version B` and no Transformer, the network inferred heavily oscillating values between epoch 1000 and epoch 2000, before stabilizing again. All other inference methods demonstrated highly stable convergence behavior. The exact values inferred are listed in Table 12. Surprisingly, the inference or parameters did work similarly well when the network was responsible for predicting its own initial states. Furthermore, parameter convergence was achieved faster than for traditional PINNs.

**Figure 32.**
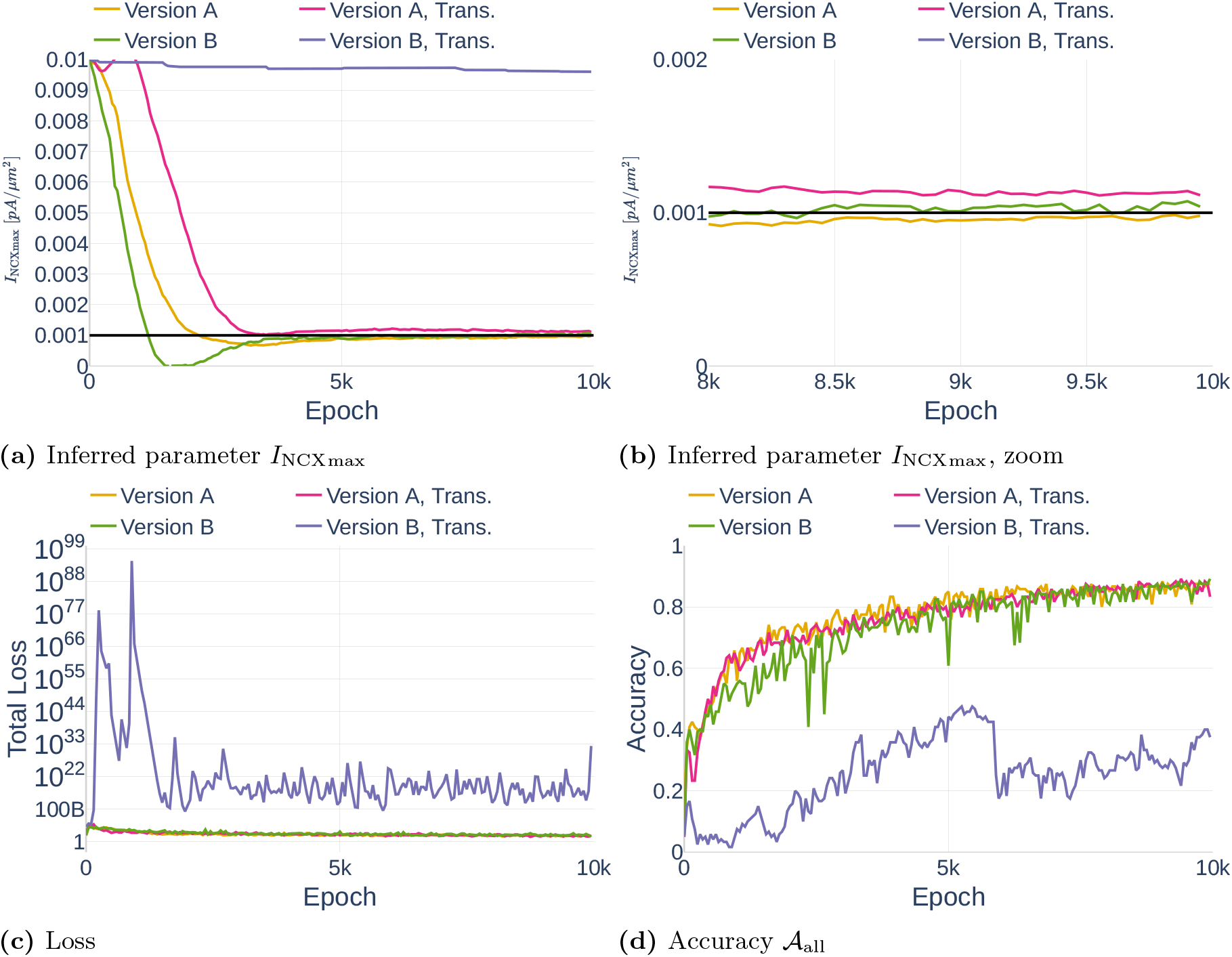
Inference of *I*_NCXmax_ using PINC in combination with `Strategy B` to avoid gradient pathologies (a). The respective loss values (c) and accuracies 𝒜_all_ (d) are also shown. The abbreviation *Trans*. stands for the addition of Transformers. The black lines in (a) and (b) indicate the original parameter value.

**Figure 33.**
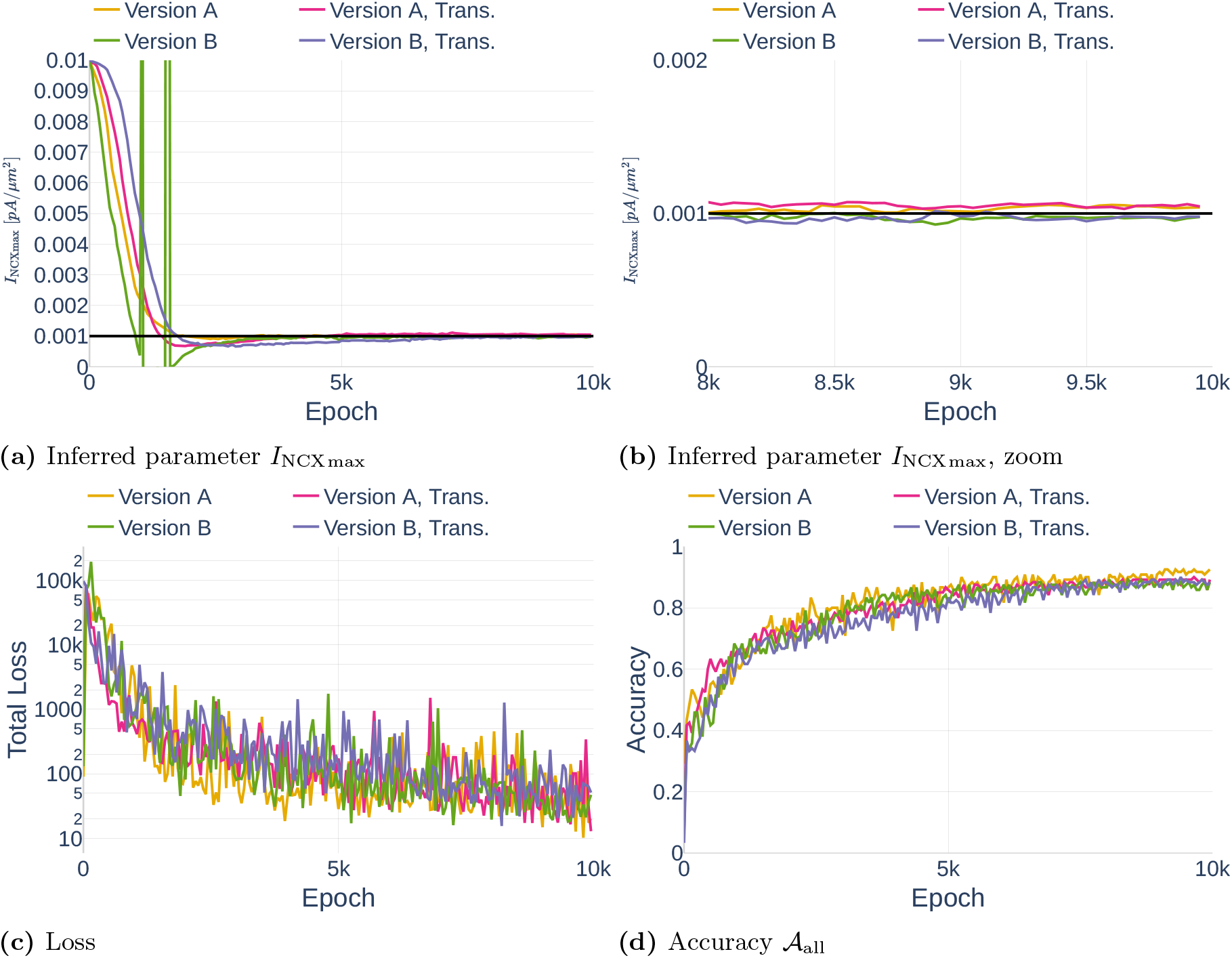
Inference of *I*_NCXmax_ using PINC in combination with `Strategy C` to avoid gradient pathologies. The abbreviation *Trans*. stands for the addition of Transformer networks. The black lines in (a) and (b) indicate the original parameter value.

#### 7.3.2 Parameter Inference with Noise

Next, we tested the inference of parameters using the data set `Noise`. Figure 34 shows the inference results of *I*_NCXmax_ without noise on the observed data and three repetitions of inference with noise on the observed data. It can be seen that for two out of three repetitions, the inference of *I*_NCXmax_ was successful. However, in `Repetition 3`, the inferred parameter converged towards a wrong value, indicating instability. When repeating the same stability experiment with parameter *ν*_*ER*_ of the *I*_Serca_ current, we did not observe the same problem (Figure 35). Not surprisingly, the achieved accuracy 𝒜 _all_ is lower for the repetitions where the observed data was noisy. Note that the term *accuracy* has a somewhat different meaning in the context of noisy data, as fitting all data perfectly would be an indication of overfitting.

**Figure 34.**
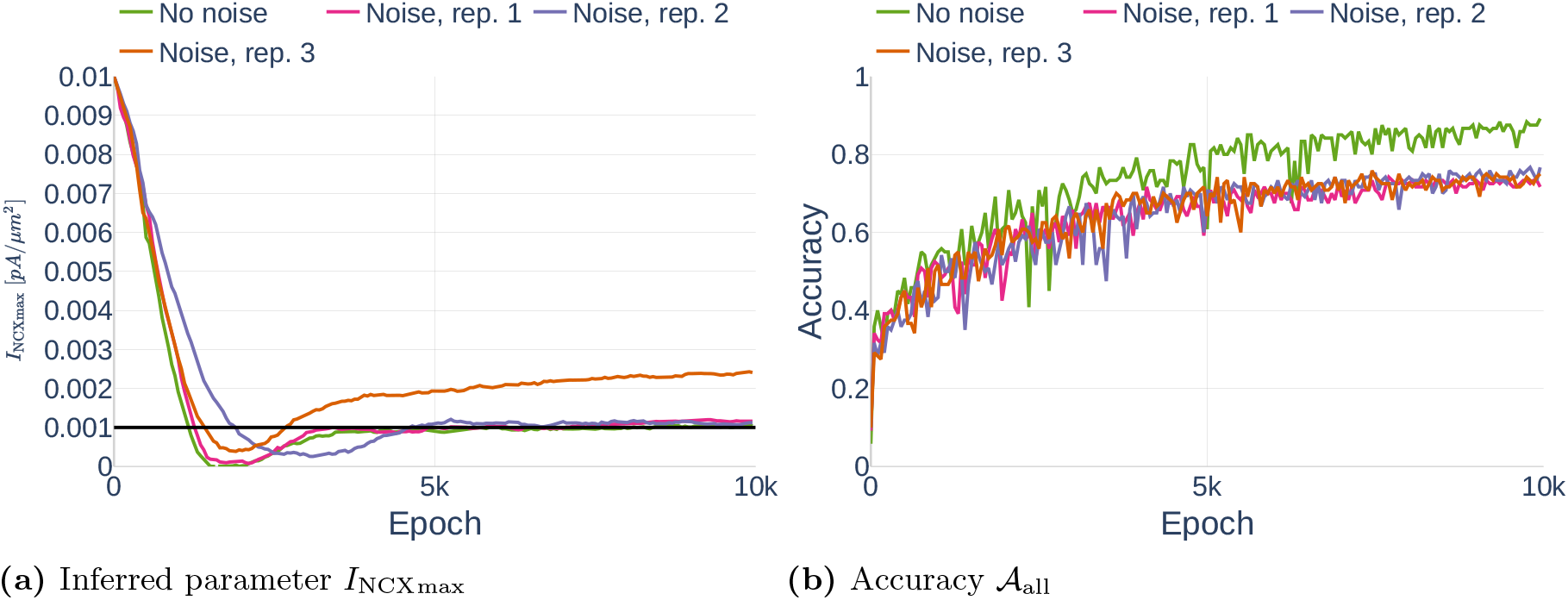
Inference of INCXmax (a) and respective accuracies 𝒜_all_ (b) when using PINC in combination with Strategy B (gradient pathologies), Version A (initial values) and noisy observation data. The black line in (a) indicates the original value.

**Figure 35.**
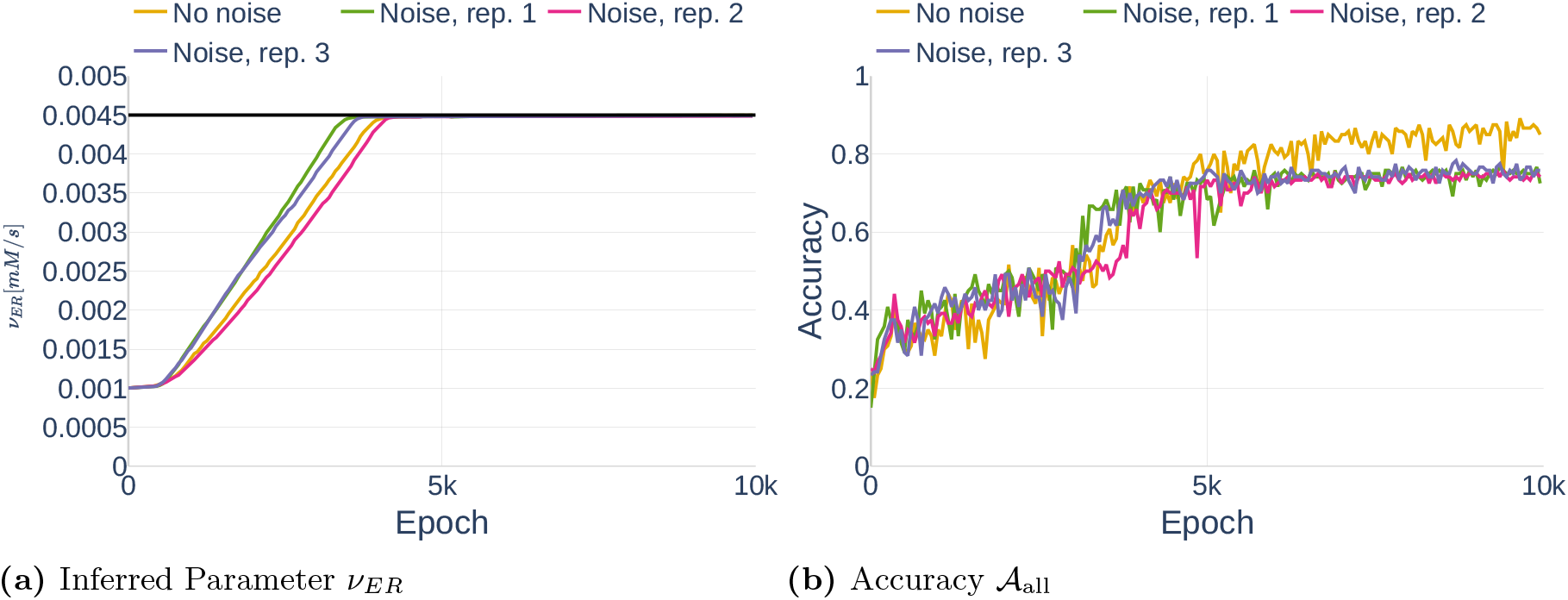
Inference of *ν*_*ER*_ (a) and the respective accuracy 𝒜_all_ (b) using PINC in combination with `Strategy B` (gradient pathologies), `Version A` (initial values) and noisy observation data. The black line in (a) indicates the original value.

### 7.4 Inference of Multiple Parameters

Using the created PINC algorithm and the PINN algorithm with learning rate annealing `Strategy C`, we inferred the parameters *ν*_*ER*_ and *K*_*ER*_ of *I*_Serca_. The dynamics of [*Ca*^2+^]_*i*_, [*Ca*^2+^]_*e*_, *h* and *V*_*m*_ were assumed to be observed and had 10% Gaussian noise on them (data set `Noise`). The results can be seen in Figure 36. Interestingly, PINN and PINC achieve around the same accuracy *A*_*all*_ and infer the same parameter *K*_*ER*_. At the same time, PINN only achieves an accuracy of 89.2% on *ν*_*ER*_ while the accuracy of PINC is as high as 97.3%. The exact inferred results can be seen in Table 13.

**Figure 36.**
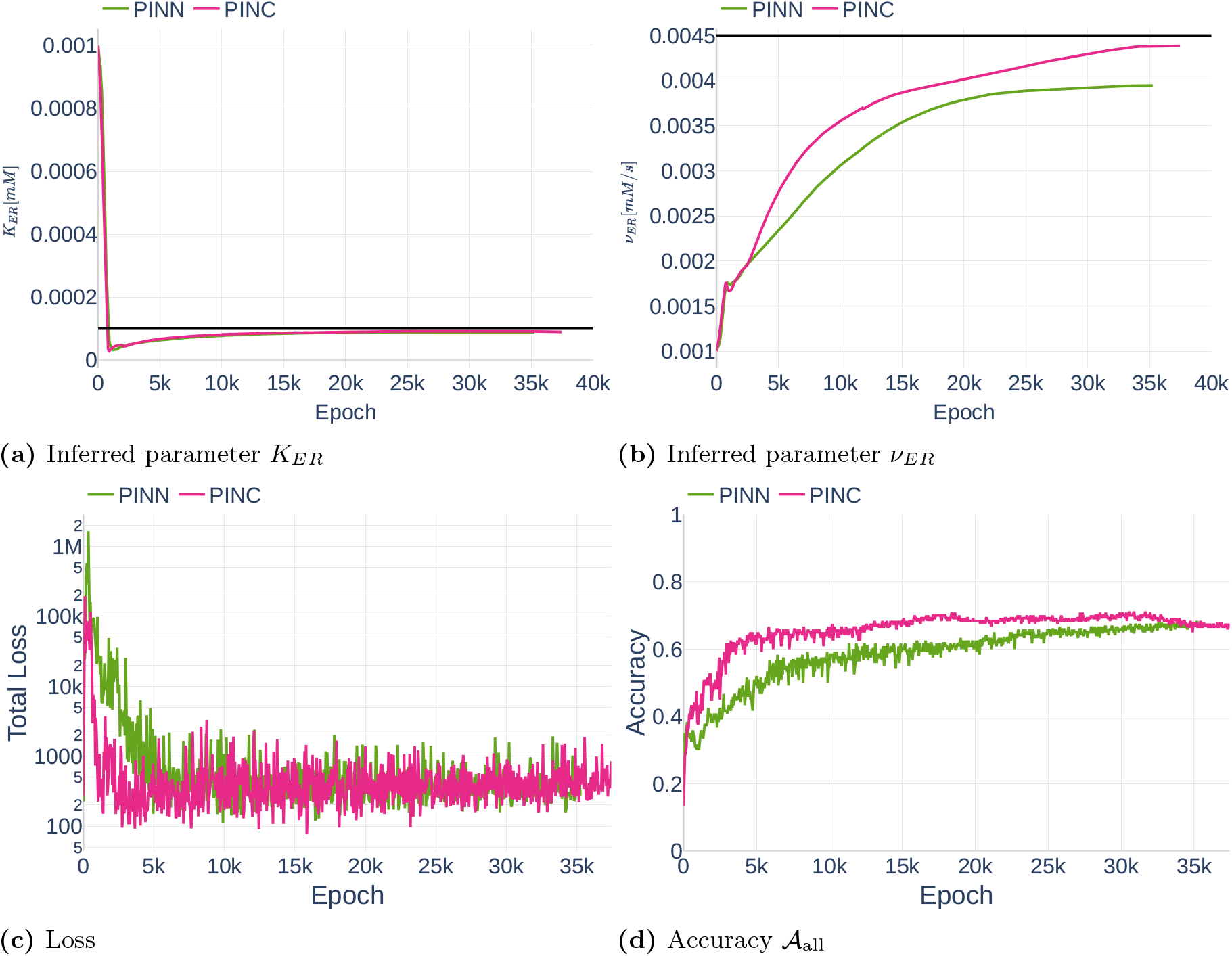
Inference of *K*_*ER*_ (a) and *ν*_*ER*_ (b) using normal physics informed neural networks in combination with `Strategy C` (PINN) and physics informed neural networks in combination with control inputs, `Version A` and `Strategy C` (PINC). The respective total loss values (c) and accuracies 𝒜_all_ (d) measured on noisy data are shown. The black lines in (a) and (b) indicate the original parameter values.

**Table 13.**
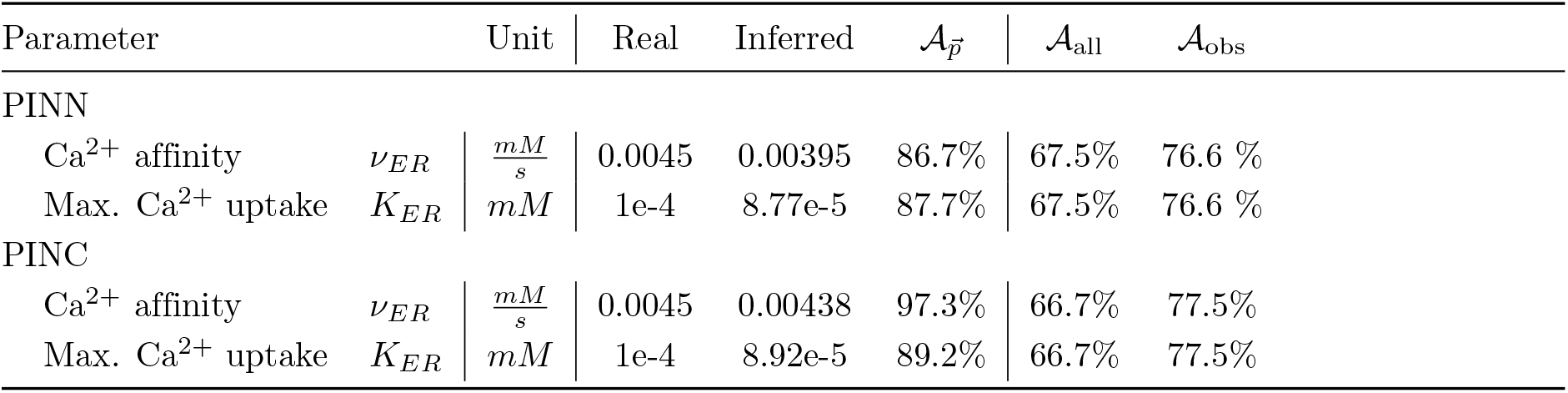
Achieved inference results when inferring the parameters of *I*_Serca_ using PINN and PINC. Note that the accuracy of the dynamics is measured on the data set `Noise`.

### 7.5 Runtimes

In this section, we examine the difference between executing the parameter inference algorithm on a GPU versus on a CPU. Furthermore, we detail the runtime of one and fifty epochs of different versions of the parameter inference algorithm.

#### 7.5.1 GPU vs. CPU

To ensure that the parameter inference can be done as fast as possible, we measured whether executing the algorithm on a normal CPU or on a GPU (Cuda) is faster. The runtime for different steps of the naive parameter inference algorithm are shown in Figure 37. Considered are data points from 50 epochs. Unexpectedly, it can be seen that the execution time is generally shorter on the CPU. This is especially true for the computation of ℒ _*ODE*_ and the optimization step.

**Figure 37.**
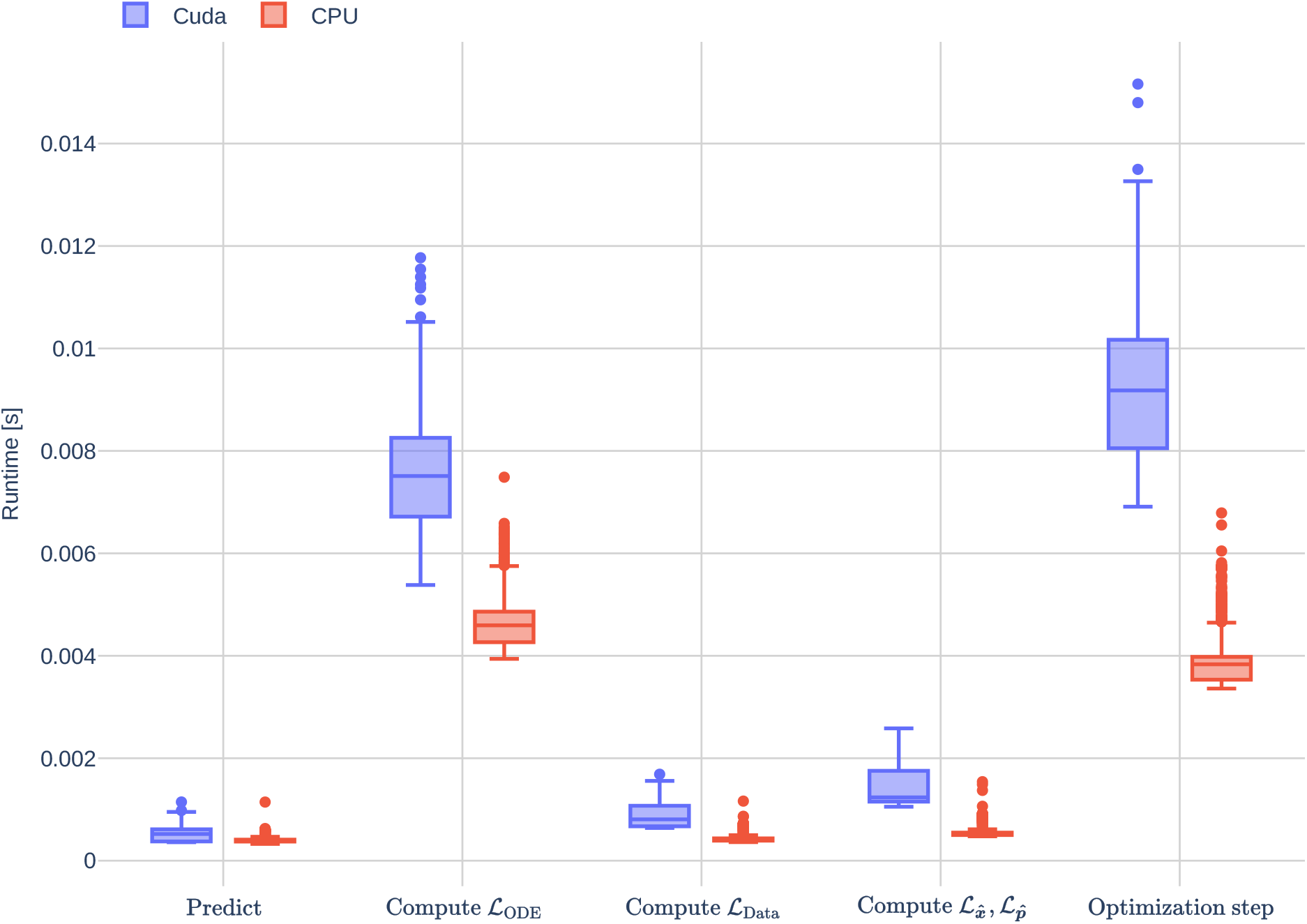
Runtimes of different parts of the deep learning algorithm on a GPU and on a CPU. Plotted are the run times of the first 50 epochs.

#### 7.5.2 Differences between Methods

The run times of one and 50 epochs of parameter inference of different versions of the algorithm are listed in Table 14. Not surprisingly, the initial version is the fastest. Computing the new weights *λ* in `Strategy B` and `Strategy C` every 10th iteration has a computational impact, increasing the runtime when looking at the runtime over 50 epochs. Because `Strategy B` requires more computations of 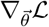, it is more expensive than `Strategy C`. As expected, the addition of Transformers increases the runtime even further. This can be explained by the fact that Transformers add additional connections, and therewith network parameters, to the neural network architecture, making the computation of 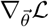 more expensive. Even more expensive is the neural network architecture with control inputs (PINC). This is likely due to the necessary prediction and concatenation of the loaded data with the initial conditions of the respective interval.

**Table 14.** This table lists the runtime of 1 or 50 epochs with different versions of the parameter inference algorithm. The runtimes were measured with a batch size of 32 and a total of 480 data points. Excluded is the time needed to initialize the models or to compute accuracies.

| Name | 1 Epoch [s] | 50 Epochs[s] | Description |
| --- | --- | --- | --- |
| Initial Version | $0.166 \pm 0.0023$ | $8.18 \pm 0.026$ | Section 3.3 |
| **Strategy B** | $0.164 \pm 0.0015$ | $8.59 \pm 0.242$ | Section 6.2.1 |
| **Strategy C** | $0.165 \pm 0.0010$ | $8.44 \pm 0.040$ | Section 6.2.1 |
| **Strategy B**, with Transformer | $0.192 \pm 0.0127$ | $9.64 \pm 0.027$ | Section 6.2.2 |
| **Strategy C**, with Transformer | $0.185 \pm 0.0004$ | $9.52 \pm 0.021$ | Section 6.2.2 |
| PINC, Version A | $0.188 \pm 0.0005$ | $9.67 \pm 0.019$ | Section 6.3 |
| PINC, Version B | $0.190 \pm 0.0005$ | $9.77 \pm 0.010$ | Section 6.3 |

## 8 Discussion

First, Section 8.1 reviews the influence of conceptual changes, different parameter sets, and other issues encountered with the Oschmann et al. [model. In section 8.2, we discuss the different versions of the parameter inference algorithm. Last, Section 8.3 gives a broad overview of possible future steps and problems that are to be expected.

### 8.1 Model by Oschmann et al. [2017]

As part of this manuscript, we implemented the astrocytic single-compartment model originally developed by Oschmann et al. [2017] in Python. We used the model to study the dynamics of [*Ca*^2+^]_*i*_, [*Ca*^2+^]_*e*_, *h*, [*IP*_3_]_*i*_, [*Na*^+^]_*i*_, [*K*^+^]_*i*_ and *V*_*m*_ and the different mGluR and GluT driven currents using three different parameter sets (`Default`, `Paper`, `Thesis`). The observed [*Ca*^2+^]_*i*_ dynamics for parameter set `Default` resembled the Ca^2+^ dynamics seen in experimental studies [Di Castro et al., 2011, Nimmerjahn and Bergles, 2015, Verkhratsky and Nedergaard, 2018]. However, by the design of the model, they were missing the random component. In contrast to the parameter set `Default`, the parameter sets `Paper` and `Thesis` produced Ca^2+^ dynamics that resembled step functions, which is not usually seen in practice.

By studying the different currents, we observed that mGluR- and GluT-driven pathways operate on completely different orders of magnitude. The only GluT current operating on the same level as mGluR is *I*_NCX_. As the only GluT-current that directly influences the intracellular Ca^2+^ level [*Ca*^2+^]_*i*_, *I*_NCX_ is the link between the GluT- and mGLuR pathway. By operating on a completely different order of magnitude, *I*_NCX_, and therefore [*Ca*^2+^]_*i*_ are unlikely to directly influence the dynamics of [*Na*^+^]_*i*_, [*K*^+^]_*i*_, and *V*_*m*_. In part, this might be due to the used glutamate stimulation levels. While the parameters of *I*_NCX_ were tuned to match biophysical responses [Ziemens et al., 2019], the used glutamate stimulation in those experiments and in Oschmann et al. [2017] was significantly higher (1*mM*) than the glutamate stimulation used in this study (≤ 0.006*mM*) as suggested by De Pittà et al. [2009], Dupont et al. [2011], and partially Oschmann [2018]. The assumption that mGluR and GluT pathways only marginally influence each other is confirmed when considering the different conceptual changes made to the Oschmann et al. [model. Temporarily changing the leak computation to use a constant reverse potential [Farr and David, 2011, Flanagan et al., 2018] influenced the steady states of [*Na*^+^]_*i*_, [*K*^+^]_*i*_, and *V*_*m*_ significantly. All other dynamics were barely affected. It is currently assumed that cell regions close to the soma have low numbers of leak channels [McNeill et al., 2021]. Since we simulated an astrocytic compartment close to the soma, the large effect of this change on [*Na*^+^]_*i*_, [*K*^+^]_*i*_, and *V*_*m*_ was surprising.

Similarly, adding the Ca^2+^ valence to *I*_NCX_ in the computation of 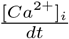 influenced [*Ca*^2+^] and [*Ca*^2+^]_*e*_ dynamics slightly, but had no effect on the other state variables. Furthermore, the removal of 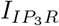 and *I*_Serca_ from the computation of 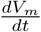 only resulted in insignificant changes in overall dynamics.

Another observation made when studying the GluT-pathway is that 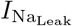 causes a positive current, implicating that the Na^+^ leak is inward pointing. This is contrary to the schematic shown in the original paper by Oschmann et al. [2017]. However, the reverse potential of Na^+^ is known to be positive [Nowak et al., 1987]. Since the membrane voltage of the astrocytic compartment was consistently negative, this behavior is likely to be correct. In summary, the astrocytic compartment model by Oschmann et al. [2017] requires further refinement to be able to accurately capture and reproduce ionic dynamics as observed in experimental data.

### 8.2 Parameter Inference

In this work, we implemented the deep learning-based parameter inference algorithm originally developed by Yazdani et al. [2020]. As a first step, we added the concepts of regularization loss, gradient clipping, and adaptive learning rates to stabilize the learning process. Second, we implemented learning rate annealing and a fully connected architecture suggested by Wang et al. [2020] to further improve and stabilize the results. Last, we added the concept of PINC and expanded the neural network to use glutamate stimulation as a control input.

The addition of gradient norm clipping to the learning process prevented the neural network from becoming unstable and predicting `NaN` values. In the literature, the problem of predicting `NaN` values is generally referred to as *exploding- or vanishing gradient problem* and is a problem more often occurring in the realms of recurrent neural networks, where gradients might become enormous due to the unrolling of network steps over several inputs [Pascanu et al., 2012]. In PINNs, these gradient pathologies also seem to be quite common and are the underlying issue Wang et al. [2020] aimed to address. Once employed, the exact gradient clipping value `c` did not significantly influence the resulting inferred parameter. However, influences in stability during the learning process and convergence speed were observed.

In another set of parameter inference experiments, we found that using an adaptive learning rate can aid or prevent PINNs from finding an appropriate solution. While the inference of *I*_NCXmax_ became unstable without a reduction in the learning rate, the same mechanism stopped the learning process of *K*_NKA*mN*_ too early. The reason for this inconsistency can probably be found in the heavy oscillations of the loss term. The employed learning rate reduction strategy, `ReducerLROnPlateau`, reduces the learning rate once the learning rate does not decrease for `patience` iterations. Due to the heavy oscillations, especially in the inference of *K*_NKA*mN*_, it is possible that exceptionally low loss outliers disturb the strategy and cause the learning rate to be decreased too early. In future work, it might be beneficial to consider registering a moving average learning rate over several epochs rather than the average learning rate of a single epoch. The general sensitivity of the parameter inference algorithm to the learning rate is unexpected. Using an adaptive learning rate mechanism such as `Adam` should lead to a low sensitivity to the used learning rate [Kingma and Ba, 2014].

The adaptive learning rate annealing strategy proposed by Wang et al. [2020] was developed for application on a single partial differential equation. We suggested three different versions to adapt their mechanism to a system of multiple ODEs and tested their application. The results showed that `Strategy A`, a strategy that aimed at weighting the loss gradient of the first ODE against all other loss gradients, was not suitable. In retrospect, this is not unexpected. The goal of the adaptive learning rate annealing is to weight ODE losses against different forms of measured data. By weighting the first ODE loss against all other ODE losses, the other ODE terms were weighted too heavily, thereby disturbing the learning process. Furthermore, the results demonstrated that `Strategy B`, weighting each ODE loss against its data counterpart, and `Strategy C`, weighting the average ODE loss against the data loss counterparts, worked equally well. Both strategies helped with stabilizing the parameter inference even without a reduction in the learning rate. To the best of my knowledge, no other adaptations of the algorithm proposed by Wang et al. [2020] to multiple ODEs exist.

Furthermore, we added Transformer networks to improve the fully connected architecture as also proposed by Wang et al. [2020]. Interestingly, this led to more heavy oscillations of the inferred parameter and the respective accuracies. While the parameter accuracy, if the average over the last 1000 epochs was taken, was higher than for the implementations without Transformer, the accuracy of inferred dynamics was lower. Transformer networks work by adding additional nodes and residual connections to the neural network. It is possible that the larger oscillations are due to the increased amount of parameters the neural network has to optimize. Wang et al. [2020] found in their paper that the addition of Transformers to their learning rate annealing mechanism reduced the end error by approximately factor three. The difference to my own experiences might be explained by the fact that they test their architecture on two-dimensional PDEs, leading them to have three input dimensions (time, x coordinate, y coordinate). In contrast, the here used ODEs consist of only one input dimension. Therefore, the addition of residual connections leading back to the input layer might not be as beneficial.

The addition of control input to the neural network, as was suggested by Antonelo et al. [2021], had several positive impacts. First, we observed that the inference of parameters is faster and more accurate than for the versions without control input. The second advantage is related to the usage of real data sets on the parameter inference algorithm. Without control input, the network would only be able to learn the data of one specific measurement series. Due to the fact that glutamate stimulation is simply assumed to be known, the network can not learn how glutamate influences the behavior of the model. Therefore, data of a possible second measurement would need to have exactly the same underlying glutamate stimulation to be useful, which might not always be achievable in practice. By adding a control input and extending it with initial conditions, multiple measurement series can now be used as long as the underlying glutamate stimulation is known. My results further indicate that learning from noisy data is possible, although it might create instabilities in the case of PINC.

Unexpectedly, runtime experiments showed that the deep learning algorithm is faster on a CPU than on a GPU. We assume that this is due to the overhead created when having to copy memory back on forth between GPU and CPU whenever the network output is fed into the computational model. Furthermore, the neural network used in this manuscript is relatively small in comparison to the size of neural networks usually used in deep learning, further increasing the significance of the overhang created by having to copy memory. By analyzing the runtime over a fixed number of epochs of different versions of my parameter inference algorithm, we found that the employed stabilization mechanisms increased the compute time. However, the increase in runtime is counteracted by the fact that the stabilized versions need fewer epochs to converge to an appropriate result.

Generally, we found that the neural network has more problems inferring parameters related to *V*_*m*_, [*Na*^+^]_*i*_, and [*K*^+^]_*i*_. This is due to several reasons. The first reason has to do with the sensitivity of the leak currents to the inferred [*Na*^+^]_*i*_ and [*K*^+^]_*i*_ values. The computation of !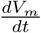 by the Oschmann et al. [model displayed disproportionately high errors in comparison to the error in the inference of [*Na*^+^]_*i*_, [*K*^+^]_*i*_ and *V*_*m*_. We attempted to solve this problem by replacing the dynamic computation of the reverse potential with constant reverse potentials, as is sometimes done in other computational models [Farr and David, 2011, Flanagan et al., 2018]. While this solution bypassed the problems of disproportional errors, it introduced a new challenge: Removing the two-way dependence between *V*_*m*_, [*Na*^+^]_*i*_ and [*K*^+^]_*i*_ seemed to make it harder for the network to appropriately infer the unobserved [*Na*^+^]_*i*_ and [*K*^+^]_*i*_. As a result, experiments attempting to infer the dynamics of [*Na*^+^]_*i*_, [*K*^+^]_*i*_, and one Na^+^-related parameter achieved lower overall 𝒜_all_ and parameter accuracy results. However, the network was more successful at learning the dynamics of the already observed data. Currently, this challenge remains open. We assume that this can be attributed to the, although faulty, stabilization process achieved by replacing the leak computation.

The second problem with inferring [*Na*^+^]_*i*_, [*K*^+^]_*i*_, and *V*_*m*_ is probably related to the behavior of the gradients over time. Usually, the gradients evaluate to almost zero, only to be extremely sharp whenever the glutamate stimulation changes. Similar problems were observed by Haghighat et al. [2021]. In their paper, they argue that PINNs do not perform well near sharp gradients as they represent highly local-rather than global behavior.

In their paper, Yazdani et al. [2020] found that they were able to infer parameters and hidden dynamics as long as they observed at least two state variables. This is in contrast to my own findings. In my experiments, the inference became unstable as soon as less than four dynamics were observed. My assumption is that this is due to the complexity of the Oschmann et al. [model. The dynamics resulting from the model are more diverse and less symmetric than the dynamics used as examples by Yazdani et al. [2020].

### 8.3 Outlook

The Oschmann et al. [model is a computational model used to simulate a single compartment of an astrocyte. Currently, it does not account for the randomness of Ca^2+^ waves or for expanded Ca^2+^ waves that are triggered by a neighboring astrocytic compartment [Di Castro et al., 2011]. Furthermore, the model assumes a constant amount of available Ca^2+^, Na^+^, and K^+^. In future work, steps could be taken to account for the phenomena of randomness and expanded Ca^2+^ transients. For example, Denizot et al. [2019] developed an astrocyte model for thin astrocytic processes that accounts for spontaneous activity. Expanded Ca^2+^ waves could be achieved by including the diffusion of Ca^2+^ between different astrocytic compartments as was originally done in the thesis of Oschmann [2018]. Steps to remove the necessity for constant amounts of Ca^2+^ were for example suggested by Taheri et al. [2017].

Several improvements are also possible in the application of PINN and PINC to the Oschmann et al. [model. These changes could focus on the stabilization of the learning process and on improving the balancing of different loss terms.

In this manuscript, we used the optimization algorithms `Adam` and `SGD`. Based on a stiffness analysis, Wang et al. [2020] suggested that using such gradient decent-based optimization strategies might not be stable. Instead, they propose further research in the use of proximal gradient algorithms [Polson et al., 2015]. Proximal gradient algorithms are an extension of classical gradient descent methods that make use of proximal operators, a mathematical, well-defined operator that poses useful properties for optimization if the minimization function *f*(*x*) is convex. Proximal gradient algorithms have for example been employed in compressive imaging [Mardani et al., 2018], in finance-related machine learning [Gu et al., 2020], or in game settings with multiple, interacting losses [Balduzzi et al., 2018]. To the best of my knowledge, no applications of proximal gradients in PINN exist so far. As Wang et al. [2020] already suggested, further work in that direction might be beneficial.

The choice of loss weights is a challenging problem encountered all over the field of parameter inference and machine learning. In the field of PINNs, it is especially challenging as the interplay between different kinds of noisy measurement data and possibly faulty differential equations has to be considered. In this manuscript, we showcased one example where a change in weights leads to enormous differences in the inference process. Furthermore, we chose an algorithm by Wang et al. [2020] to automatically compute the loss weights of the different loss terms. However, the implemented algorithm only performs well if the different terms have been approximately weighted correctly in the beginning. In future work, alternative methods should be applied. One alternative is an algorithm proposed by Xiang et al. [2021]. They suggested a method that automatically sets the loss weights based on a maximum likelihood estimation. A completely different, yet still interesting approach was developed by McClenny and Braga-Neto [2020]. Their algorithm trains multiple networks at once, attempting to learn the different weights by minimizing the total loss while maximizing the different weights.

One of the next steps is to test the algorithm developed in this manuscript on real data. However, there are multiple problems to consider. First, usually Ca^2+^ data is measured in light intensity. In contrast, the model by Oschmann et al. [2017] used in this manuscript works with ion concentrations. Measured signals can therefore not be transferred without further consideration. Second, the current implementation relies on knowledge about the used glutamate stimulation. Depending on the experiment, this information might not be readily available. Approaches not further investigated in this study might be to infer the glutamate at each time step similarly to the different parameters or to model the extracellular glutamate as an ODE, allowing inference similar to the inference of for example the *IP*_3_ concentration. The last problem is concerned with the different occurrence patterns of Ca^2+^ transients. As already mentioned earlier in this section, the Oschmann et al. [model does not account for extended Ca^2+^ waves or for randomness. This might lead to faulty inference results and future work should therefore attempt to find ways to solve this problem. Furthermore, it remains to be explored how parameter inference would work for multiple compartments.

## 9 Conclusion

Astrocytes are an important type of glial cell that are responsible for a multitude of functions in the central nervous system. However, due to their complexity and diversity, their pathways often remain unknown. To help with the general understanding of their functionality, many computational models have been developed. One of these models is the computational model of a single astrocytic compartment by Oschmann et al. [2017] that was used throughout this study. The model focuses on the separation of two pathways. The first pathway is related to the binding of glutamate through mGluR, causing the production of *IP*_3_ and associated Ca^2+^ exchanges between the ER and the cytosol. The second pathway consists of NCX, NKA, and glutamate transporters that drive the exchange of Ca^2+^, Na^+^, and K^+^ between cytosol and extracellular space. Both pathways react with changes in behavior when glutamate stimulation is present. While this model has been useful in studying the influence of the different pathways on Ca^2+^ transients, it is generally difficult to set its parameter correctly. In this work, we implemented a deep learning algorithm-based parameter inference algorithm to aid with finding these parameters.

The first version of my algorithm was an extension of the algorithm proposed by Yazdani et al. [2020]. The algorithm aims at learning the behavior of different state variables in dependence on time using PINNs. In the first step, we extended the algorithm with regularization losses, gradient clipping, and learning rate reduction to increase stability. With the resulting implementation, we were able to infer single parameters on noiseless data, as long as most dynamics were observed. However, the success of the algorithm was heavily dependent on setting exactly the correct network parameters. Otherwise, the neural network failed to converge or became unstable over time.

In the next step, we added two methods to aid with gradient pathologies as initially suggested by Wang et al. [2020]. The first method is concerned with the dynamic weighting of different loss terms. Since their paper focuses on methodologies for systems with one equation and the Oschmann et al. [model consists of seven equations, we adapted their suggestion in three different ways and tested the different strategies against each other. The most successful strategy (`Strategy C`) weighted the gradient of the sum of all ODE losses against each data loss separately. With this method, we was able to stabilize the inference of parameters without having to guess the appropriate learning rate reduction schedule perfectly. The second suggestion of Wang et al. [2020] was the addition of Transformers. In this manuscript, we did not find that their addition improves performance.

A major problem of the first two versions of my parameter inference algorithm was that the neural network only learned the time dependence but did not know about eventual changes in glutamate stimulation. Therefore, it would not have been possible to train the neural network on sets of measurement data with differing glutamate stimulation. To counteract this problem, we followed an adaptation of PINNs from control theory (PINC). The respective algorithm was proposed by Antonelo et al. [2021] and consists of splitting the data into several intervals according to changes in control input (glutamate). The neural network input was extended with nodes for the control input value and the initial conditions of the current interval. By adding the possibility of PINC to my algorithm, we sped up convergence times. In parameter inference experiments with two parameters, we showed that the PINC version achieves better results than the version without control input. However, we also observed that PINC might result in inconsistent inference results if noisy data is used.

Analysis of different currents underlying the Oschmann et al. [model revealed problems with NCX and the Na^+^- and K^+^ leaks. The influence of the leak currents on the resulting dynamics is likely too high. At the same time, NCX is barely affected by the current Na^+^ levels. Improving the dependencies within the Oschmann et al. [model might improve the stability of the parameter inference algorithm.

With the end version of my algorithm, we were able to infer parameters from artificial, noisy data.

The more dynamics were observed, the more stable the inference results became. In contrast to the original paper by Yazdani et al. [2020], we were not able to leave more than three dynamics unobserved without the results becoming unusable. Therefore, more work is necessary to make the parameter inference algorithm applicable in practice. Possible directions include further improvements in the automatic weighting of different loss terms [McClenny and Braga-Neto, 2020, Xiang et al., 2021] or the stabilization of the gradient descent function using more advanced methods than `Adam` [Polson et al., 2015].

## Notes

### Competing Interest Statement

The authors have declared no competing interest.

